# Immunometabolic state modulation of sequential decision making in patch-foraging mice

**DOI:** 10.1101/2025.09.07.674667

**Authors:** David Lau, Patrick Gagnon, Anna La Fay, Daniel Palmer, Frédérick Boisjoly, Danie Majeur, Romane Manceau, Shingo Nakajima, Josephine Robb, Yashar Zeighami, Stephanie Fulton

## Abstract

Animals have evolved sophisticated behavioural and metabolic adaptations to respond to threats to homeostasis, including resource scarcity and infectious pathogens. Energy deficits associated with lack of food availability and sickness-associated anorexia elicit distinctive hypometabolic states, however how such states are integrated with higher-order cognition is largely unknown. Patch-foraging paradigms have proven useful for deciphering evolutionarily conserved and ethologically grounded insights into cost-benefit decision-making as animals continually deliberate between exploiting and exploring their environment. We developed and extensively validated a touchscreen-based patch-foraging task for mice in which food reward dynamically varied across trials, in a dataset comprising over 111,000 sequential decisions from 35 adult male mice. Contrary to predictions that emphasize the impact of inflammation to blunt effortful reward-driven behaviour, our results demonstrate that systemic lipopolysaccharides-induced inflammation promotes hyper-exploitation by attenuating exploratory choice behaviour in animals interacting with complex food environments. Such behaviour can be seen as a bias towards immediate outcomes, with impulsivity as a feature affecting the weighting of temporal factors. Given the ubiquity of systemic inflammation in numerous infectious, metabolic and psychiatric disorders featuring dysfunctional value- and cost-sensitive behaviour, these results provide insight into how immunometabolic states are linked to altered decision-making.

## Introduction

Biological systems have evolved coordinated responses to homeostatic insults. Integrating metabolic and behavioural strategies for acquiring and expending energy in accordance with homeostatic priorities is fundamental for survival across species^1^. Two common threats to survival include resource scarcity and infectious pathogens, provoking hunger and host inflammatory responses, respectively. The associated metabolic adaptations in hunger and inflammation are characterized by partially overlapping and distinct economies of energy usage and conservation^2,3,4,5^. Both responses are associated with hypometabolism under duress of energetic deficit, with shared inhibition of costly growth and reproduction programs^6,7,8,9^. However, while starvation promotes energetically expensive foraging behaviour to acquire food^10,11^, the threat of infectious pathogens prioritizes energy conservation through lethargy in concert with the metabolic demands of immunological defense^6,12^.

The brain is a central interface that integrates both interoceptive and exteroceptive processes in organisms interacting with the environment^13^. Behavioural responses to food stimuli under conditions of hunger and inflammation indicate markedly altered reward processing and decision-making tuned to interoceptive state. Hunger potentiates appetite, food reward value, and the vigor of effortful food-directed behaviour^14,15,16,17,18^. By contrast, the constellation of behavioural responses elicited by inflammation (often termed “sickness behaviour”) includes anergia, fatigue, blunted reward motivation, and anorexia^19,20,21,22^.

Recent studies inform how interoceptive sensing of energy deprivation and inflammation influence food consummatory behaviour^23,24,25^. However, considerably less is understood about how animals make decisions during appetitive behaviour to procure food, particularly in complex food environments^26^, and recent studies have begun to interrogate the impact of unique metabolic states to shape cognition^27,28,29,30,31,32,33,34^. Behavioural patch-foraging tasks are recognized as underutilized experimental paradigms to study ethologically-inspired decisions that provide insight into cognition^35,36,37,38^. Foraging engages cost-benefit evaluation processes for making choices about how to expend energetic costs (including time and effort) in relation to the benefit of energy that could be acquired (food reward)^39,40^. Food resources are often distributed in naturalistic environments in finite “patches” that become depleted as animals consume reward. Thus, animals face the trade-off of continuing to harvest diminishing rewards from an increasingly depleted patch (exploitation) or to travel in search of a better source of reward (exploration)^41,42^. Management of this explore-exploit dilemma is thought to involve sequential comparisons of current patch reward against a global environmental average^43,44,45,46^.

To probe the impact of distinct metabolic states on cost-benefit decisions, we developed and extensively validated a touchscreen-based patch-foraging paradigm for measuring reward-sensitive choice and latency behaviour in mice. Compared to fasting conditions, acute pre-feeding transiently increased early exploratory decision-making while reducing foraging reward yield. By contrast, endotoxin-triggered acute activation of immunity strongly attenuated exploratory choice behaviour, leading to hyperexploitation. Food restriction and systemic endotoxemia elicited notable metabolic and behavioural differences. Causal linkages between metabolic state and subsequent behaviour were evident in multiple correlations between parameters reflecting energy intake/metabolism and patch foraging choice, elucidating control of cognitive decision strategy by immunometabolic state. Given the contribution of ecological immunity and immunometabolism to understanding immune theories of depression^47,48,49^, our results provide a preclinical framework in mice for studying biases in decision-making with relevance to psychiatric conditions^50,51,52^.

## Results

### Reward-sensitive behaviour in a touchscreen-based patch foraging task for mice

To assess ecological decision-making in mice, we developed a touchscreen-based patch-foraging task combining elements of existing foraging and touchscreen behavioural tasks^53,54,55,56,57,58^. Food-restricted male mice were trained to make touch responses to visual cues on a touchscreen to make a binary choice decision between either harvesting reward from a patch or to travel from their current patch to a new one (Figure 1A). The first choice option was to harvest food reward from an active patch (P_A_) by making touch responses (fixed ratio 5) to a patch cue located laterally on the touchscreen. Following a patch harvest choice, mice were required to enter the reward tray to collect a vanilla Ensure food reward. In Schedule 1, all patches began with an initial reward volume of 15µl. Sequential harvests from the same active patch decreased in volume by 1µl following each subsequent harvest (Figure 1B), to mimic declining reward rate associated with patch depletion. The alternative choice option was to “travel” to a new (non-depleted) patch by making a single touch response to an active Travel cue (T_A_) located centrally on the touchscreen, which initiated a travel cost (delay) of 5 seconds (see Methods). Following the 5 second time-out period, mice were required to complete the travel by entering the (unrewarded) tray. Following the completion of travel, a new, replenished patch opened, as represented by a new active visual cue on the opposite side of the touchscreen. In this manner, animals made choices about when to leave a patch offering increasingly smaller, but more immediate, rewards versus incurring a time delay (travel cost) to open a new, replenished patch.

**Figure 1.**
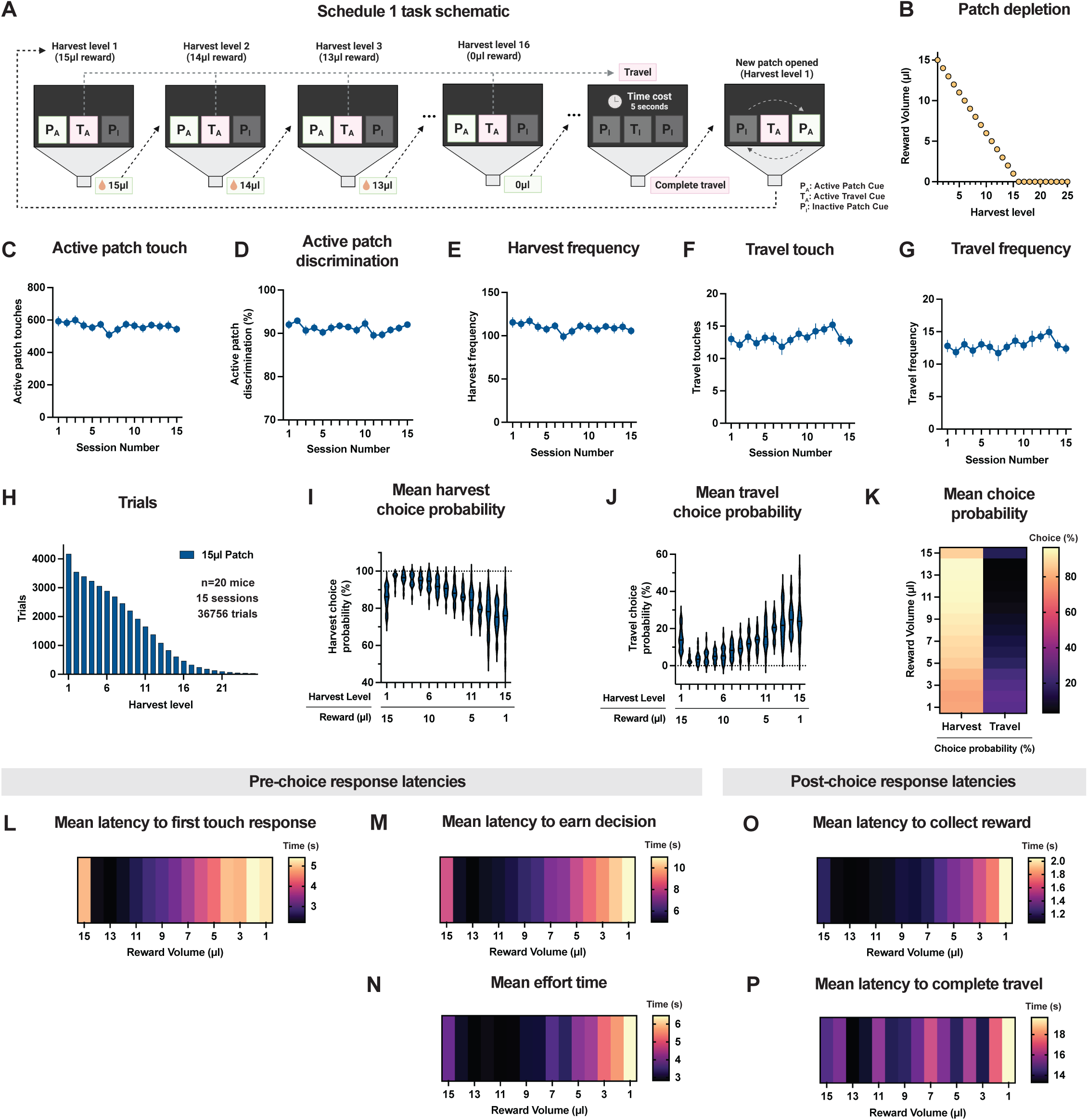
Mice perform a touchscreen-based patch-foraging task. **A.** Schedule 1 task schematic. Mice could make touch responses to either an active patch cue (P_A_) indicating the possibility to harvest a food reward from the patch or an active travel cue (T_A_) indicating the possibility to “travel” to a new (nondepleted) patch following a time cost (5 seconds). Touches to the inactive patch cue (P_I_) were recorded but did not produce any outcome. Following the completion of a “travel” a new, replenished patch would be opened. The position of the active patch alternated between the left/right sides following each subsequent travel (i.e. the position of P_A_ and P_I_ cues switched following each travel). **B.** Patch depletion during Schedule 1 task. All patches began with an initial reward volume of 15µl and depleted linearly by 1µl following each subsequent harvest. **C.** Mean active patch touch responses per session. **D.** Mean active patch discrimination per session. Active patch discrimination was calculated as the number of touch responses to the active patch cue divided by the number of total touch responses to both the active and inactive patch cues during trial decision time. **E.** Mean harvest frequency per session. **F.** Mean travel touch responses per session. **G.** Mean travel frequency per session. **H.** Histogram of total trials (sessions 1-15). **I.** Mean harvest choice probability. **J.** Mean travel choice probability. **K.** Mean harvest and travel choice probability, a summary of data shown in (**I**) and (**J**). **L.** Mean latency to first touch response. The latency to first touch response was recorded as the time difference from the start of each trial to the first touch response to an active visual cue. **M.** Latency to earn decision. The latency to earn a decision was recorded as the time difference from the start of the trial to the acquisition of the requisite touch responses to either harvest from the patch (FR5 at P_A_) or travel (FR1 at T_A_). **N.** Mean effort time. The “effort time” was calculated as the time difference between the time to earn a decision time and the latency to first touch response time. **O.** Mean latency to collect reward. The latency to collect reward was calculated as the time difference from completing a patch harvest choice and the first subsequent entry to the reward tray. **P.** Mean latency to complete travel. The latency to complete travel was calculated as the time difference between the conclusion of the 5 second time cost (when the travel completion became available), and the time for the first entry of into the reward tray. See methods for additional details on calculation of latency data. Data represented as mean ± SEM (**C-G**), histogram (**H**), mean with distribution (**I-J**), or mean (**K-P**).

Across 15 behaviour sessions in the Schedule 1 task, mice exhibited high task engagement, making hundreds of touch responses to the active patch cue per 30-minute session (Figure 1C), with high selectivity of touch responses to the active, rather than inactive, patch (Figure 1D). These active patch touch responses corresponded to on average over 100 harvests per session (Figure 1E). Food rewards harvested during the task (on average around 1ml/session, Figure S1E) represented on average above 7% of the daily calories consumed in the food restricted mice (Figure S1I). In contrast to active patch touch responding, mice made on average fewer than 15 travel touch responses (Figure 1F) and travels (Figure 1G) per session, suggesting an avoidance of the time-cost associated with travelling.

Aggregating trials across the 15 sessions from 20 mice comprised over 36,000 sequential decisions. To describe the structure of data in this task with reference to the degree of patch depletion, trials were sorted according to “harvest levels”, with “harvest level 1” representing the full, non-depleted patch, “harvest level 2” representing 1µl of depletion following one harvest from the patch, “harvest level 3” representing 2µl of depletion following 2 harvests from the patch, and so on (Figure 1H). Upon entering patches (i.e. for harvest level 2 and beyond), there was a clear sensitivity of choice behaviour to the declining patch reward (Figure 1K), as shown by a decreasing mean probability of continuing to make patch harvest choices (Figure 1I), and a corresponding increasing mean probability of travelling (Figure 1J) as the reward within patches declined. All mice tested successfully performed the task, with choice behaviour that was qualitatively similar across mice (Figure S2A-T), suggesting patch foraging tasks may be a useful alternative to avoid noted challenges associated with delay discounting tasks in mice^61,62^. Reward value was also evident in the mean latency to make touch responses, earn decisions, and collect rewards, with shorter latencies with higher reward value. The latency to harvest rewards increased as the reward within patches depleted (Figure 1L-O). By contrast, the mean latency to complete travels (an unrewarded action), was not clearly modulated by harvest level (Figure 1P).

### Across session choice behaviour learning improves reward acquisition efficiency

To examine how patch-foraging choice behaviour may change across sessions in our task, we analyzed data in blocks of 5 sessions. The mean number of rewards harvested and the travel frequency per session of each block (Figure 2A-C) and the mean total reward volume earned per session (Figure 2E-F) was unchanged across session blocks. However, the mean frequency of travels moderately increased in later sessions (Figure 2D), accompanied by an overall reduction in the mean harvest level at which mice travelled in later behavioural sessions (Figure 2G-H). This shift in choice behaviour was reflected both in changes in the mean choice probability of harvesting and travelling (Figure 2M-N), a reduction in travels in unrewarded harvest levels (Figure 2O), and changes in the distribution of travels across harvest levels (Figure 2P). Across mice, the variability in the mean harvest level at travel also decreased (Figure 2I), suggesting choice behaviour was converging to a more homogenous pattern. Although mice did not increase the overall reward obtained across sessions (Figure 2E-F), the shift in choice behaviour to tend to make travels at lower harvest levels (i.e. in states with less patch depletion) was associated with an increase in the efficiency by which mice acquired reward per effortful operant action (Figure 2K-L). In other words, by shifting their travel choice behaviour, during later behaviour sessions mice obtained more reward per unit of effort, or conversely performed less effort to obtain a given reward quantity. Mean latency data was qualitatively similar across session blocks (Figure S4A-E). Together, these results suggest that with increasing experience with the patch-foraging task, mice learned to improve their choice behaviour to more efficiently utilize energy to acquire reward.

**Figure 2.**
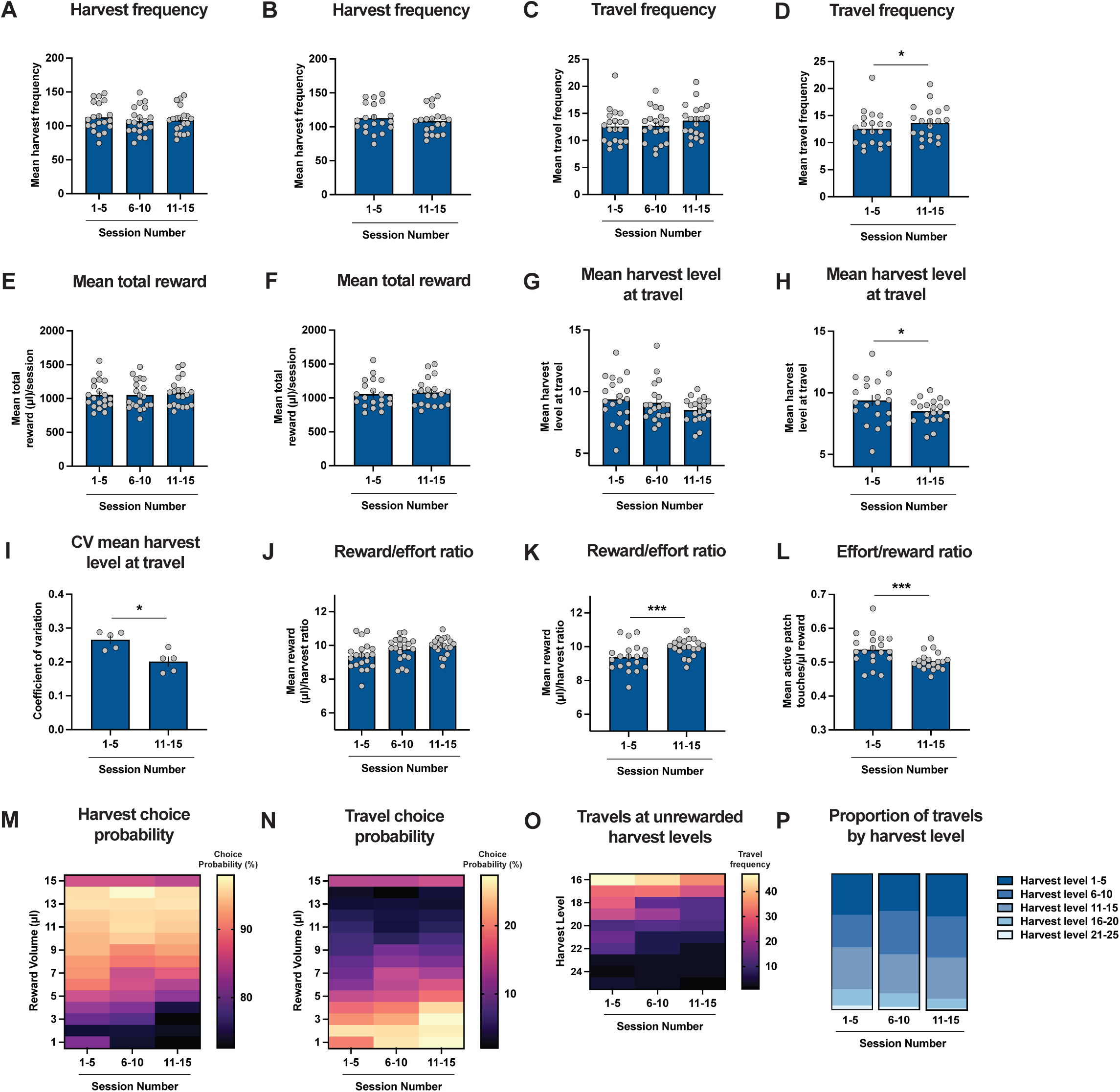
Mice improve their choice behaviour across sessions to more efficiently acquire reward. **A.** Harvest frequency by session block. **B.** Harvest frequency. **C.** Travel frequency by session block. **D.** Travel frequency. **E.** Mean total reward earned per session by session block. **F.** Mean total reward. **G.** Mean harvest level at travel by session block. **H.** Mean harvest level at travel. **I.** Coefficient of variation of mean harvest level at travel. **J.** Reward/effort ratio by session block. **K.** Reward/effort ratio. **L.** Effort/reward ratio by session. **M.** Mean harvest choice probability. **N.** Mean travel choice probability. **O.** Frequency of travels at unrewarded harvest levels by session block. **P.** Proportion of travels across harvest levels by session block. Data represented as mean ± SEM (**A-L**) or mean (**M-O**); *p<0.05, ***p<0.001.

### Acute satiety transiently promotes early exploratory choice behaviour

Changes in energy status are well appreciated to impact numerous facets of behaviour^16,63,64^. To begin to explore how acute variation in metabolic state may influence patch-foraging choice behaviour in mice, we tested the impact of pre-feeding food restricted mice for 1 hour prior to behaviour sessions compared to baseline performance under regular food-restriction (Figure 3A). Metabolic phenotyping data demonstrate 1 hour of pre-feeding in fasted mice is sufficient to significantly alter metabolic state (Figure S15E-J). While body weight 1hr prior to behaviour was relatively well controlled across sessions (Figure 3B), mice consumed on average over 2 kcal of food during pre-feed periods preceding behaviour sessions (Figure 3C). These differences in food consumption were associated with corresponding changes in body weight (Figure 3D-F), with mice having higher body weight before beginning pre-fed behaviour sessions (Figure 3G). Likely reflecting the heterogeneity in food consumed during the pre-feed period, ranging from under 2 kcal to over 6 kcal (Figure 3C), body weight variability was higher following pre-feeding compared to baseline (Figure 3H).

**Figure 3.**
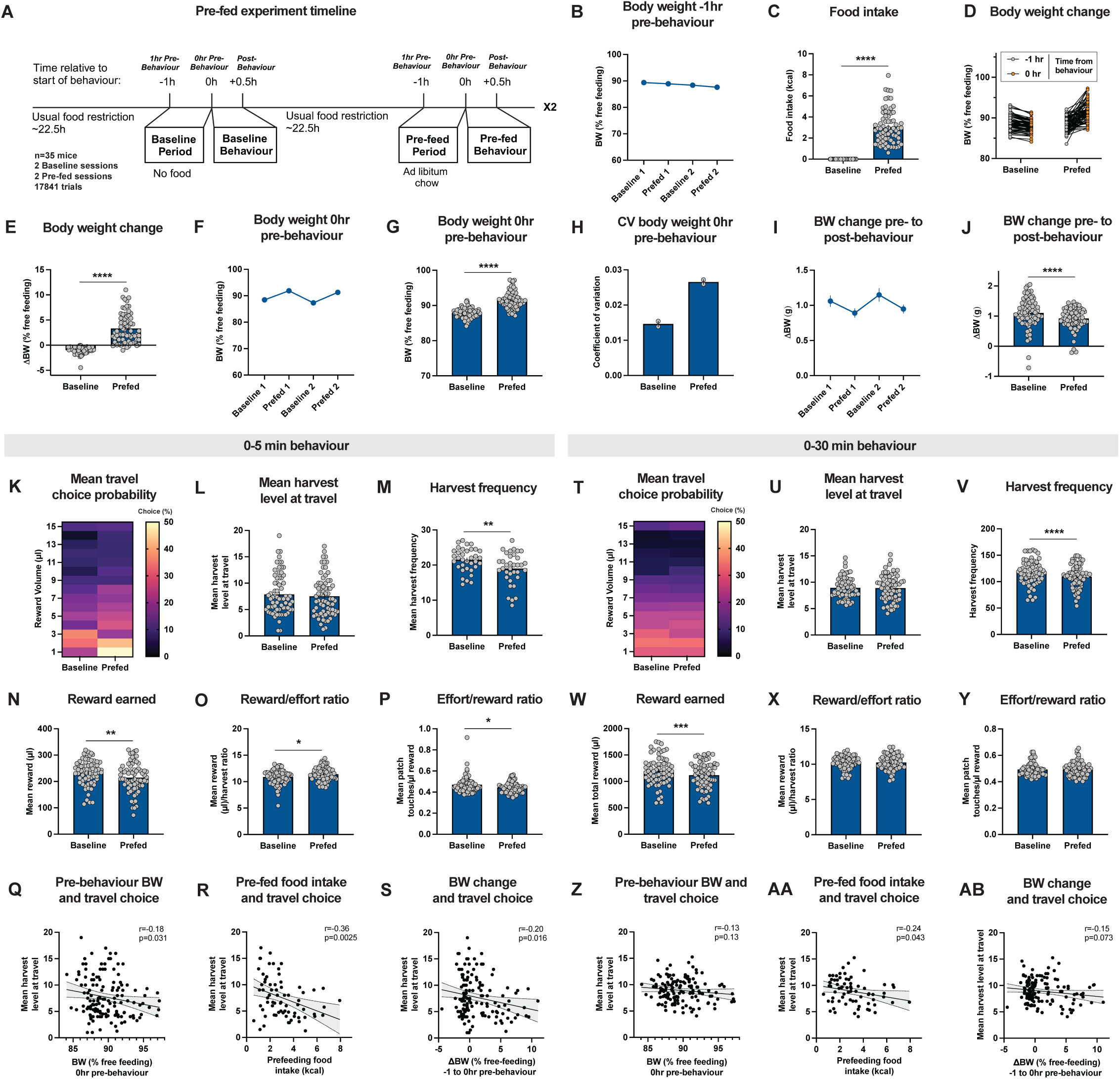
Acute variation in metabolic state controls early exploratory choice. **A.** Schematic of pre-fed experiment timeline. **B.** Relative body weight (% free-feeding body mass) 1hr pre-behaviour. **C.** Food intake during baseline period and pre-feed period. **D.** Relative body weight (% free-feeding body mass) at start and end of baseline period and pre-feed period. **E.** Change in relative body weight shown in (**D**). **F.** Relative body weight (% free-feeding body mass) 0hr pre-behaviour. **G.** Relative body weight shown in (**E**). **H.** Coefficient of variation of session relative body weight shown in (**G**). **I.** Change in absolute body weight from 0hr pre-behaviour to post-behaviour period. **J.** Change in absolute body weight shown in (**I**). **K.** Mean travel choice probability during (0-5 minutes behaviour). **L.** Mean harvest level at travel (0-5 minutes behaviour). **M.** Mean harvest frequency (0-5 minutes behaviour). **N.** Reward earned (0-5 minutes behaviour). **O.** Reward/effort ratio (0-5 minutes behaviour). **P.** Effort/reward ratio (0-5 minutes behaviour). **Q.** Relative body weight (% free-feeding) at 0hr pre-behaviour and mean harvest level at travel (0-5 minutes behaviour). **R.** Pre-feeding food intake (kcal) and mean harvest level at travel (0-5 minutes; pre-fed behaviour). **S.** Change relative body weight (% free-feeding) during baseline period and pre-feed period and mean harvest level at travel (0-5 minutes behaviour). **T.** Mean travel choice probability (0-30 minutes behaviour). **U.** Mean harvest level at travel (0-30 minutes behaviour). **V.** Harvest frequency (0-30 minutes behaviour). **W.** Reward earned (0-30 minutes behaviour). **X.** Reward/effort ratio (0-30 minutes behaviour). **Y.** Effort/reward ratio (0-30 minutes behaviour). **Z.** Relative body weight (% free-feeding) at 0hr pre-behaviour and mean harvest level at travel (0-30 minutes behaviour). **R.** Pre-feeding food intake (kcal) and mean harvest level at travel (0-30 minutes; pre-fed behaviour). **S.** Change relative body weight (% free-feeding) during baseline period and pre-feed period and mean harvest level at travel (0-30 minutes behaviour). Data represented as mean ± SEM (**B, C, E-J, L-P, U-Y**), mean (**K, T**), or individual points (**Q-S, Z-AB**); *p<0.05, **p<0.01, ***p<0.001, ****p<0.0001. Black lines and shaded areas represent models and 95% confidence intervals, respectively, from simple linear regression (**Q-S, Z-AB**). Paired two-tailed t-test (**C, E, G, J, L-P, U-Y**), and output of two-tailed Pearson’s correlation shown in top right of (**Q-S, Z-AB**).

We began by examining overall choice behaviour from the entire 30-minute behaviour sessions. Mean travel choice behaviour and mean harvest level at travel (Figure 3T-U) and travel frequency (Figure S5O-P) were unchanged between baseline and pre-fed sessions. However, mice in baseline sessions had significantly elevated harvest frequency (Figure 3V), with correspondingly increased total reward earned (Figure 3W). Across Schedule 1 behaviour sessions, variation in total session reward strongly correlated with changes in body weight during behaviour sessions (Figure S1G). These results suggest mice adjusted their behavioural responding in accordance with the degree of energetic deficit, resulting in compensation for the additional body weight lost during baseline sessions corresponding to a significant increase in the gain of body weight during baseline behaviour sessions (Figure 3I-J). As mice were pre-fed chow, these differences reflect differences in metabolic state rather than sensory-specific satiety^65^. Consistent with the augmented harvest (Figure 3V) and trial frequency (Figure S5Q-R) in baseline sessions, faster response latencies were observed in baseline sessions (Figure S6A-E). In line with choice behaviour (Figure 3T-U), the reward/effort ratio (total Ensure earned/total active patch touch responses) was unchanged from the overall behaviour period (Figure 3X).

We considered two factors which may occlude differences in metabolic state to impact decision making between baseline and pre-fed sessions. First, consistent with prior literature^66^, animals made more exploratory choices early during behaviour sessions, allowing for greater potential to observe variance in exploratory choice early in behaviour sessions. Additionally, within-session learning may produce more homogenous behavioural strategies as sessions progress. Second, within-session changes in metabolic state as animals consume food reward may alter ongoing decision processes (Figure S1L, Figure 3I-J), and we reasoned that differences in metabolic state to influence decision choice may be most apparent during the initial behaviour period. This possibility was particularly pertinent since mice increased their body weight to a greater extent in baseline sessions, suggesting convergence of metabolic states late in behaviour sessions. We thus focused analysis on trials from the first 5 minutes of behaviour. Harvest frequency and rewards earned were elevated in baseline sessions (Figure 3M-N), consistent with the entire session. However, the reward/effort ratio and effort/reward ratio were decreased and increased, respectively, in baseline sessions (Figure 3O-P), suggesting differences in exploratory choice in addition to higher response rate during baseline sessions. Consistent with this, mean travel choice probability noticeably shifted to higher harvest levels (lower reward volumes) in baseline sessions (Figure 3K). Additionally, since there was notable variation in pre-feeding food intake and body weight prior to pre-fed sessions, we analyzed whether variation in body weight during the period prior to behaviour was related to subsequent choice in the following first 5 minutes of behaviour. Notably, a significant negative correlation was observed (Figure 3Q), suggesting mice with greater satiety were more likely to leave the patch to explore. Implying this relationship was related to energy deprivation state, no correlation was observed when comparing to absolute body weight (Figure S7C). Additionally, this relationship was driven by acute changes in metabolic state as this correlation was not observed when analyzing body weight 1 hr prior to behaviour (prior to the baseline and pre-feed period; Figure S7A). Additional correlations also linked other features related to metabolic state, including food consumption during pre-feeding sessions (Figure 3R), and change in body weight prior to behaviour (Figure 3S) with exploratory choice. Similar correlation analyses demonstrated that metabolic state-related body weight or feeding did not explain, or only more weakly explained, variation in overall choice behaviour from the 30-minute behaviour period (Figure 3Z-AB). By contrast, moderate correlations were observed between the absolute (not relative) body weight prior to sessions and choice behaviour during the 30-minute period (Figure S7F-G).

### Behavioural reward-sensitivity is conserved within and across patches of varying initial reward magnitude

To extend our behaviour task, we developed a Schedule 2 version in which three patch types were introduced (Figure 4A). Following the completion of a travel, mice had ∼33% chance of entering a patch with either 6µl (poor), 12µl (medium), or 18µl (rich) initial reward volume, with uniform linear patch depletion rates of −1µl per harvest (Figure 4B). Nineteen mice were run on the Schedule 2 behaviour task. Over 18 behaviour sessions comprising over 51,000 trials (Figure 1C), mice exhibited high task engagement, performing >100 harvests on average (Figure 4D-E) and upwards of 20 travels per 40-minute behaviour session (Figure 4F-G). The majority of harvests and rewards earned were attributable to the 18µl patch (Figure 4E, 4H-I). Consistent with this, harvest and travel choice behaviour and the mean harvest level at travelling were highly discriminable between patch types (Figure 4J-K, 4N). Congruently, when examining trials with reference to reward within different patch types (rather than harvest level), sensitivity to reward magnitude was observed following entry into patches (Figure 4L-M), indicating patch-leaving choice behaviour within and across patch types was strongly linked to reward. Consistent with these findings, sensitivity to reward magnitude within and across patches was additionally observed in mean pre-choice latency responses (Figure 4O-Q) and the mean latency to collect reward (Figure 4R). Collectively, these results demonstrate robust within- and across-patch reward sensitivity in the Schedule 2 task.

**Figure 4.**
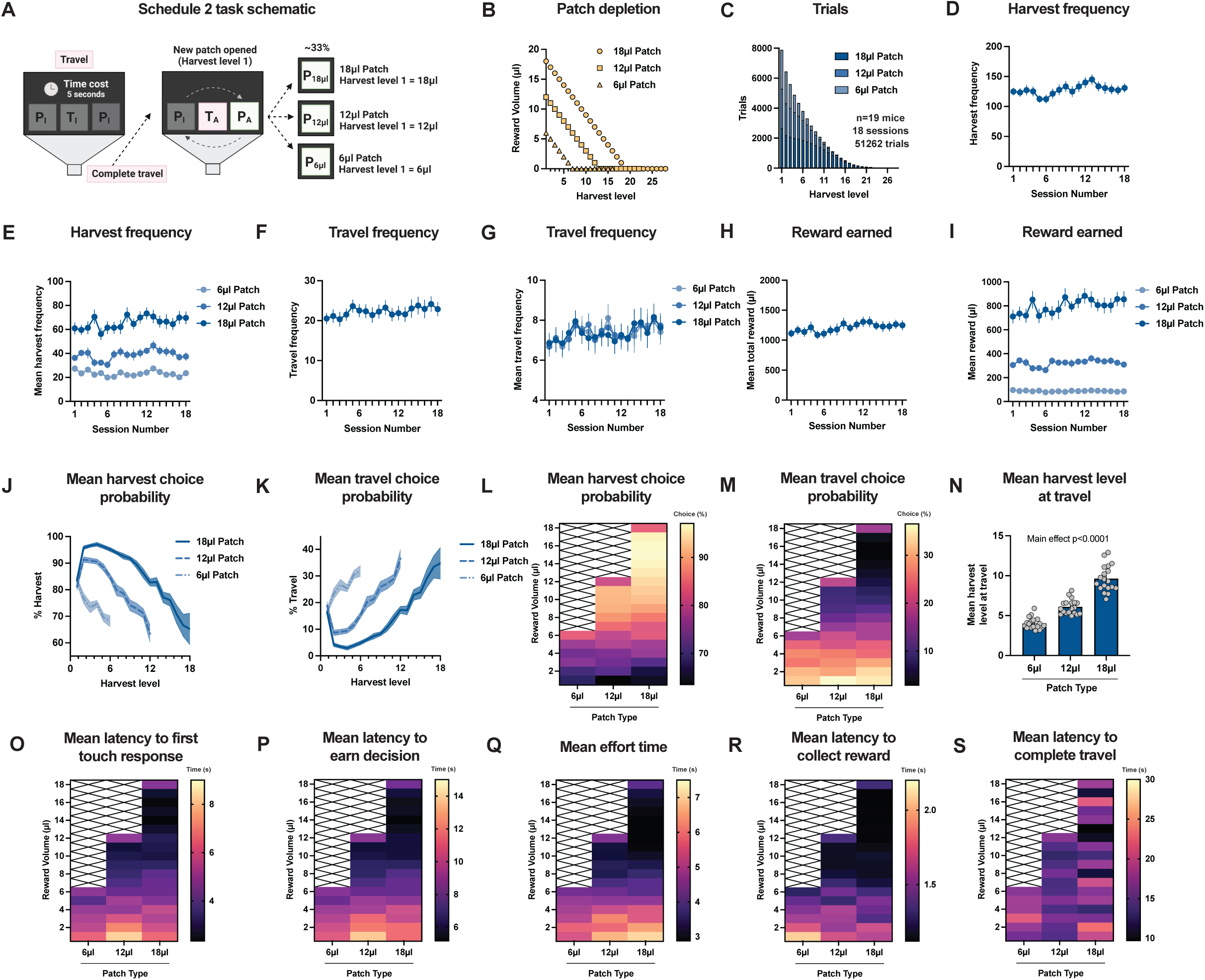
Behavioural reward sensitivity is conserved across patches of varying initial reward. **A.** Schematic of Schedule 2 task. In contrast to Schedule 1 in which all patch types began with an initial 15µl reward volume, three patch types were introduced with initial reward volumes of 6µl, 12µl, or 18µl. Mice had approximately 33% chance of encountering each of the 3 patch types following the completion of a travel. **B.** Patch depletion during Schedule 2 task. While initial reward volumes varied by patch type (6µl, 12µl, or 18µl), all patches depleted linearly at the same rate of 1µl following each subsequent harvest in the respective patch. **C.** Histogram of total trials (session 1-18). **D.** Mean harvest frequency by session. **E.** Mean harvest frequency by session, separated by patch type. **F.** Mean travel frequency by session. **G.** Mean travel frequency by session, separated by patch type. **H.** Mean total reward earned per session. **I.** Mean total reward earned per session, separated by patch type. **J.** Mean harvest choice probability. **K.** Mean travel choice probability. **L.** Mean harvest choice probability, summary of data shown in (**J**). **M.** Mean travel choice probability, summary of data shown in (**K**). **N.** Mean harvest level at travel. **O.** Mean latency to first touch response. **P.** Mean latency to earn decision. **Q.** Mean effort time. **R.** Mean latency to collect reward. **S.** Mean latency to complete travel. Data represented as histogram (**C**), mean ± SEM (**D-K, N**) or mean (**L, M, O-S**). Hatched areas indicate no data. One-way ANOVA, main effect of patch type (**N**).

### Distinct behavioural and metabolic states provoked by hunger, re-feeding, and systemic LPS-induced immune activation

To determine the metabolic consequences of food restriction and systemic inflammation, we subjected mice to metabolic phenotyping using the Comprehensive Lab Animal Monitoring System (CLAMS), in a manner designed to emulate the corresponding behavioural experiment (Figure 6). Acute systemic inflammation was induced via lipopolysaccharides (LPS), a potent activator of innate immunity and modulator of metabolism via TLR4 activation^12^. Food-restricted mice (Figure 5B-C) were introduced into the CLAMS with food availability restricted to 6 hours/day beginning during the dark cycle (∼ZT16), for three experimental days. On the first experimental day, mice were intraperitoneally injected with saline. Food intake was thus structured during the food restriction period (post-saline days 1-3) according to periods of food availability (Figure 5D-E)^67^. When comparing 1hr transition periods from fasting to re-feeding during this period (Figure S15A-B), expected differences were observed in energy expenditure (Figure S15E-F), respiratory exchange ratio (Figure S15G-H), and fatty acid oxidation (Figure S15I-J). These results confirm that 1 hour of chow-refeeding in chronically food-restricted mice is sufficient to acutely alter metabolic state compared to fasting (Figure 3). Mice gradually increased their food intake over the three food availability periods (post-saline days 1-3), suggesting habituation to the feeder and food availability regime in the CLAMS (Figure S16A-B). As food intake on post-saline day 3 was more representative of feeding during a related behaviour experiment (Figure 6B), we focused our comparisons to this time period (however, similar results were obtained when comparing analyses with respect to injection time; Figure S17A-N). At the beginning of the fourth experimental day, mice were intraperitoneally injected with 0.33mg/kg LPS, with *ad libitum* food available for the remainder of the time in the CLAMS (Figure 5A). A separate experiment confirmed this dose of LPS was sufficient to robustly increase mRNA expression of pro-inflammatory cytokines *Tnf* (tumor necrosis factor) and *Il1b* (interleukin-1 β), and chemokine *Ccl2* in the arcuate nucleus (ARC), ventral tegmental area (VTA), and nucleus accumbens (NAc) (Figure S18A-P).

**Figure 5.**
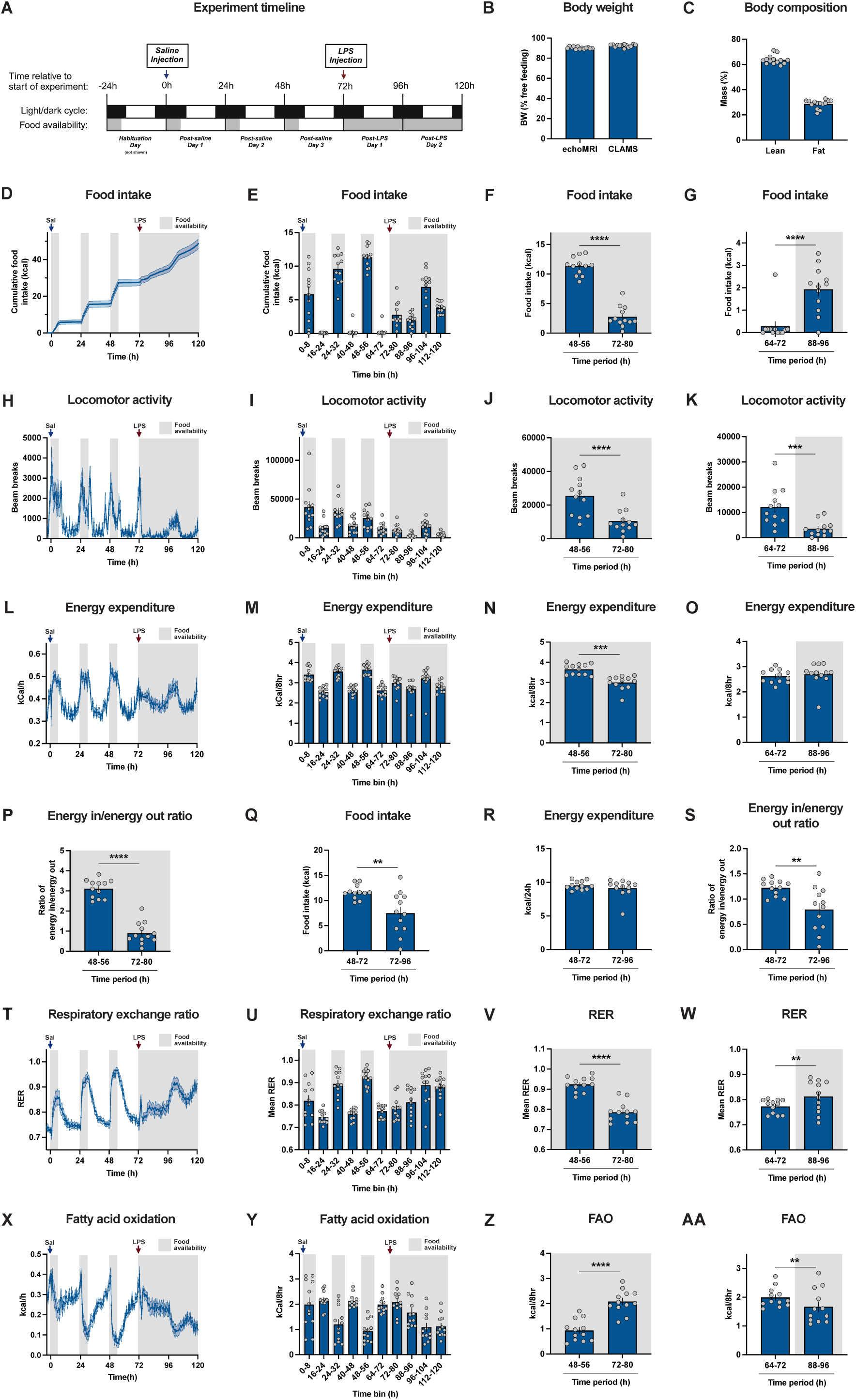
Distinctive behavioural and metabolic states are induced by hunger, re-feeding, and LPS-induced inflammation. **A.** Schematic of CLAMS experiment timeline. During the first 72h of the experiment, mice had access to food within the CLAMS for 6h each day, beginning 4-5h into the dark cycle (ZT 16-17). Mice were injected (i.p.) with saline at 0h (start of experiment) and LPS (0.33mg/kg) at 72h. Mice had ad libitum access to food following LPS injection. In light/dark cycle timeline, black indicates dark cycle, and white indicates light cycle. In food availability timeline, grey indicates food availability within CLAMS, and white indicates lack of food availability within CLAMS. **B.** Relative body weight during echoMRI measurement and prior to CLAMS experiment. **C.** Lean and fat mass. **D.** Cumulative food intake. **E.** Summary of food intake in (**D**). **F.** Food intake during feeding period of post-saline day 3 (48-56h) and the corresponding period following LPS injection (72-80h). **G.** Food intake during the end period of post-saline day 3 (64-72h) and the corresponding period of post-LPS day 1 (88-96h). **H.** Mean locomotor activity. **I.** Summary of locomotor activity in (**H**). **J.** Locomotor activity during feeding period of post-saline day 3 (48-56h) and the corresponding period following LPS injection (72-80h). **K.** Locomotor activity during the end period of post-saline day 3 (64-72h) and the corresponding period of post-LPS day 1 (88-96h). **L.** Mean energy expenditure. **M.** Summary of energy expenditure in (**L**). **N.** Energy expenditure during feeding period of post-saline day 3 (48-56h) and the corresponding period following LPS injection (72-80h). **O.** Energy expenditure during end period of post-saline day 3 (64-72h) and the corresponding period of post-LPS day 1 (88-96h). **P.** Ratio of energy in/energy out, calculated as the ratio of food intake (kcal; shown in **F**) divided by energy expenditure (kcal; shown in **N**) during the feeding period of post-saline day 3 (48-56h) and the corresponding period following LPS injection (72-80h). **Q.** Total daily food intake during post-saline day 3 (48-72h) and post-LPS day 1 (72-96h). **R.** Total energy expenditure during post-saline day 3 (48-72h) and post-LPS day 1 (72-96h). **S.** Energy in/energy out ratio during post-saline day 3 (48-72h) and post-LPS day 1 (72-96h). **T.** Mean respiratory exchange ratio (RER). **U.** Summary of RER in (**T**). **V.** Mean RER during feeding period of post-saline day 3 (48-72h) and the corresponding period following LPS injection (72-80h). **W.** Mean RER during the end period of post-saline day 3 (64-72h) and the corresponding period of post-LPS day 1 (88-96h). **X.** Mean fatty acid oxidation. **Y.** Summary of fatty acid oxidation in (**X**). **Z.** Fatty acid oxidation during feeding period of post-saline day 3 (48-72h) and the corresponding period following LPS injection (72-80h). **AA.** Fatty acid oxidation during end period of post-saline day 3 (64-72h) and the corresponding period of post-LPS day 1 (88-96h). Data represented as mean ± SEM (**B-AA**); **p<0.01, ***p<0.001, ****p<0.0001. Paired two-tailed t-test (**F, G, J, K, N-S, V, W, Z, AA**).

**Figure 6.**
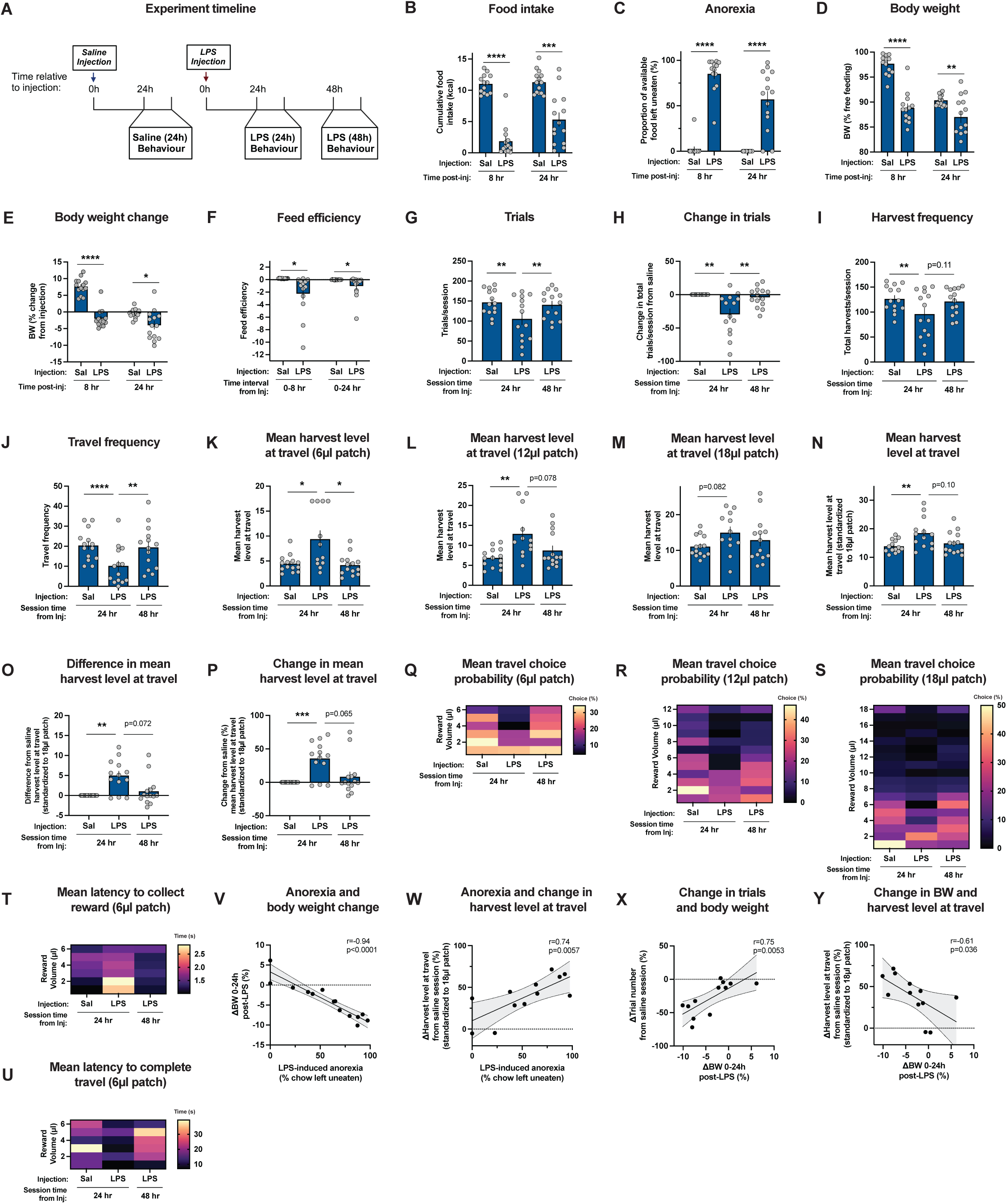
LPS-induced inflammation strongly inhibits exploratory choice during patch-foraging. **A.** Schematic of behavioural experiment timeline. Mice were injected with saline or LPS (0.33mg/kg) and performed Schedule 2 behavioural patch-foraging task at 24h (saline and LPS) and 48h (LPS) following injection. **B.** Cumulative food intake period 8h (left) and 24h (right) following injection. **C.** Proportion of available food left uneaten at 8h (left) and 24h (right) following injection. **D.** Body weight at 8h (left) and 24h (right) following injection. **E.** Body weight change from injection time to 8h (left) and 24h (right) following injection. **F.** Feed efficiency during 0-8h (left) and 0-24h (right) period following injection. **G.** Trials per session. **H.** Change in total trials per session. **I.** Harvest frequency per session. **J.** Travel frequency per session. **K.** Mean harvest level at travel (6µl patch). **L.** Mean harvest level at travel (12µl patch). **M.** Mean harvest level at travel (18µl patch). **N.** Mean harvest level at travel (all trials standardized to 18µl patch reward). **O.** Difference in mean harvest level at travel (all trials standardized to 18µl patch reward). **P.** Change in mean harvest level at travel (all trials standardized to 18µl patch reward). **Q.** Mean travel choice probability (6µl patch). **R.** Mean travel choice probability (12µl patch). **S.** Mean travel choice probability (18µl patch). **T.** Mean latency to collect reward (6µl patch). **U.** Mean latency to complete travel (6µl patch). **V.** LPS-induced anorexia (chow left uneaten, shown in right **C**) and change in body weight from injection time to 24h following injection (shown in right **E**). **W.** Anorexia 24h following LPS injection (shown in right **C**) and change in mean harvest level at travel during LPS (24h) behaviour session relative to saline (24h) behaviour session (shown in **P**). **X.** Change in body weight 24h following LPS injection (shown in right **E**) and change in trial frequency during LPS (24h) behaviour session (shown in **H**). **Y.** Change in body weight 24h following LPS injection (shown in right **E**) and change in mean harvest level at travel during LPS (24h) behaviour session relative to saline (24h) behaviour session (shown in **P**). Data represented as mean ± SEM (**B-P**), mean (**Q-U**), or individual points (**V-Y**); *p<0.05, **p<0.01, ***p<0.001, ****p<0.0001. Black lines and shaded areas represent models and 95% confidence intervals, respectively, from simple linear regression (**V-Y**). Paired two-tailed t-test (**B-F**), Friedman test with Dunn’s multiple comparisons (**G-I**), one-way ANOVA with Geisser-Greenhouse correction and Tukey’s multiple comparisons (**J**), mixed effects model (REML) with Geisser-Greenhouse correction and Tukey’s multiple comparisons (**K-P**), and output of two-tailed Pearson’s correlation shown in top right (**V-Y**).

During the first 8 hours following injection of LPS, food intake was markedly decreased (Figure 5E-F), consistent with sickness-induced anorexia. Reductions in energy intake were attributable to reduced meal frequency (Figure S16C-E), rather than changes in meal size (Figure S16F-H). By contrast, food intake was elevated during the 16-24 hour period following LPS injection (Figure 5G), approximating feeding patterns during a related behaviour experiment (Figure 6B). Consistent with sickness behaviours of lethargy and anergia, locomotor activity was strongly suppressed following LPS injection (Figure 5H-K), an effect which may be driven by systemic glucose metabolism^68^. Periods of increased energy expenditure were linked to feeding during food-restriction (post-saline days 1-3; Figure 5L-M), in line with food-entrainable clock synchronization of metabolic cycles during chronic daily food restriction^69,70^ and the thermic effect of food. Food intake and energy expenditure data suggest no alterations in circadian cycle occurred throughout the experiment (Figure S16C, Figure 5L-M). In the first 8 hours following LPS injection, energy expenditure was reduced (Figure 5N), likely due to hypolocomotion, reduced digestion, and hypothermia^12^. The ratio of energy ingested to energy expended during this period was correspondingly altered by LPS injection (Figure 5P), consistent with the utilization of stored energetic fuels in activation of immunity^12,71^. When examining the entire 24-hour period following LPS injection, food intake remained suppressed (Figure 5Q). However, 24-hour energy expenditure was unchanged (Figure 5R), demonstrating similar energetic usage in the hypometabolic states of hunger and acute inflammation characterized in our protocol. Such results emphasize dramatic restructuring of metabolic prioritization of energy use in food restriction compared to LPS-induced sickness. While LPS-triggered inflammation initiated avoidance of energetic costs of locomotor activity and digestion, similar overall energy expenditure compared to food restriction was likely accounted for by the metabolic expense of host defense.

Consistent with changes in energy intake and usage, substrate utilization as measured by the respiratory exchange ratio was altered by fasting, re-feeding, and LPS-induced suppression of feeding (Figure 5T-U). Increased carbohydrate utilization corresponded to periods of feeding during food restriction (Figure 5T, Figure S15G-H). Consistent with altered feeding patterns following LPS injection, RER was reduced in the first 8 hr following LPS injection (Figure 5V), while increased in the period 16-24 hr following LPS injection (Figure 5W). Relatedly, fatty acid oxidation was elevated during the first 8 hrs following LPS injection (Figure 5Z) and reduced 16-24 hr following LPS injection (Figure 5AA). Collectively, these results demonstrate food restriction and LPS-induced inflammation elicit distinctive behavioural and metabolic adaptations, underscoring dissociable physiological responses to homeostatic threats of resource scarcity and pathogens.

### LPS-induced sickness promotes hyper-exploitation and blunted reward valuation during patch-foraging

To evaluate the impact of distinct immunometabolic states on sequential decision-making, fourteen food-restricted mice (Figure S19A-B) were within-subject tested 24 hours following intraperitoneal injections of saline or LPS in the Schedule 2 patch-foraging task (Figure 6A). To study recovery from inflammation, mice were additionally tested 48 hours following LPS injection. Mice consumed all or a majority of available food during the regular food restriction in the 8 hours following saline injection, with 13/14 mice consuming all food (Figure 6C). By contrast, food consumption was attenuated during the first 8 hours following LPS injection (Figure 6B-C), consistent with CLAMS data (Figure 5) and confirming that an inflammatory response was provoked in all mice. While the majority of saline injected mice were completely fasted for the 16-hour period from 8-24 hours post injection, cumulative food intake continued to modestly increase in the corresponding 16-hour period following LPS injection (Figure 6B). In accordance with divergent feeding patterns, body weight was lowered at 8- and 24-hours following LPS injection (Figure 6D-E). Consistent with lowered energy intake/expenditure ratios observed in the CLAMS (Figure 5P, 5S), feed efficiency was also reduced following LPS injection (Figure 6F).

Control behavioural data 24 hr following saline injection was comparable to a previous cohort of mice (Figure 4) with reward-sensitivity observed within and across patches in travel choice and pre-choice response latency behaviour (Figure S19K, S19N, Figure S20A, S20D, S20G). When examining patch-foraging behaviour 24 hours following LPS injection, the number of trials performed during the 40-minute behaviour sessions were reduced compared to 24 hours following saline injection (Figure 6G-H), with associated reductions in both harvest and travel frequency (Figure 6I-J). However, numerous metrics indicate this reduction was not explained by memory loss of the task, nonspecific responding, or impaired task performance generally. Active patch discrimination was not altered (Figure S19J), significant discrimination of patch type was evident in travel choice behaviour (Figure S19K-M) and latency to travel choice (Figure S19Q-S), and reward-sensitivity was broadly present within and across patches in patch-leaving choice behaviour (Figure S19N-P) and pre-choice response latencies (Figure S20A-I) in all behaviour sessions. Additionally, all mice still consumed all earned Ensure during the task despite marked appetite loss for chow following LPS injection. Collectively, these data strongly suggest that the structure of task behaviour remained reward-guided during sickness, allowing for direct comparison across sessions. As a strong indication of altered choice behaviour, the mean harvest level at travelling at 24 hours following LPS injection was significantly elevated from 6µl and 12µl patches, with a similar trend observed in 18µl patches (Figure 6K-M). Mean travel choice probability patterns within and across patches were also altered (Figure 6Q-S). When standardizing trials across patch types to the reward offered (Figure 6N), the mean harvest level at travelling was elevated in 10/13 mice, and recovered close to saline control levels following 48 hours in a majority of the mice (Figure 6O-P). Together, these results suggest that overharvesting and increased bias for immediate rewards was induced by acute systemic inflammation.

As approach and reward collection latencies scale with reward and are often interpreted as reflecting reward value representations^72^, it is plausible that changes in reward-oriented behaviours may reflect reward anhedonia associated with sickness and/or sickness-related lethargy or anergia. Pre-choice response latencies to the first touch response were slower in the 18µl patch (Figure S21A), and reward collection latencies were similarly elevated across patches (Figure S21J-K, 6T). Suggesting these effects were not driven by motor impairment, mean effort time latencies (a proxy for operant response rate) were similar within the 18µl patch (Figure S21G) and appeared only modestly increased in the 6µl and 12µl patches (Figure S21H-I). By contrast, the mean latencies for mice to complete travels exiting from all patch types were reduced, rather than increased at 24 hours following LPS injection (Figure S21L-M, 6U). These results suggest that acute inflammation may have facilitated more rapid transitions from states of exploration to exploitation despite slower exploitation-oriented reward-acquisition and reward-collection behaviour. These findings are consistent with the notion that changes in response latency were not solely attributable to motor slowing following LPS injection.

Notable heterogeneity was observed in the magnitude of LPS-induced anorexia at 24 hours following LPS injection (Figure 6C), changes in total number of completed trials (Figure 6H), and mean travel choice behaviour (Figure 6P). We speculate this may be related to underlying variation in immune activation and/or recovery across mice. Variation in LPS-induced anorexia was strongly correlated with change in body weight 24 hours following LPS injection (Figure 6V). Suggesting that the degree of sickness explains variation in choice behaviour, LPS-induced anorexia positively correlated with changes in travel choice behaviour (Figure 6W), indicating that mice with the greatest loss of appetite had the largest shift towards exploitation. Change in body weight following LPS injection was additionally positively correlated with changes in trial frequency (Figure 6X) and negatively correlated with changes in mean travel choice behaviour (Figure 6Y). Together, these results demonstrate that behavioural and metabolic changes associated with suppressed feeding and loss of body weight affect sequential choice decision-making during patch-foraging.

## Discussion

Measures of ethological decision-making can offer valuable insights into how the brain processes value under different metabolic conditions during interactions with complex food environments. Over recent decades, changes in the food environment are theorized to have propelled increases in obesity and metabolic disease, with associated chronic low-grade inflammation. However, the impacts of immunometabolic states to influence naturalistic food-based choices remain incompletely understood. We developed a touchscreen-based patch-foraging paradigm for mice with reward-sensitive choice and latency behaviour within and across patches of varying initial reward. Our results illustrate divergent impacts of varying immunometabolic states on sequential decision-making during patch-foraging. In comparison to standard food-restriction, acute chow pre-feeding transiently promoted exploratory choice early within behaviour sessions. By contrast, LPS-induced sickness, a categorically different appetite-suppressing manipulation from pre-feeding, durably attenuated exploratory choice throughout behaviour sessions, resulting in an unexpected hyper-exploitative choice strategy. These results are in agreement with a recent study on IL-6 signalling in cancer cachexia^73^, suggesting behavioural responses may be shared across inflammatory challenges. Both pre-feeding and LPS-injection promoted motor slowing, as suggested by increased pre-choice reaction latencies, but were associated with oppositional changes in body weight and markedly distinct impacts on exploratory choice behaviour. Food restriction and LPS-injection both were associated with reduced body weight compared to pre-feeding, but exerted opposing impacts on response latencies; while food-restriction invigorated faster foraging behaviours, LPS induced motor slowing in pre-choice latencies. Metabolic phenotyping revealed that food-restriction and LPS-induced sickness were both characterized by similar overall metabolic expenditure under our experimental conditions, but distinctive patterns of energy usage reflecting divergent metabolic priorities under conditions of hunger and inflammation. Strongly suggesting a direct linkage between metabolic state and subsequent behavioural choice, robust correlations were observed between parameters of appetite/metabolism and patch-foraging choice behaviour.

Anorexic behaviour is a highly conserved response to infection that may be adaptive for survival^19^, despite the considerable energetic burden of immunological defense^6^. While a net anabolic response may be required to mount pathogen resistance responses^2^, catabolic states may be important for eliciting host tolerance programs to bacterial pathogens^12,74,75,76^. Balancing appropriate levels of immune reactivity and resolution is critical for restoration of homeostasis^5^, and is in line with the well-appreciated potential for nutritional factors to impact clinical outcomes to infection and sepsis^77,78,79^. Reflecting distinguishable metabolic priorities under conditions of hunger and inflammation, food-restriction and LPS-injection elicited distinct patterns of metabolic responses. Food intake was suppressed in the early 8-hour period following LPS injection, with concomitant reductions in respiratory exchange ratio and increased fatty acid oxidation. In some respects, sickness-induced anorexia may utilize similar organismal level molecular metabolic programs to fasting resulting from nutrient scarcity, including activation of adipose triglyceride lipase-catalyzed lipolysis, peroxisome proliferator-activated receptor (PPAR)-α regulated metabolic programs, ketone body synthesis, and autophagy induction^5^. Differential metabolic responses may be more evident at the level of individual tissues, including hepatic lipid metabolism reprogramming through suppression of PPARα during inflammation to fuel activation of innate immune responses^5,80,81,82^. In line with distinct behavioural states connected to hunger and inflammation, endotoxemia reduced locomotor activity compared to food-restriction. The use of food rewards can complicate interpretation of whether changes in choice behaviour are attributed to decreased appetite (sickness-associated anorexia) rather than altered reward processes. We note this challenge is likely not unique to food rewards, as profound reductions in hydration and deficits in social behaviour also accompany LPS-induced sickness^19,83,84^. However, reward sensitivity was indicated in both choice and latency data within and across patches following LPS injection, suggesting behaviour remained reward-driven, but that thresholds for patch-leaving exploratory choice were increased across patches.

Food restriction encourages learning and performance of instrumental food reward-directed behaviour, and we elected to perform all experiments under the same conditions of food restriction. Recent results have elucidated nutritional control of multiple facets of immunity^77^, and the potential use of calorie restriction for immunosuppressive or longevity benefits continues to attract research interest^9^. Fasting elicits changes in leukocyte migration and distribution^85^, including through suppressed monocyte egress and increased T cell accumulation in bone marrow^86,87^. Although our study utilized a relatively low-dose of LPS, it is likely that the food restriction dietary regimen also restrained inflammatory responses^88,89,90^, emphasizing the metabolic and behavioural impact of even modest inflammatory challenge.

Our behavioural results provide an entry point for understanding neural pathways transducing systemic metabolic states into altered cognitive programs^91,92^. First-order arcuate nucleus neurons integrate hormonal, nutrient, and neural signals to regulate whole-body energy metabolism and behaviour, with elevated agouti-related peptide (AgRP) neuron activity associated with fasting^3,11^. Artificial AgRP neuron activation in sated mice can mimic neural, phenomenological, and behavioural correlates of hunger^93,94^, including voracious feeding and effortful food-directed foraging^17,18^. Consistent with the notion that hunger impelled greater behavioural patch-foraging task engagement, mice performed with faster vigor and achieved a higher number of trials and total reward in baseline, food-restricted sessions compared to pre-feeding. Previous studies have revealed AgRP neurons are a key neural population promoting exploration in response to energetic scarcity, an adaptation advantageous for overcoming aversion to energetic costs and predation risks associated with foraging^95,95,96,97,98^. However, our results also demonstrated acute pre-feeding in chronically food-restricted mice transiently promoted, rather than inhibited, exploratory choice behaviour when patch-foraging. Following encounters with food resources in scarce or uncertain environments, hungry animals may be less willing to forgo smaller, more immediate rewards, in search of larger, but temporally more delayed, rewards. Consistent with this notion, optogenetic AgRP neuron activation in sated mice alters temporal discounting by introducing a bias for more immediate food rewards^99^. In this manner, AgRP neuron activity may encode multiple dimensions of exploratory choice behaviour during foraging in response to energetic scarcity by promoting exploratory behaviour in the absence of food, while conversely inhibiting exploratory behaviour upon patch encounters. AgRP neurons may additionally provide a direct link between control of nutrient homeostasis and peripheral immunity^100^. Although our study did not test manipulations impacting travel costs along temporal or effortful dimensions, it will be interesting to explore whether hunger may attenuate sensitivity to increased travel costs in resource poor environments or risk aversion^101,102,103^.

In contrast to behavioural control of increased appetite during starvation, converging evidence suggests AgRP neuron activity is dispensable for LPS-induced anorexia, which may rely upon extra-arcuate controls. While initial sensing and proinflammatory response to LPS are transduced by TLR4 activation, subsequent signalling via inflammatory mediators activates many neural populations, including immunoceptive vagal and circumventricular organ neurons^83,104,105,106,107,108^. Prominent pro-inflammatory cytokine mediators in endotoxemia include tumor necrosis factor (TNF), interleukin-18 (IL-18), interferon-gamma (IFN-γ), and interleukin-1 beta (IL-1β)^109^. Activation of *Adcyap1*-expressing dorsal-vagal complex neurons recapitulates anorexia and lethargy elicited by LPS^83^, suggesting hindbrain systems serve an important role in controlling appetite and behaviour during endotoxin-triggered inflammation. Systemic LPS administration does not reduce endogenous AgRP neuron activity despite strongly suppressing appetite^110^. Supporting the notion that circuits orthogonal to the leptin-melanocortin pathway control anorexigenic programs resulting from endotoxin-triggered inflammation, artificial AgRP neuron activation is insufficient to overcome LPS-induced anorexia^83,110,111,112^, a phenomenon distinct from other anorexigenic stimuli^112^. LPS additionally blunts AgRP neuron inhibition to food presentation and nutrient ingestion^110^, congruent with altered food valuation and physiological state associated with LPS inflammatory challenge. However, the precise neural controls regulating changes in both appetitive and consummatory drives during inflammation remain to be determined. Additionally, it will be interesting for future studies to address which neural populations, if not canonical first-order arcuate neurons, coordinate systemic metabolic adaptations to endotoxemia.

While these findings suggest dissociable hypothalamic and hindbrain systems mediate the transduction of distinct immunometabolic states in inflammation and hunger to the brain, it is probable that circuits communicating different internal states may converge to influence overlapping forebrain systems important for decision-making. The mesolimbic dopamine pathway comprises ventral tegmental area dopamine neurons projecting to the nucleus accumbens, and is well-implicated in the control of food-reward motivated behaviour^16^. A large body of work suggests a key role of dopamine to mediate reward valuation to food outcomes and effortful invigoration of motivated behaviour^113,114,115^. Acute increases in dopamine may be important for facilitating food-directed exploitative behaviour during foraging^116,117^. Of note, dopaminergic responses are sensitive to metabolic state, with mesoaccumbens dopamine opposingly enhanced by hunger^14,15,118,119,120,121,122,123^ and suppressed by inflammation^22,73,124,125,126^. Striatal dopamine signalling may be involved in adjudicating energetic resources to meet survival priorities, which can be conceptualized in a resource allocation decision framework^127^. The integration of dopaminergic motivational controls with the internal metabolic state of animals, which are in turn modified by reward feedback during interactions with an external environment, could serve to flexibly adjust behaviour in a manner sensitive to both internal and external energy economies^128,129^.

Basal accumbal dopaminergic levels are correlated with reward rate^130,131^, consistent with a hypothesized role of encoding background environmental reward rate^132^. Relatedly, tonic dopamine may control response vigor^132,133^, with greater opportunity cost of lethargic movements in abundant environments^46,134^. In this manner, dopamine may play a critical role in tuning motor response latencies during foraging^135^. Dopamine may additional influence affective states, which have been theorized to impact foraging engagement during interaction with the environment^136^. Dynamic changes in dopamine signalling may also control exploratory choice^137,138^, with some studies indicating decreased dopamine tone promotes exploration or patch-leaving behaviour^139,140,141,142^. Although dopamine may also play a role in novelty-directed exploration^143,144,145,146^, mice in our study had extensive task experience. Striatal dopamine type 1 (D1R) and type 2 receptors (D2R) have different affinities for dopamine; higher affinity D2Rs may be saturated under basal conditions and thus have higher sensitivity to decreases in dopamine tone, while lower affinity D1Rs may have higher responsivity to phasic increases in dopamine release^147^. In this manner, it is possible that complimentary “exploitative” D1R and “exploratory” D2R signalling functions may exert important control of decision-making during foraging^148^, including high-level behavioural control of patch-leaving choice. Relatedly, some theories emphasize D1R and D2R signalling in cost-benefit decision-making through encoding of benefit and cost, respectively^149^. The precise manner in which such dopamine signalling functions are instantiated may involve highly complex dynamics during organism-environment interactions over varying foreground and background reward contexts^150^. Evidence from clinical Parkinson’s Disease patients indicate loss of dorsal striatal dopamine produces perseverative responding and an overharvesting bias, which can be corrected by dopamine replacement medication^151,152^. Evidence from healthy controls additionally suggests dopamine synthesis capacity and receptor availability are correlated with patch foraging decision-making^135^, and that D2R pharmacological manipulation alters patch-leaving behaviour in resource poor environments^137^. Collectively these findings suggest altered cost-benefit computations controlled by mesoaccumbal dopamine signalling may be an attractive hypothesis to explain how hunger and inflammation exercise varying effects on cognition and behaviour. Cortical areas also important in foraging decision-making include the ventromedial prefrontal cortex, anterior cingulate cortex, anterior insular cortex, orbitofrontal cortex, and secondary motor cortex^53,153,154,155,156,157,158,159,160,161,162,163,164^, in addition to neuromodulation from serotoninergic, noradrenergic, and cholinergic systems^57,165,166,167^.

Immune theories of depression emphasize the evolutionary legacy of host defense responses that arose in highly pathogenic environments, and the potential for such relationships to aid in understanding depression etiology^47^. Notable similarities in inflammation-induced sickness behaviour and episodes of major depression include anergia, fatigue, lethargy, low mood, anhedonia, changes in appetite, social withdrawal, and psychomotor slowing^19,20^. Inflammatory challenges often induce depression-related neurovegetative and cognitive symptoms^168^, and both depression and inflammatory challenges are linked with perturbed mesoaccumbal dopamine signalling^22^. Suggesting a potential causal relationship between inflammatory states and depression, risk-factors for depression are uniformly pro-inflammatory^47^, including severe infection and autoimmune disease^169^. Meta analyses indicate patients with MDD present higher levels of C-reactive protein, TNF, and IL-6^170,171,172,173,174^, and PET imaging studies suggest both depression and LPS injection are associated with central neuroinflammation^175,176^. Nonetheless, the clinical presentation of MDD is highly heterogeneous^177^, and may be stratified by immunometabolic status^178^. Atypical depression has stronger association with indices of central inflammation and metabolic syndrome^49^, and over-active inflammation is especially prevalent in MDD patients with resistance to conventional treatments^179,180^.

Symptoms of psychomotor slowing, lethargy, anhedonia, and amotivation are similarly shared between inflammation and depression, and may be jointly mediated via reduced striatal dopamine function^50,177^. In line with this possibility, pre-choice and reward collection response latencies were increased by LPS with concomitant suppression of trial number. Previous studies have demonstrated that systemic inflammation generally reduces incentive motivation and attenuates effortful reward-driven behaviour^181,182,183,184,185,186,187,188,189,190,191^, and that anti-inflammatory treatment in depressed patients can improve willingness to exert effort for rewards^192^. These findings align with the notion that inflammation and depression are marked by dysfunctional amotivation. However, cross sectional theoretical accounts additionally indicate impulsive choice may be increased by inflammation^193,194^. Consistent with this, our data suggest that when interacting with ethological food environments during patch foraging, mice overharvested and in fact worked harder per unit of reward following LPS injection. Reduced exploration is a well-established characteristic of sickness behaviour^195^, which may aid in energy conservation, predator evasion, and possibly mitigating pathogen spread to kin. In this manner, the explore-exploit dimension of ecological decision-making may have revealed a latent hyper-exploitative phenotype resulting from increased exploratory avoidance accompanying sickness behaviour.

Early life stress is a major risk factor for depression^196^, and childhood adversity is associated with reduced exploration during patch foraging^51^. Intriguingly, a recent study examined patch foraging behaviour in a clinical MDD population and discovered a relationship between depression symptomology and foraging choice behaviour; higher depression severity was correlated with leaving patches in more depleted states (overharvesting)^52^. Poorer environmental reward statistics promote increased (more depleted) patch-leaving thresholds in normative models^46^. In the context of depression, an overharvesting bias may be driven by lowered subjective reward rate or a pessimistic bias in evaluating environmental context^52^. Taken further, it is tempting to speculate that measures reflecting inferred background reward rate in patch-foraging paradigms might serve as an implicit behavioural correlate of negative cognitive schema, including persistent negative beliefs about the world/environment, that feature prominently in cognitive theories of depression^197^. The lack of translationally relevant animal models for depression remains a challenge in preclinical research that has led to the development of alternative research frameworks^198^. In this regard, we suggest that inflammatory insult in murine patch foraging paradigms may present an appealing entry point to study changes in environmental belief states.

### Limitations of the study

As the patch foraging paradigm required extensive behavioural shaping, our study exclusively studied behaviour in adult mice. Additionally, the extent to which findings may generalize to female mice remains to be determined^199,200^, and is a critical area for future investigation in light of sex differences in immunity, metabolism, and depression^201,202,203,204^. Our study examined the impact of lipopolysaccharides from *Escherichia coli*, due to their highly potent capacity to modulate of immunometabolic state^12^. However, immunometabolic responses to viral infection are distinct^74,205^ and pathogen-specific host-pathogen interactions may impact metabolism and behaviour^71,206^.

## Resource availability

### Lead Contact

Further information and requests for resources should be directed to and will be fulfilled by the lead contact, Stephanie Fulton.

### Materials availability

This study did not generate any new materials.

### Data and code availability

Datasets and code used to analyze data will be made available by time of publication.

## Methods

### Key resources table

#### Reagent or Resource, Source, Identifier

|  |
| --- |
| **Experimental models: Organisms/strains** |
| Wild-type C57Bl/6J mice, Strain 000664, Jackson Laboratory |
| Heterozygous *Drd1*-Cre mice, 030989-UCD, MMRRC |
| **Commercial assays and reagents** |
| Lipopolysaccharides (LPS) from E Coli (O127:B8), Sigma, L4516 |
| 0.9% Sodium Chloride, Baxter, 00060208 |
| TRIzol, Invitrogen, 15596018 |
| Choloroform, Bioshop, CCL402.1 |
| Random Hexamer Phosphorylated; pd(N)6, GeneLink, 26-4000-10 |
| M-MLV Reverse Transcriptase, Invitrogen, 28025013 |
| dNTP set (100mM), Invitrogen, 10297018 |
| GlycoBlue Coprecipitant, Invitrogen, AM9516 |
| RNAasin Plus RNase Inhibitor, ThermoFisher, PRN2611 |
| Rotor Gene SYBR Green PCR Kit (2000), Qiagen, 204076 |
| **Software and algorithms** |
| Prism 9.3.1, GraphPad Software, https://www.graphpad.com |
| ABET II, Lafayette Instruments, n/a |
| WhiskerServer, Lafayette Instruments, n/a |
| MATLAB, MathWorks, https://www.mathworks.com/products/matlab.html |
| Oxymax for Windows V5.40, Columbus instruments, n/a |
| Rotor-Gene Q Series Software 2.3.1, Qiagen, n/a |
| **Diet** |
| Regular Vanilla Ensure, Abbott Laboratories, 6890713 |

##### Animals

All experimental procedures were approved by the CRCHUM animal care committee, and conducted in accordance with Canadian Council on Animal Care guidelines. A total of 35 adult male mice were used for this study, aged over 3 months at the beginning of task training. The cohort included heterozygous *Drd1*-Cre mice backcrossed on a C57Bl6/J background (N=14, B6.FVB(Cg)-Tg(Drd1-cre)EY262Gsat/Mmucd colony; MMRRC; 030989-UCD), and wild-type C57Bl6/J mice (N=21, strain 000664; Jackson). Mice were housed in a sterile barrier facility with light (reverse 12 hr/12 hr light cycle, lights on 10:00pm) and temperature (22°C) control. Mice had ad libitum access to water in their home-cage throughout. Mice were single-housed throughout behavioural training and experiments to monitor feeding behaviour. In-cage tunnels were used for all mice throughout the duration of training and experiments to reduce handling stress.

##### Dietary Manipulations

Behavioural training and testing: Throughout behavioural training and testing, mice were food restricted to maintain necessary motivation to learn and perform the operant patch-foraging task. Mice were first habituated to handling. Body weight and food intake were measured daily, with food given to maintain mice between ∼85-90% of free-feeding body weight. During initial periods of training, 2% was added per week to the estimated free-feeding body weight to account for normal growth. All behavioural testing occurred during the dark cycle, with food delivered to the home-cage after the conclusion of behavioural training or testing on any days on which animals were trained or tested. Body weight was always measured both prior to and following any behaviour training or testing session. During all behaviour training and testing sessions, Ensure® Regular, Vanilla (Abbott, 6890713) was used as a liquid food reward. Prior to the first behaviour training session, mice received free access to ∼1-1.5g of Vanilla Ensure within their home-cage to prevent neophobia and to learn the incentive value of the food reward prior to behavioural training. Additional dietary manipulations are described in the pre-feeding experiment and LPS behaviour experiment.

##### Behaviour Shaping Protocol

All behavioural training and testing occurred within first-generation Bussey-Saksida touchscreen chambers (Lafayette Instruments) using custom programmed behavioural tasks designed and run in ABET II (Lafayette Instruments). Investigations have previously established the utility of Bussey-Saksida touchscreen systems for the design and execution of standardized, translationally-relevant behavioural tasks for probing cognition and decision-making^54,207,208,209^. The reward troughs were positioned at the rear of the Bussey-Saksida behavioural chambers, and a 3-panel plastic mask was placed inside the front of the chamber to restrict touch responses to 3 equally-sized square areas on the touchscreen matching the size and position of square visual cues displayed on the left, middle, and right part of the touchscreen. All behavioural training and testing occurred during the dark cycle. Mice were trained to perform the touchscreen-based patch-foraging task in sequential stages. Training was organized into three main stages comprised of a total of 11 individual behavioural training tasks, presented to mice in order of increasing complexity.

Training Part A: During the initial stage (including tasks A.1-A.2), mice were first habituated to the touchscreen system and taught to collect food rewards.

Training Part B: During the second stage (including tasks B.1-B.7), mice were taught to make touch responses to a visual cue indicating an open “Patch” in order to harvest food rewards. During all stages of behavioural training, the patch was localized to the lateral (left or right) positions of the touchscreen. Only one patch was available at any time during the task, termed the “Active Patch”, on either the left or right side of the touchscreen. The opposite side of the touchscreen was denoted the “Inactive Patch”, where touch responses would not yield any reward. The left/right position of the active patch on the first trial of a behaviour session was randomized. The position of the active patch alternated back and forth (from left to right, or right to left) within individual behaviour sessions. The criterion for the alternation of the active patch varied depending on the training task. During later parts of the second stage (tasks B.5-B.6), a 5 second time-out punishment was incurred for touch responses to the Inactive (incorrect) Patch position or the centre position of the touchscreen. This punishment was necessary to discourage mice from utilizing a strategy of non-selective touch-responding across the touchscreen or idiosyncratic behavioural chains of touch responding^94^. The time-out punishment featured illumination of a house-light and removal of all visual cues for a 5 second period, and re-setting of the patch FR operant counter. Selectivity for patch touch responding to the Active Patch rather than the Inactive Patch was calculated with a measure termed “Active Patch Discrimination”, computed as the ratio of touch responses during a behaviour session to the Active Patch divided by the total number of touch responses to both the Active and Inactive Patch positions during the decision period.

Training Part C: During the third stage (including tasks C.1-C.2), mice were taught to make touch responses to a visual cue indicating the possibility to “Travel”, in order to open a new, replenished “Patch”. During this stage of behavioural training (Training Part C), the middle position of the touchscreen denoted the patch-leaving, “travel” position. The term “Travel” is borrowed from terminology commonly used in behavioural ecology literature, and is meant to simulate a time-cost associated with traveling between patches. An overview of the training stages is listed in Table 1, with a brief description of training schedule elements.

**Table 1.** Overview of Behaviour Shaping Protocol.

| Training Stage | Training Part A (Habituation) |  | Training Part B (Learning to harvest from active patch) |  |  |  |  |  |  | Training Part C (Learning to travel) |  |
| --- | --- | --- | --- | --- | --- | --- | --- | --- | --- | --- | --- |
| Training Task | A.1 | A.2 | B.1 | B.2 | B.3 | B.4 | B.5 | B.6 | B.7 | C.1 | C.2 |
| Session Length | 10 min | 15-30 min | 30 min | 30 min | 30 min | 30 min | 30 min | 30 min | 30 min | 30 min | 30 min |
| Sessions/Mouse | 1 | 3-4 | 1 | 3 | 1 | 2 | 3 | 22 | 6 | 17 | 17 |
| Trial (Decision) Timer | n/a | n/a | 30s | unlimited | unlimited | unlimited | unlimited | unlimited | unlimited | unlimited | unlimited |
| Patch Visual Cue | n/a | n/a | White square | White square | White square | White square | White square | White square | White square | White square | White square |
| Patch Touch Tone | n/a | n/a | n/a | n/a | 40ms tone A | 40ms tone A | 40ms tone A | 40ms tone A | 40ms tone A | 40ms tone A | 40ms tone A |
| Patch Touch FR Criterion | n/a | n/a | If touch 3x reward | FR1 | FR1 | FR2 | FR3 | FR5 | FR5 | FR5 | FR5 |
| Patch FR Operant Outcome | n/a | n/a | Ensure reward | Ensure reward | Ensure reward | Ensure reward | Ensure reward | Ensure reward | Ensure reward | Ensure reward | Ensure reward |
| Reward Delay | n/a | n/a | 0s | 0s | 0.5s | 0.5s | 0.5s | 0.5s | 0.5s | 0.5s | 0.5s |
| Reward Probability | n/a | 1 | 1 | 1 | 1 | 1 | 1 | 1 | 1 | 1 | 1 |
| Initial Reward Volume | n/a | 20µl | 20µl or 60µl | 20µl | 20µl | 20µl | 20µl | 20µl | 20µl | 15µl | 15µl |
| Reward Depletion | n/a | n/a | n/a | n/a | n/a | n/a | n/a | n/a | n/a | -15µl, every 6-10 harvests (mean 8) | -15µl, every 6-10 harvests (mean 8) |
| Reward Availability Cue | n/a | 1s tone A/traylight | 1s tone A/traylight | 1s tone A/traylight | 1s tone A/traylight | 1s tone A/traylight | 1s tone A/traylight | 1s tone A/traylight | 1s tone A/traylight | 1s tone A/traylight | 1s tone A/traylight |
| Travel Visual Cue | n/a | n/a | n/a | n/a | n/a | n/a | n/a | n/a | n/a | White square (available following patch depletion) | White square (always available) |
| Travel Touch Tone | n/a | n/a | n/a | n/a | n/a | n/a | n/a | n/a | n/a | 40ms tone B | 40ms tone B |
| Travel Touch FR Criterion | n/a | n/a | n/a | n/a | n/a | n/a | n/a | FR5 | FR3 | FR5 | FR1 |
| Travel FR Operant Outcome | n/a | n/a | n/a | n/a | n/a | n/a | n/a | 5s time-out punishment with houselight | 5s time-out punishment with houselight | Initiates travel time cost | Initiates travel time cost |
| Travel Time Cost | n/a | n/a | n/a | n/a | n/a | n/a | n/a | n/a | n/a | 5s | 5s |
| Travel Completion Availability Cue | n/a | n/a | n/a | n/a | n/a | n/a | n/a | n/a | n/a | 1s tone B/traylight | 1s tone B/traylight |
| Inactive (Incorrect) Patch Visual Cue | n/a | n/a | n/a | n/a | n/a | n/a | n/a | n/a | n/a | n/a | n/a |
| Inactive Patch Touch Tone | n/a | n/a | n/a | n/a | n/a | n/a | n/a | n/a | n/a | n/a | n/a |
| Inactive Patch Touch FR Criterion | n/a | n/a | n/a | n/a | n/a | n/a | n/a | FR3 | FR1 | n/a | n/a |
| Inactive Patch Operant Outcome | n/a | n/a | n/a | n/a | n/a | n/a | n/a | 5s time-out punishment with houselight | 5s time-out punishment with houselight | n/a | n/a |
| Active Patch L/R Position Switch | n/a | n/a | Every 10 trials | Every 10 trials | Every 10 trials | Every 10 trials | Every 10 trials | Every 10 trials | Every 10 trials | After travel completion | After travel completion |
| Inter-Trial Interval | n/a | n/a | 4.5 sec | 4.5 sec | 4.5 sec | 4.5 sec | 4.5 sec | 4.5 sec | 4.5 sec | 4.5 sec | 4.5 sec |

##### Schedule 1 Protocol: 1 Patch Type (15µl)

Food-restricted mice were in a fasted state during the start of all behaviour sessions, and food-restriction food was provided immediately following the end of behaviour sessions. Mice were weighed just prior to beginning behaviour sessions, with all sessions occurring in the dark cycle. At the start of the behaviour session visual cues indicated the Active Patch position (left or right, P_A_ in Figure 1) and the Travel cue (centre, T in Figure 1) displayed on a touchscreen at the front of the touchscreen chamber. The initial left/right position of the Active Patch was randomized across sessions, and the Inactive Patch position (P_I_ in Figure 1) was located laterally opposite to the Active Patch. Mice made touch responses to the visual cues to indicate a binary choice decision to either harvest from the Active Patch (fixed ratio 5 operant criterion) or to Travel to a new patch (fixed ratio 1 operant criterion) following an incurred 5 second time cost. Touch responses to the Active Patch cue triggered cue offset and a 40ms tone, while touch responses to the travel cue triggered image offset and a distinct 40ms tone. Touch responses to the Inactive Patch position (with no visual cue) were recorded but did not produce any outcome. Active patch discrimination, a measure of discriminative touch responding during the task, was calculated as the total number of touch responses to the Active Patch cue, divided by the total touch responses to both the Active and Inactive Patch positions during the decision time, multiplied by 100.

All Schedule 1 Tasks ran for a duration of 30 minutes, however all trials were self-paced and there were no limitations on the number of trials animals could perform. If mice indicated their choice to harvest from the Active Patch by making 5 touch responses, a food reward would subsequently be delivered. Vanilla Ensure liquid reward was used throughout all behaviour sessions, delivered from a pump into a reward trough at the rear of the touchscreen chamber. In the Schedule 1 Task, all patches began with an initial reward volume of 15µl. Onset of reward delivery occurred 500ms following active patch operant criterion achievement (5 touch responses to Active Patch cue)t, and was marked by a reward tone and illumination of a traylight. Time of entry into the reward tray to collect food reward was recorded, and initiated a 4.5s inter-trial interval separating trials.

To describe the sequential decisions of mice performing this task, trials were organized according to “harvest levels” within a patch, with harvest level 1 being the initial, full patch (15µl Ensure). Patches depleted linearly by −1µl for every subsequent harvest from the active patch. In this manner, harvest level 2 would indicate a patch with −1µl depletion (14µl Ensure if mouse continued to harvest from patch), harvest level 3 would indicate a patch with −2µl depletion (13µl Ensure), harvest level 4 would indicate a patch with −3µl depletion (12µl Ensure), and so on. In this manner, the reward feedback mice received from continuing to harvest from a patch declined over subsequent harvests from the same Active Patch. At any time, mice could choose to leave an Active Patch to open to a new, replenished patch by choosing to “Travel”. A touch response to the travel cue would initiate a 5s time cost, after which the availability of the ability to complete the travel would be indicated by a 1s travel tone (distinct from reward tone) and traylight illumination. Mice were required to enter the (unrewarded) reward tray to complete the travel. The time of entry into the tray was recorded and would initiate a 4.5s inter-trial interval separating trials. The start of the next trial following a trial would be marked by the opening of a new, replenished patch on the laterally opposite position on the touchscreen. In other words, if the active patch was previously on the left side of the touchscreen, it would switch to the right side following a travel, or vice-versa.

Mice could fully deplete patches if they chose to, by perseverative continuation of harvesting 15 times from an Active Patch. Once a patch was fully depleted (harvest level 16, 0µl Ensure), mice could still continue to harvest from the unrewarded (empty) patch for an additional 10 trials (i.e. up to harvest level 25). Unrewarded harvests utilized the same reward cues as rewarded patch harvests, and mice were still required to enter the unrewarded tray to initiate the inter-trial interval to start the following trial. If mice completed 10 unrewarded harvests, it would initiate a “forced travel” sequence in which only the travel cue was available, and additional touch responses to the formerly available active patch position would not produce any outcome. In the “forced travel” sequence, mice were required to make a touch response to the travel cue and complete a travel to proceed in the task. Forced travel trials were not included in choice data analysis, and if included would comprise less than 0.1% of total trials in the Schedule 1 Task validation dataset. Sessions ended at the termination of a 30 minute timer, regardless of task state. Mice were weighed immediately following conclusion of the task, and chambers were thoroughly cleaned with 70% ethanol between sessions. Loss of body weight due to excrement during behaviour tasks was not measured. Regular food-restriction chow was provided to the home-cage following behaviour. A total of 15 behaviour sessions from 20 mice were analyzed in the Schedule 1 Task validation dataset.

##### Schedule 2 Protocol: 3 Patch Types (6µl, 12µl, 18µl)

The structure of the Schedule 2 Task was very similar to Schedule 1. Mice made touch responses to indicate their choice to harvest from an Active Patch or to Travel, however in contrast to Schedule 1, three distinct patch types were introduced which varied in their initial reward volume of either 6µl, 12µl or 18µl Ensure reward. Patches were selected from a 9-item list comprised of 3 patches of each type, by random without replacement, ensuring mice would see an equal number of each patch type in blocks of 9 travels (or 8 travels for the first block). All patch types linearly depleted by −1µl for every subsequent harvest, and as in Schedule 1, harvest levels were used to describe trials with reference to patch depletion from the initial, full Active Patch. In other words, a 6µl Active Patch would begin with 6µl reward (harvest level 1), depleted by −1µl following a harvest to 5µl reward (harvest level 2), depleted by −2µl following an additional harvest to 4µl reward (harvest level 3), and so on. Similarly, an 18µl Active patch would begin with 18µl reward (harvest level 1), depleted to −1µl following a harvest to 17µl reward (harvest level 2), depleted by −2µl following an additional harvest to 16µl reward (harvest level 3), and so on. Patch types were indicated by distinct visual cues of lines of varying orientation, however mice appeared to primarily utilize reward feedback rather than visual cues to guide choice behaviour (Figure S8E). Following a Travel, a new patch would be selected and opened on the opposite side of the touchscreen, in a similar manner to Schedule 1. All sessions in the Schedule 2 Task were run for 40 minutes. A total of 18 behaviour sessions from 19 mice were analyzed in the Schedule 2 Task validation dataset. An overview of Schedule 1 and Schedule 2 behaviour tasks is listed in Table 2.

**Table 2.** Overview of Behaviour Tasks.

| Behavioural Task: | Schedule 1 Task | Schedule 2 Task |
| --- | --- | --- |
| Session length: | 30 min | 40 min |
| Sessions/mouse: | 15 sessions/20 mice for task validation;<br>2 baseline and 2 prefed sessions/35 mice for Pre-feeding experiment | 18 sessions/19 mice for task validation;<br>1 saline (24hr), 1 LPS (24hr), and 1 LPS (48hr) session/14 mice for LPS experiment |
| Trial (decision) Timer: | Unlimited | Unlimited |
| Patch Type: | 15µl | 6µl, 12µl, or 18µl |
| Patch Probability: | 1 | ~0.33 (3x each patch in 9 item list, random without replacement) |
| Patch Visual Cue: | Parallel horizontal lines | Parallel 30°, 90°, or 150° lines for 6µl, 12µl, or 18µl patch type, respectively |
| Patch Touch Tone: | 40ms tone A | 40ms tone A |
| Patch Touch FR Criterion: | FR5 | FR5 |
| Patch FR Operant Outcome: | Ensure reward | Ensure reward |
| Reward Delay: | 0.5s | 0.5s |
| Reward Probability: | 1 | 1 |
| Initial Reward Volume: | 15µl | 6µl, 12µl, or 18µl, depending on patch type |
| Reward Depletion: | -1µl per harvest | -1µl per harvest |
| Reward Availability Cue: | 1s tone A/traylight | 1s tone A/traylight |
| Travel Visual Cue: | Parallel horizontal lines | Parallel horizontal lines |
| Travel Touch Tone: | 40ms tone B | 40ms tone B |
| Travel Touch FR Criterion: | FR1 | FR1 |
| Travel FR Operant Outcome: | Initiates travel time cost | Initiates travel time cost |
| Travel Time Cost: | 5s | 5s |
| Travel Completion Availability Cue: | 1s tone B/traylight | 1s tone B/traylight |
| Inactive (incorrect) Patch Visual Cue: | n/a | n/a |
| Inactive Patch FR Criterion: | n/a | n/a |
| Inactive Patch Operant Outcome: | n/a | n/a |
| Active Patch L/R Position Switch: | After travel completion | After travel completion |
| Inter-Trial Interval: | 4.5s | 4.5s |

##### Schedule 1 Task Protocol: Pre-feeding experiment

To assess the impact of an acute manipulation of metabolic state, in a separate experiment, mice were tested for the effect of acute chow food availability 1 hour prior to behaviour on the Schedule 1 Task. For all mice the Pre-feeding experiment occurred following a minimum of 15 prior behaviour sessions on the Schedule 1 Task, ensuring similar task experience across mice. Four behaviour sessions (2 baseline sessions, and 2 pre-fed sessions in alternating order) were performed. The order of the sessions for all mice was Baseline session 1, Pre-fed session 1, Baseline session 2, Pre-fed session 2, with all sessions being performed in the same week for respective mice. Schedule 1 behaviour was performed as previously described, with the addition of one additional measurement of body weight performed 1 hour before the start of behaviour for all sessions. For pre-fed sessions, ad libitum chow food was returned to the home-cage for 1 hour before the start of behaviour. Mice were returned to regular food-restriction at the conclusion of each behaviour session, with ad libitum chow food removed before returning each animal to home-cage following pre-fed behaviour sessions. As in other behaviour tasks, body weight was measured immediately before and following behaviour sessions.

##### LPS Preparation and Injection

Lipopolysaccharides (LPS) from E. Coli (O127:B8, Sigma, L4516) were prepared in 0.9% physiological saline at a stock concentration of 0.033mg/mL, with aliquots stored at −20°C and thawed to room temperature prior to single use. For all experiments with LPS administration, mice were intraperitoneally injected at a dose of 0.33mg/kg, which has been previously demonstrated is sufficient to robustly alter food-directed motivated behaviour^188,210^. To prevent variance due to batch effects, the same batch of LPS was used for all experiments. Volume-matched sterile 0.9% physiological saline was used for control intraperitoneal injections. To mitigate confounds related to endotoxin tolerance^211^, repeat injections of LPS were separated by a minimum of 8 weeks. The order of LPS injections for all mice began with behaviour, followed by CLAMS.

##### Schedule 2 Task Protocol: LPS experiment

To assess the impact of different metabolic states associated with food restriction and systemic endotoxin-induced inflammation, food-restricted mice were tested on the Schedule 2 Task following injections of saline or LPS. Mice were extensively habituated to the stress of handling and intraperitoneal injections prior to the experiment. Mice received at least one refresher Schedule 2 Task behaviour session prior to the start of the experiment. For all experiments the saline injection was performed prior to the LPS injection, with both saline and LPS behaviour experiments performed within the same week for each respective mouse. All injections were performed around 2:00pm during the dark cycle (ZT16), with corresponding behaviour experiments performed 24 hours following injection. Food intake and body weight was measured 8 hours following injections. Similar to all behaviour experiments, food and body weight were measured immediately prior to behaviour sessions, with body weight also measured immediately following behaviour sessions. Feed efficiency was calculated as the ratio of the change in body weight (g) divided by the cumulative food intake (kcal) consumed over the respective time period. Regular food restriction chow-food was provided following behaviour sessions. Following LPS injection, an additional behaviour session was run 48 hours following injection. As trial numbers were reduced (Figure 6G-H), coupled with a trend towards increased forced travel frequency (Figure S19H-I) observed at 24 hours following LPS injection, forced travels were included in all data sessions for the analysis of the LPS behaviour experiment (Figure 6). Forced travels were coded in data analysis as travels at harvest level 17, 23, and 29 in 6µl, 12µl, and 18µl patches, respectively, which we acknowledge may underestimate the extent of hyper-exploitation induced by LPS.

##### Body Composition

Prior to the beginning of metabolic phenotyping, mice were chronically food-restricted to ∼90% of free-feeding body weight for at least two weeks, with food given daily between 2:00-3:00pm. Fat and lean mass were evaluated using a nuclear magnetic resonance apparatus (EchoMRI) three days prior to the start of the indirect calorimetry experiment. Measurement of body composition occurred approximately 30-60 minutes prior to delivery of daily food under the usual food restriction regimen.

##### Metabolic Phenotyping

Metabolic phenotyping with indirect calorimetry was performed using the Comprehensive Lab Animal Monitoring System (CLAMS, Columbus Instruments) at 22 °C. Experimental conditions were designed to recapitulate as closely as possible the metabolic and immunological states produced by the LPS behavioural experiment. Animals were maintained on reverse light cycle (12 hr light/12 hr dark cycle, lights on at 10:00pm) throughout the CLAMS experiment. The light cycle was synchronized with reference to the first animal in each cohort. Animals were habituated for over 24 hours in the CLAMS prior to the first injection. Animals were maintained on food restriction at the beginning of the CLAMS experiment with food first becoming available at 2:00pm (ZT16) for the first mouse. The programmed times were staggered by an additional 8 minutes for each additional mouse. Food continued to be available within the CLAMS chamber up to a 6 hr time period limit (ending at 8:00pm (ZT22) for the first mouse). An additional limit was programmed to terminate food availability when a pre-determined food mass limit for each mouse was reached to prevent mice from gaining above an estimation of approximately 95% of free-feeding body weight during the experiment, however this limit was not engaged in any mice during the experiment. Following this period, food became unavailable again in the CLAMS until the following day, beginning at 2:00pm for the first mouse. Following the 24 hour habituation period, mice were removed from the CLAMS cage with a volume-matched intraperitoneal injection of 0.9% saline (Baxter) performed approximately 5 minutes prior to the food availability of each respective mouse. The same cycle of food availability, with identical programmed restriction parameters, was repeated over the 3 days (Post-saline day 1, 2, 3), representing a time period of 72 hours following saline injection. At 72 hours following saline injection, mice received intraperitoneal injection of LPS (0.33mg/kg). Feeders were converted to an ad libitum availability for the following duration of the experiment, comprising over 48 hours post LPS injection.

##### CLAMS Data Analysis

CLAMS data was extracted from Oxymax software (Columbus Instruments). Data shown and analyzed in figures was aligned to the nearest cycle of the food availability period of the first mouse of each cohort. Analysis of 18 consecutive CLAMS cycles, representing ∼7.7 hours, is referred to as 8 hours within text and figures. Locomotor activity was calculated form the sum of both X and Z ambulatory beam breaks. Fatty acid oxidation was calculated using the formula FAO=heat*(1-RER)/0.3. CLAMS feeder data was additionally analyzed in MATLAB (Mathworks) using custom scripts for meal pattern analysis^212^. Meals were defined as the sum of feeding bouts with summed mass exceeding 0.03g, with duration exceeding 10 seconds, and occurring in a time period less than 5 minutes. Time bins for meal analysis were programmed for each mouse to correspond to the onset of food availability with the CLAMS.

##### Tissue Collection

Mice were food-restricted for a period of at least two weeks to recapitulate the chronic metabolic state during behaviour and metabolic phenotyping, with food and body weight recorded around 2:00-3:00pm (ZT16-17) daily. On sacrifice day, mice were randomly assigned to either receive intraperitoneal volume-matched saline or LPS (0.33mg/kg) injections between 2:00-3:30pm, with injection times staggered across mice. Mice received ad libitum home-cage food following injections. Three hours following injection time, body weight and food intake were measured, after which mice were sacrificed under isoflurane anesthesia. Brains were rapidly extracted and snap frozen in isopentane with dry ice, and stored at −80 °C.

##### Biochemical Analyses

RT-qPCR experiments were performed to assess RNA content within the arcuate nucleus (ARC), ventral tegmental area (VTA), and nucleus accumbens (NAc). Following cryosectioning of coronal brain sections (100µm), tissue punches containing the ARC, VTA, and NAc were acquired with a biopsy punch and stored at −80°C. Punches were homogenized and RNA extracted with TRIzol (Invitrogen), with cDNA prepared using M-MLV Reverse Transciptase (Invitrogen) and random hexamers (GeneLink). Real-time quantitative-PCR reactions were performed using Rotor Gene SYBR Green PCR (Qiagen). Primers used for RT-qPCR experiments are listed in Supplementary Table 1. The first 10 cycles for all samples were not included in analysis. Expression was normalized to cyclophilin, and all RT-qPCR data is expressed as relative fold change to saline injection control (normalized at 1.0).

##### Exclusion Criteria

For the calculation of mean time to travel across patch types in schedule 2 data, outliers were identified by ROUT with Q = 1% (Graphpad Prism). For the calculation of correlations in Figure 6, values associated with linear regression residuals with absolute value greater than 40 were considered outliers and not included in analysis. For the purpose of visual clarity only rewarded harvest levels are shown in many graphs, however choice trials from all harvest levels were included in analysis.

##### Quantification and Statistical Analyses

Individual session raw behavioural data was first exported from ABET II using the “Final Values” function. All other specific choice and latency behavioral data were extracted from ABET II using custom Analysis Set Design programs. Extracted data were then processed in Microsoft Excel. Summary data was first calculated as biological replicate means, with the group mean calculated from biological replicate means and then typically visualized as group mean in heatmaps, or mean ± standard error of mean (SEM) for bar and line graphs. Correlational data was analyzed using Pearson’s correlation test and simple linear regression. Comparisons of within-subject data at two different time points were typically analyzed using paired two-sided t-test, within-subject data with more than two time points were typically analyzed using one-way ANOVA. For some analyses of Schedule 2 Task data, trials were standardized with reference to the reward state of the 18µl patch in order to analyze trials from all three patch types in the same measure. Differences were considered significant at p<0.05 for all comparisons.

## Acknowledgments

DL, RM, and DM were supported by PhD scholarships from Fonds de Recherche Québec Santé (FRQS), and JR was supported by a postdoctoral scholarship from FRQS. FB was supported by MSc fellowship from CRCHUM. We are grateful to Peter Shizgal and Becket Ebitz for thoughtful comments and suggestions. We thank the small animal phenotyping and imaging core facilities at CRCHUM for use of EchoMRI and CLAMS.

## Author Contributions

DL and SF designed studies. DL collected behavioural data. DL, PG, and DP wrote programs for ABET II data analysis. DL wrote ABET II behaviour schedules with assistance from DP. DL, ALF, PG, RM, DM, FB, and SF analyzed data. YZ and DL performed computational analyses. DP, SN, and JR provided critical guidance on experimental design. RM wrote programs for meal pattern analysis and analyzed CLAMS meal pattern data. DM, FB, DL and SN performed biochemical experiments. DL and SF wrote manuscript. All authors critically contributed to the review of manuscript.

