## Supplemental information for "Immunometabolic state modulation of sequential decision making in patch-foraging mice"

#### Supplementary Figure Legends

**Figure S1. Additional schedule 1 metabolic and behavioural data.** Related to Figure 1. **A.** Pre-session body weight. **B.** Relative pre-session body weight (expressed as a percentage of free-feeding body weight). **C.** Relative post-session body weight. **D.** Total Ensure reward earned per session. **E.** Mean total reward earned per session from (**D**). **F.** Change in absolute body weight from pre-session (**A**) to post-session. **G.** Total reward earned per session (related to **D**) and change in body weight from pre- to post-session period (mean shown in **F**). **H.** Daily chow food eaten during food-restriction during day prior to each behaviour session. **I.** Proportion of total daily calories from Ensure reward, calculated as the reward earned (kcal) during behaviour sessions and total food (kcal) consumed each day (from behaviour and food-restriction chow). Data represented as mean ± SEM (**A-F, H-I**) or individual points (**G**). Black line and shaded area represent model and 95% confidence interval, respectively, from simple linear regression (**G**). Output of two-tailed Pearson's correlation shown top right in (**G**).

**Figure S2. Summaries of schedule 1 mean harvest and travel choice probabilities for individual mice.** Related to Figure 1. **A.-T.** Individual mean choice probabilities for Mouse 1-20, calculated from schedule 1 behaviour sessions 1-15. Data represented as mean (**A-T**).

**Figure S3. Schedule 1 mean latency data for individual mice.** Related to Figure 1. **A.** Individual mean latency to first touch response for mouse 1-20. **B.** Individual mean latency to earn decision for mouse 1-20. **C.** Individual mean effort time for mouse 1-20. **D.** Individual mean latency to collect reward for mouse 1-20. **E.** Individual mean latency to collect reward for mouse 1-20, including unrewarded harvests. Dotted line indicates threshold for complete patch depletion after harvest level 15. **F.** Overall mean latency to collect reward from (**E**). **G.** Individual mean latency to complete travel. Data represented as mean (**A-T**). Hatched areas indicate no data.

**Figure S4. Additional schedule 1 mean latency data sorted by session block.** Related to Figure 2. **A.** Mean latency to first touch response by session block. **B.** Mean latency to earn decision by session block. **C.** Mean effort time by session block. **D.** Mean latency to collect reward by session block. **E.** Mean latency to complete travel by session block. Data represented as mean (**A-E**).

**Figure S5. Additional pre-feeding experiment metabolic and behavioural data.** Related to Figure 3. **A.** Body weight 1hr pre-behaviour. **B.** Absolute body weight change from period 1hr to 0hr pre-behaviour. **C.** Relative body weight change (% free feeding body weight) from period 1hr to 0hr pre-behaviour. **D.** Absolute body weight change from period 1hr to 0hr pre-behaviour in (**B**). **E.** Absolute body weight 0hr pre-behaviour. **F.** Absolute body weight 0hr pre-behaviour in (**E**). **G.** Active patch touch frequency per session. **H.** Active patch touch frequency in (**G**). **I.** Active patch discrimination per session. **J.** Active patch discrimination in (**I**). K. Harvest frequency per session. **L.** Mean harvest frequency per session in (**K**). **M.** Travel touch frequency per session. **N.** Travel touch frequency in (**M**). **O.** Travel frequency per session. **P.** Travel frequency in (**O**). **Q.** Trial frequency per session. **R.** Trial frequency in (**Q**). S. Mean harvest choice probability (0-5 min behaviour). **T.** Mean harvest choice probability (0-30 min behaviour). Data represented as mean ± SEM (**A-R**) or mean (**S, T**); ***p<0.001, ****p<0.0001. Paired two-tailed t-test (**D, F, H, J, L, N, P, R**).

**Figure S6. Latency data from pre-feeding experiment.** Related to Figure 3. **A.** Mean latency to first touch response (0-30 min behaviour). **B.** Mean latency to earn decision (0-30 min behaviour). **C.** Mean effort time (0-30 min behaviour). **D.** Mean latency to collect reward (0-30 min behaviour). **E.** Mean latency to complete travel (0-30 min behaviour). Data represented as mean (**A-E**).

**Figure S7. Additional pre-feeding experiment correlation data.** Related to Figure 3. **A.** Relative body weight (% of free-feeding) at 1hr pre-behaviour and mean harvest level at travel (0-5 min behaviour). **B.** Absolute body weight at 1hr pre-behaviour and mean harvest level at travel (0-5 min behaviour). **C.** Absolute body weight 0hr pre-behaviour and mean harvest level at travel (0-5 min behaviour). **D.** Absolute body weight change from period 1hr to 0hr pre-behaviour and mean harvest level at travel (0-5 min behaviour). **E.** Relative body weight (% free-feeding) 1hr pre-behaviour and mean harvest level at travel (0-30 min behaviour). **F.** Absolute body weight 1hr pre-behaviour and mean harvest level at travel (0-30 min behaviour). **G.** Absolute body weight 0hr pre-behaviour and mean harvest level at travel (0-30 min behaviour). **H.** Absolute body weight change from period 1hr to 0hr pre-behaviour and mean harvest level at travel (0-30 min behaviour). Data represented as individual points (**A-H**). Black lines and shaded areas represent models and 95% confidence intervals, respectively, from simple linear regression (**A-H**). Output of two-tailed Pearson's correlation shown top right in graphs (**A-H**).

**Figure S8. Additional schedule 2 behavioural data.** Related to Figure 4. **A.** Active patch touch frequency per session. **B.** Active patch discrimination per session. **C.** Travel touch frequency per session. **D.** Mean latency to travel choice by patch type, calculated as the time from the start of trial at harvest level 1 to the travel choice time. **E.** Travel choice probability at harvest level 1 by patch type. **F.** Reward/effort ratio by patch type. **G.** Effort/reward ratio by patch type. **H.** Mean harvest level at travel (all trials standardized to 18µl patch reward). **I.** Mean harvest level at travel by patch type (all trials standardized to18µl patch reward). **J.** Mean harvest level at travel by session block (6µl patch). **K.** Mean harvest level at travel by session block (12µl patch). **L.** Mean harvest level at travel by session block (18µl patch). **M.** Mean travel choice probability by session block (6µl patch). **N.** Mean travel choice probability by session block (12µl patch). **O.** Mean travel choice probability by session block (18µl patch). Data represented as mean ± SEM (**A-L**) or mean (**M-O**). One-way ANOVA with Geisser-Greenhouse correction (**D-G, I-L**), main effect of patch type shown in graph (**D, F, G, I**).

**Figure S9. Summaries of schedule 2 mean travel choice probabilities for individual mice.** Related to Figure 4. **A.-S.** Mean travel choice probabilities by patch type for individual mice (schedule 2 behaviour sessions 1-18). Data are represented as mean (**A-S**).

**Figure S10. Schedule 2 mean latency to first touch response data for individual mice.** Related to Figure 4. **A.-S.** Mean latency to first touch response by patch type for individual mice (schedule 2 behaviour sessions 1-18). Data are represented as mean (**A-S**).

**Figure S11. Schedule 2 mean latency to earn decision data for individual mice.** Related to Figure 4. **A.-S.** Mean latency to earn decision by patch type for individual mice (schedule 2 behaviour sessions 1-18). Data are represented as mean (**A-S**).

**Figure S12. Schedule 2 mean effort time data for individual mice.** Related to Figure 4. **A.-S.** Mean effort time by patch type for individual mice (schedule 2 behaviour sessions 1-18). Data are represented as mean (**A-S**).

**Figure S13. Schedule 2 mean latency to collect reward data for individual mice.** Related to Figure 4. **A.-S.** Mean latency to collect reward by patch type for individual mice (schedule 2 behaviour sessions 1-18). Data are represented as mean (**A-S**).

**Figure S14. Schedule 2 mean latency to complete travel data for individual mice.** Related to Figure 4. **A.-S.** Mean latency to complete travel by patch type for individual mice (schedule 2 behaviour sessions 1-18). Data are represented as mean (**A-S**).

**Figure S15. Additional behavioural and metabolic CLAMS data in 1hr time bins.** Related to Figure 3 and 5. **A.** Food intake (1hr time bins around select transition points). **B.** Mean food intake 1hr before and after food availability (mean calculated from post-saline day 1-3; -1-0h time bin (left): -1-0h, 23-24h, 47-48h; 0-1h time bin (right): 0-1h, 24-25h, 48-49h). **C.** Locomotor activity (1hr time bins around select transition points). **D.** Mean locomotor activity 1hr before and after food availability (mean calculated from post-saline day 1-3; -1-0h time bin (left): -1-0h, 23-24h, 47-48h; 0-1h time bin (right): 0-1h, 24-25h, 48-49h). **E.** Energy expenditure (1hr time bins around select transition points). **F.** Mean energy expenditure 1hr before and after food availability (mean calculated from post-saline day 1-3; -1-0h time bin (left): -1-0h, 23-24h, 47-48h; 0-1h time bin (right): 0-1h, 24-25h, 48-49h). **G.** Respiratory exchange ratio (1hr time bins around select transition points). **H.** Mean RER 1hr before and after food availability (mean calculated from post-saline day 1-3; -1-0h time bin (left): -1-0h, 23-24h, 47-48h; 0-1h time bin (right): 0-1h, 24-25h, 48-49h). **I.** Fatty acid oxidation (1hr time bins around select transition points). **J.** Mean FAO 1hr before and after food availability (mean calculated from post-saline day 1-3; -1-0h time bin (left): -1-0h, 23-24h, 47-48h; 0-1h time bin (right): 0-1h, 24-25h, 48-49h). Shaded areas indicate food availability within CLAMS. Data represented as mean ± SEM (**A-J**); $^{**}p<0.01$, $^{****}p<0.0001$. Paired two-sided t-test (**B, D, F, H, J**).

**Figure S16. CLAMS meal pattern data.** Related to Figure 5. **A.** Food intake during feeding period of post-saline day 1-3. **B.** Food intake during feeding period of post-saline day 1 and 3. **C.** Mean meal frequency during CLAMS food availability periods in 6hr time bins. **D.** Mean meal frequency during 6 hr period following saline (0-6h) or LPS (72-78h) injections. **E.** Mean meal frequency during post-saline day 3 feeding period (48-54h) and corresponding period following LPS injection (72-78h). **F.** Mean meal size during CLAMS food availability periods in 6hr time bins. **G.** Mean meal size during 6 hr period following saline (0-6h) or LPS (72-78h) injections. **H.** Mean meal size during post-saline day 3 feeding period (48-54h) and corresponding period following LPS injection (72-78h). **I.** Mean meal duration during CLAMS food availability periods in 6hr time bins. **J.** Mean meal duration during 6 hr period following saline (0-6h) or LPS (72-78h) injections. **K.** Mean meal duration during post-saline day 3 feeding period (48-54h) and corresponding period following LPS injection (72-78h). Data represented as mean ± SEM (**A-K**); *p<0.05, ***p<0.001. Paired two-sided t-test (**B, D, E, G, H, J, K**).

**Figure S17. Additional CLAMS behavioural and metabolic data.** Related to Figure 5. **A.** Cumulative food intake during 8 hr period following either saline injection (left, 0-8 hr) or LPS injection (right, 72-80 hr). **B.** Food intake during end of day period for post-saline day 1 (left, 16-24 hr) or post-LPS day 1 (right, 88-96 hr), corresponding to period 16-24 hr following injection. **C.** Locomotor activity during 8 hr period following either saline injection (left, 0-8 hr) or LPS injection (right, 72-80 hr). **D.** Locomotor activity during end of day period for post-saline day 1 (left, 16-24 hr) or post-LPS day 1 (right, 88-96 hr), corresponding to period 16-24 hr following injection. **E.** Energy expenditure during 8 hr period following either saline injection (left, 0-8 hr) or LPS injection (right, 72-80 hr). **F.** Energy expenditure during end of day period for post-saline day 1 (left, 16-24 hr) or post-LPS day 1 (right, 88-96 hr), corresponding to period 16-24 hr following injection. **G.** RER during 8 hr period following either saline injection (left, 0-8 hr) or LPS injection (right, 72-80 hr). **H.** RER during end of day period for post-saline day 1 (left, 16-24 hr) or post-LPS day 1 (right, 88-96 hr), corresponding to period 16-24 hr following injection. **I.** Fatty acid oxidation during 8 hr period following either saline injection (left, 0-8 hr) or LPS injection (right, 72-80 hr). **J.** Fatty acid oxidation during end of day period for post-saline day 1 (left, 16-24 hr) or post-LPS day 1 (right, 88-96 hr), corresponding to period 16-24 hr following injection. **K.** Daily total food intake during 24 hr period following either saline injection (left, 0-24 hr) or LPS injection (right, 72-96 hr). **L.** Daily total energy expenditure during 24 hr period following either saline injection (left, 0-24 hr) or LPS injection (right, 72-96 hr). **M.** Energy in/energy out ratio during period 8 hr period following either saline injection (left, 0-8 hr) or LPS injection (right, 72-80 hr). Ratio was calculated by dividing food intake (kcal; shown in **A**) by energy expenditure (kcal; shown in **E**). **N.** Energy in/energy out ratio during 24 hr period following either saline injection (left, 0-24 hr) or LPS injection (right, 72-96 hr). Ratio was calculated by dividing food intake (kcal; shown in **K**) by energy expenditure (kcal; shown in **L**). Data represented as mean ± SEM (**A-N**); *p<0.05, **p<0.01, ****p<0.0001. Paired two-sided t-test (**A-N**).

**Figure S18. Inflammatory gene mRNA expression following saline or LPS.** Related to Figure 5. **A.** Absolute body weight at injection time. **B.** Relative body weight at injection time. **C.** Relative body weight 3 hr post-injection. **D.** Cumulative food intake 3hr post-injection. **E.** ARC *Tnf* expression. **F.** ARC *Ccl2* expression. **G.** ARC *Il1b* expression. **H.** ARC *Ifng* expression. **I.** VTA *Tnf* expression. **J.** VTA *Ccl2* expression. **K.** VTA *Il1b* expression. **L.** VTA *Ifng* expression. **M.** NAc *Tnf* expression. **N.** NAc *Ccl2* expression. **O.** NAc *Il1b* expression. **P.** NAc *Ifng* expression. Data represented as mean ± SEM (**A-P**); *p<0.05, **p<0.01, ***p<0.001, ****p<0.0001. Unpaired two-tailed t-test (**A-P**).

**Figure S19. Additional metabolic and behavioural data.** Related to Figure 6. **A.** Absolute body weight at injection time. **B.** Relative body weight at injection time. **C.** Trial histogram from saline (24 hr) behaviour session. **D.** Trial histogram from LPS (24 hr) behaviour session. **E.** Trial histogram from LPS (48 hr) behaviour session. **G.** Feed efficiency. **H.** Ratio of forced travel frequency/total trials. **I.** Proportion of forced travels of total travels. **J.** Active patch discrimination. **K.** Mean harvest level at travel by patch type from saline (24 hr) session. **L.** Mean harvest level at travel by patch type from LPS (24 hr) session. **M.** Mean harvest level at travel by patch type from LPS (48 hr) session. N. Mean travel choice probability by patch type from saline (24 hr) session. **O.** Mean travel choice probability by patch type from LPS (24 hr) session. **P.** Mean travel choice probability by patch type from LPS (48 hr) session. **Q.** Mean time to travel choice by patch type from saline (24 hr) session. **R.** Mean time to travel choice by patch type from LPS (24 hr) session. **S.** Mean time to travel choice by patch type from LPS (48 hr) session. **T.** Change in trials and mean harvest level at travel from LPS (24 hr) session. **U.** Cumulative distribution of travels (18µl patch). **V.** Cumulative distribution of travels (12µl patch). **W.** Cumulative distribution of travels (6µl patch). Data represented as mean ± SEM (**A-B, G-M, Q-S**), histogram (**C-E**), mean (**N-P**), individual points (**T**), or cumulative distribution (**U-W**); *p<0.05. Black lines and shaded areas represent model and 95% confidence interval, respectively, from simple linear regression (**T**). Paired two-tailed t-test (**G-H**), Friedman test (**J**), Friedman test with Dunn's multiple comparisons (**K, M**), mixed effects model (REML) with Geisser-Greenhouse correction and Tukey's multiple comparisons (**L, R, S**), one-way ANOVA with Geisser-Greenhouse correction and Tukey's multiple comparisons (**Q**), and Kolmogorov-Smirnov test (**U-W**). Output of two-tailed Pearson's correlation shown in top right (**T**).

**Figure S20. Mean latency data sorted by patch type.** Related to Figure 6. **A.** Mean latency to first touch response by patch type from saline (24 hr) session. **B.** Mean latency to first touch response by patch type from LPS (24 hr) session. **C.** Mean latency to first touch response by patch type from LPS (48 hr) session. **D.** Mean latency to earn decision by patch type from saline (24 hr) session. **E.** Mean latency to earn decision by patch type from LPS (24 hr) session. **F.** Mean latency to earn decision by patch type from LPS (48 hr) session. **G.** Mean effort time by patch type from saline (24 hr) session. **H.** Mean effort time by patch type from LPS (24 hr) session. **I.** Mean effort time by patch type from LPS (48 hr) session. **J.** Mean latency to collect reward by patch type from saline (24 hr) session. **K.** Mean latency to collect reward by patch type from LPS (24 hr) session. **L.** Mean latency to collect reward by patch type from LPS (48 hr) session. **M.** Mean latency to complete travel by patch type from saline (24 hr) session. **N.** Mean latency to complete travel by patch type from LPS (24 hr) session. **O.** Mean latency to complete travel by patch type from LPS (48 hr) session. Data represented as mean (**A-O**). Hatched areas indicate no data.

**Figure S21. Mean latency data sorted by session.** Related to Figure 6. **A.** Mean latency to first touch response by session (18µl patch). **B.** Mean latency to first touch response by session (12µl patch). **C.** Mean latency to first touch response by session (6µl patch). **D.** Mean latency to earn decision by session (18µl patch). **E.** Mean latency to earn decision by session (12µl patch). **F.** Mean latency to earn decision by session (6µl patch).

**G.** Mean effort time by session (18µl patch). **H.** Mean effort time by session (12µl patch). **I.** Mean effort time by session (6µl patch). **J.** Mean latency to collect reward by session (18µl patch). **K.** Mean latency to collect reward by session (12µl patch). **L.** Mean latency to complete travel by session (18µl patch). **M.** Mean latency to complete travel by session (12µl patch). Data represented as mean (**A-M**). Hatched areas indicate no data.

#### Supplementary Figure 1

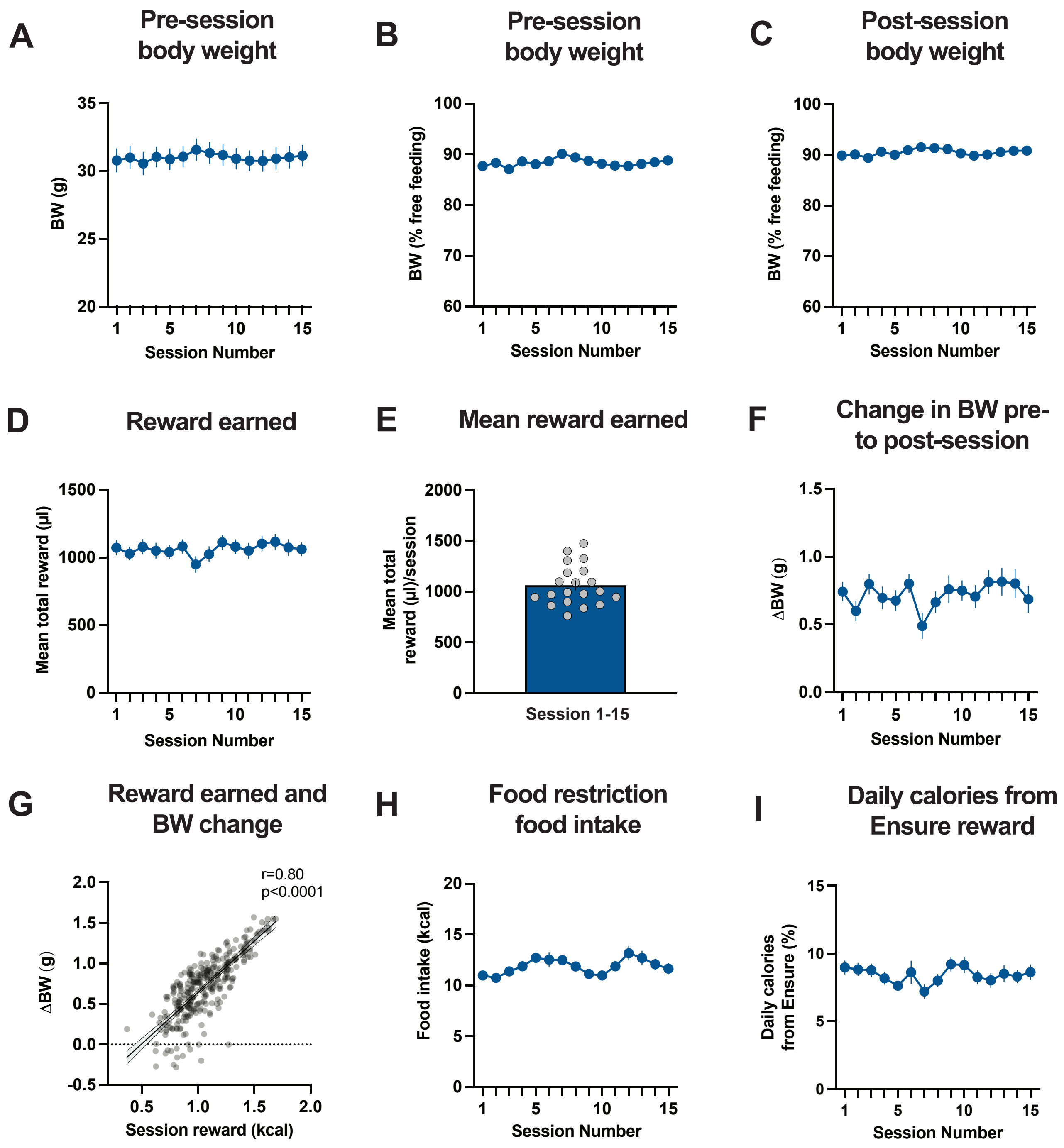

### Supplementary Figure 2

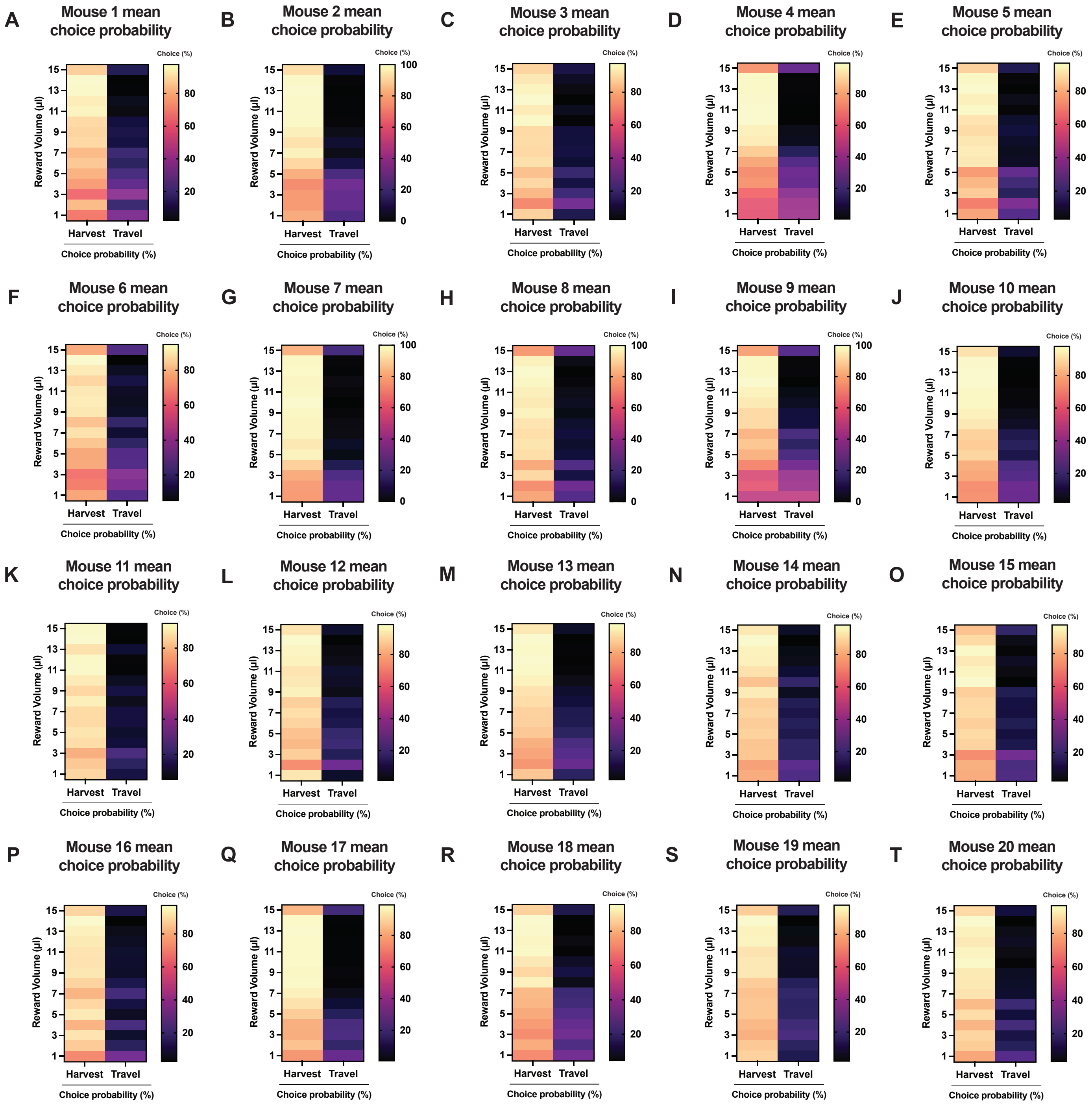

### Supplementary Figure 3

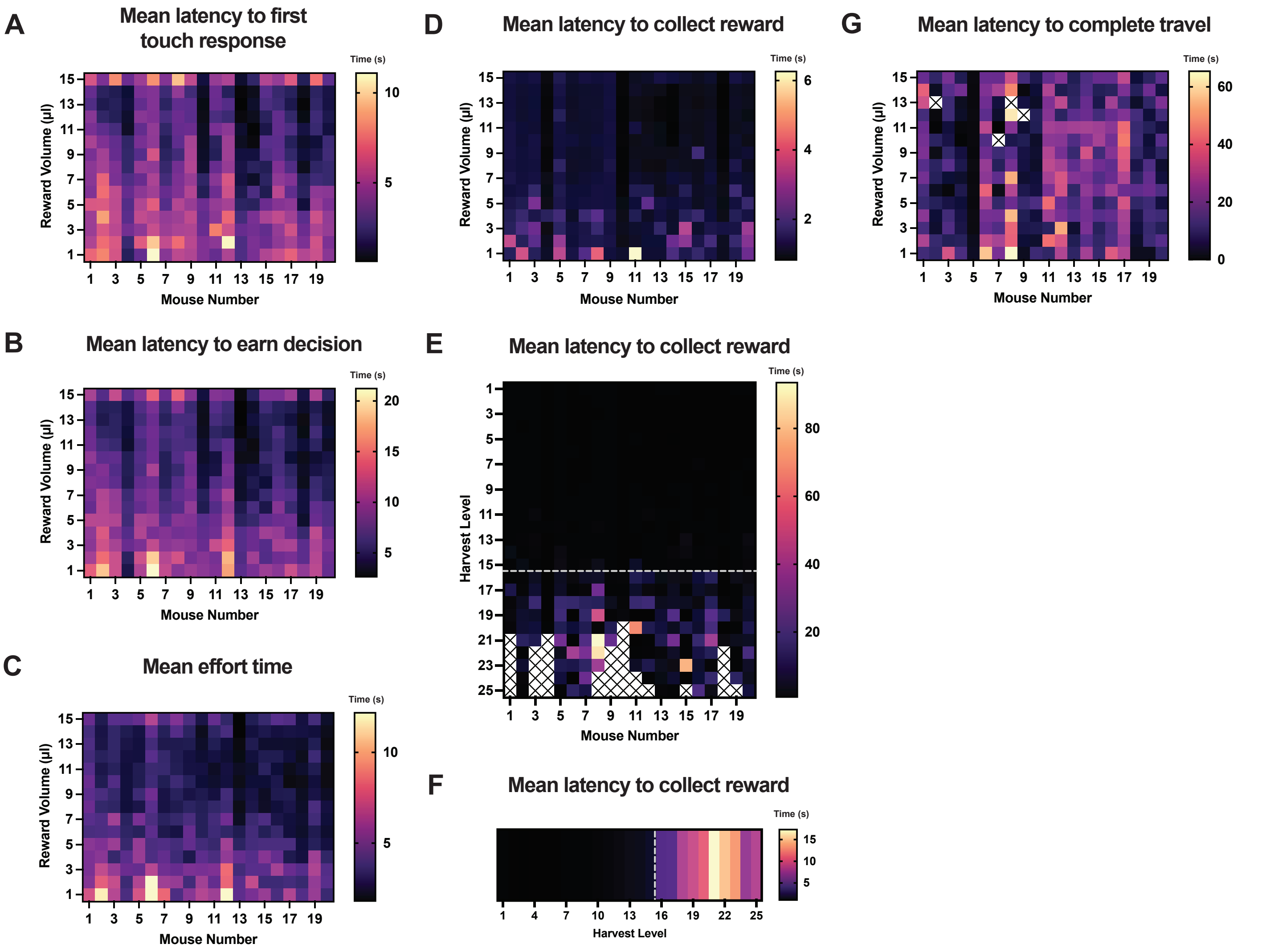

### Supplementary Figure 4

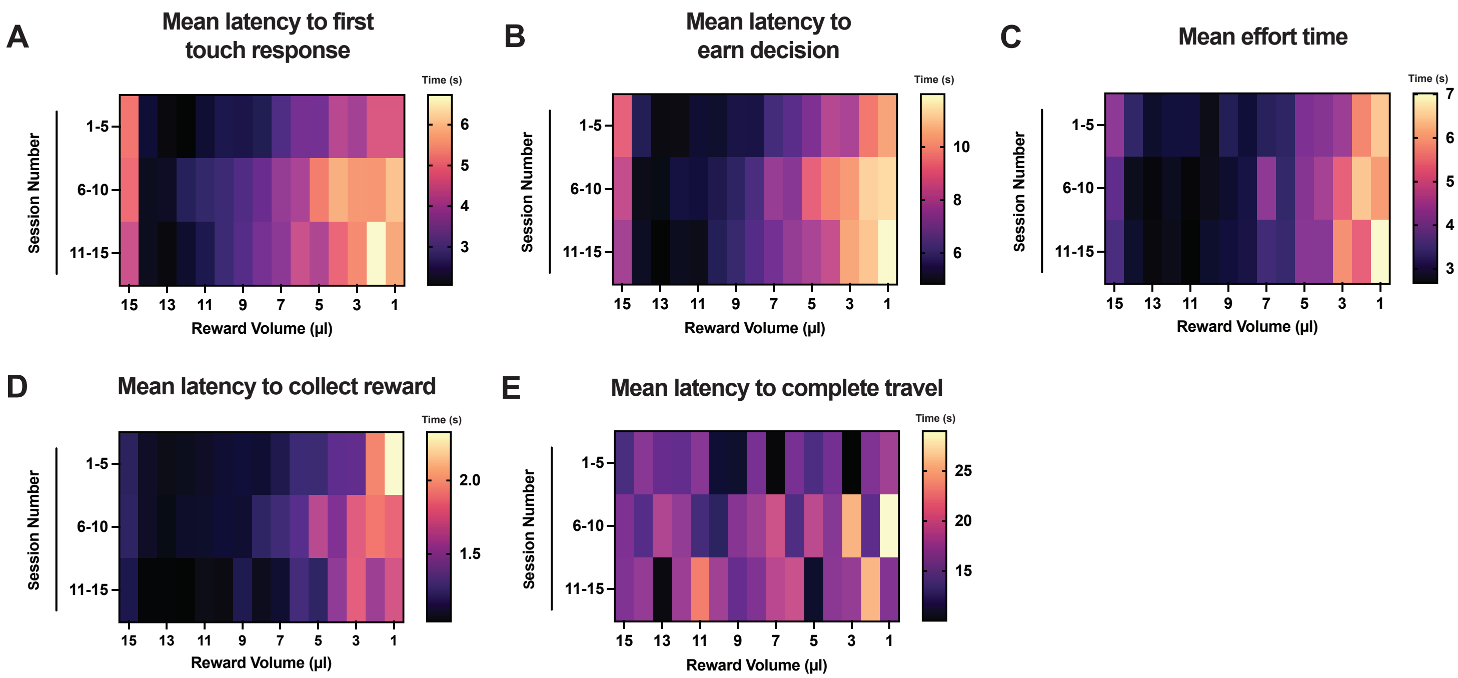

### Supplementary Figure 5

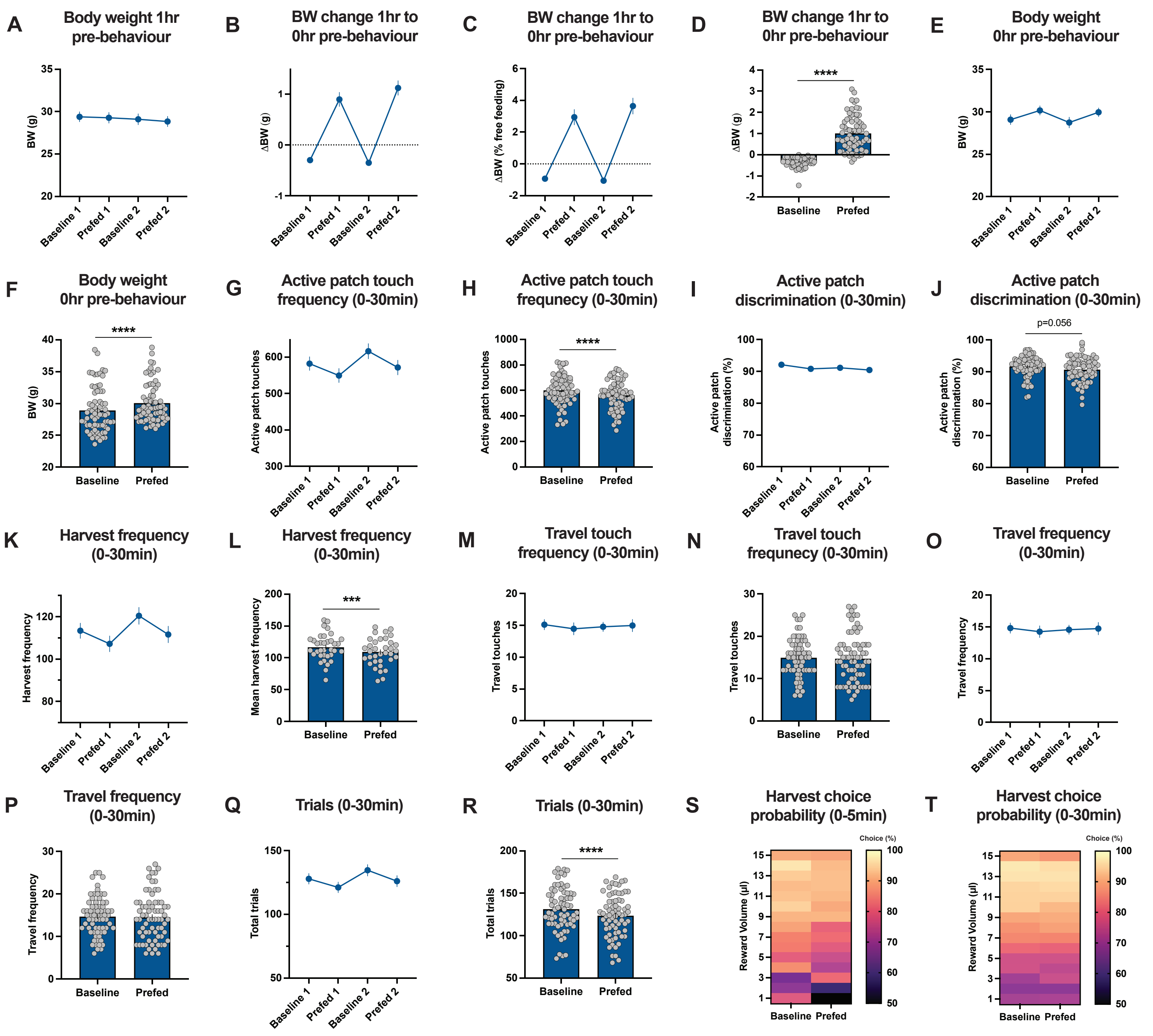

### Supplementary Figure 6

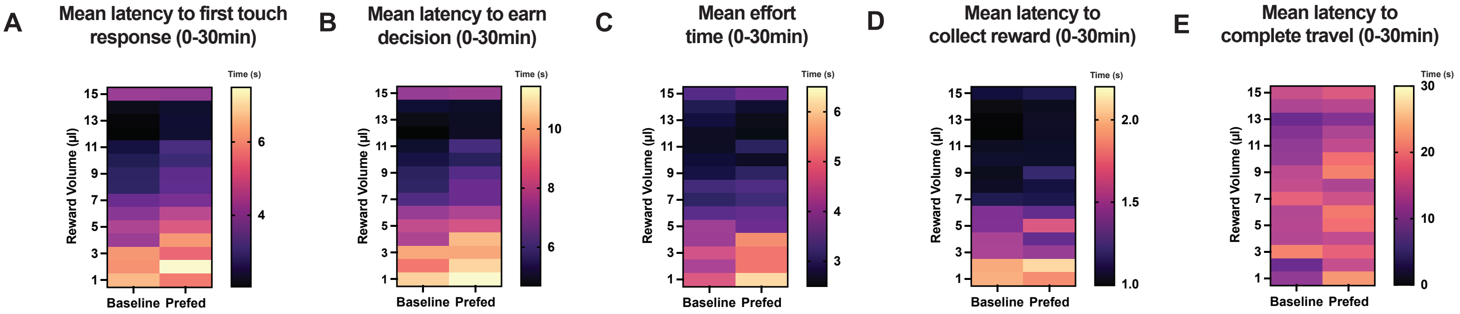

### Supplementary Figure 7

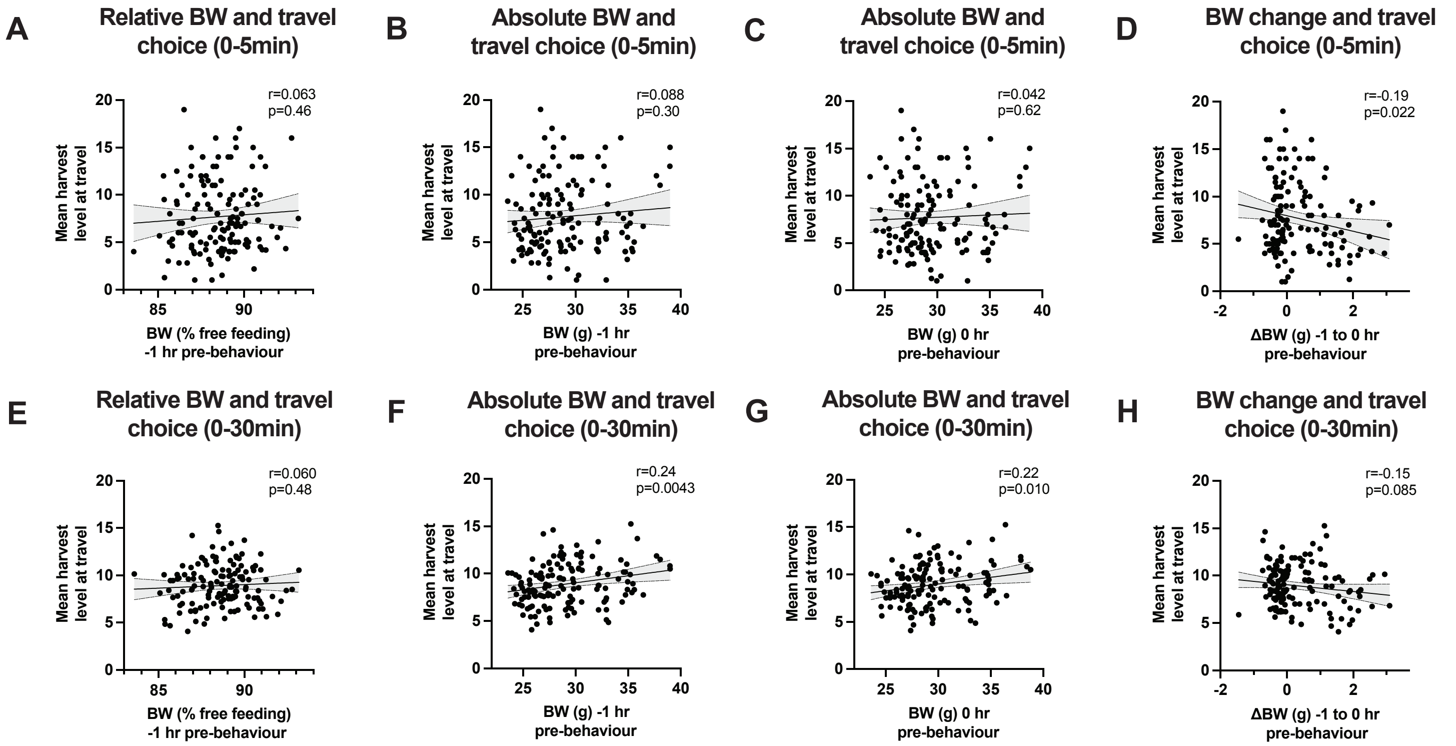

#### Supplementary Figure 8

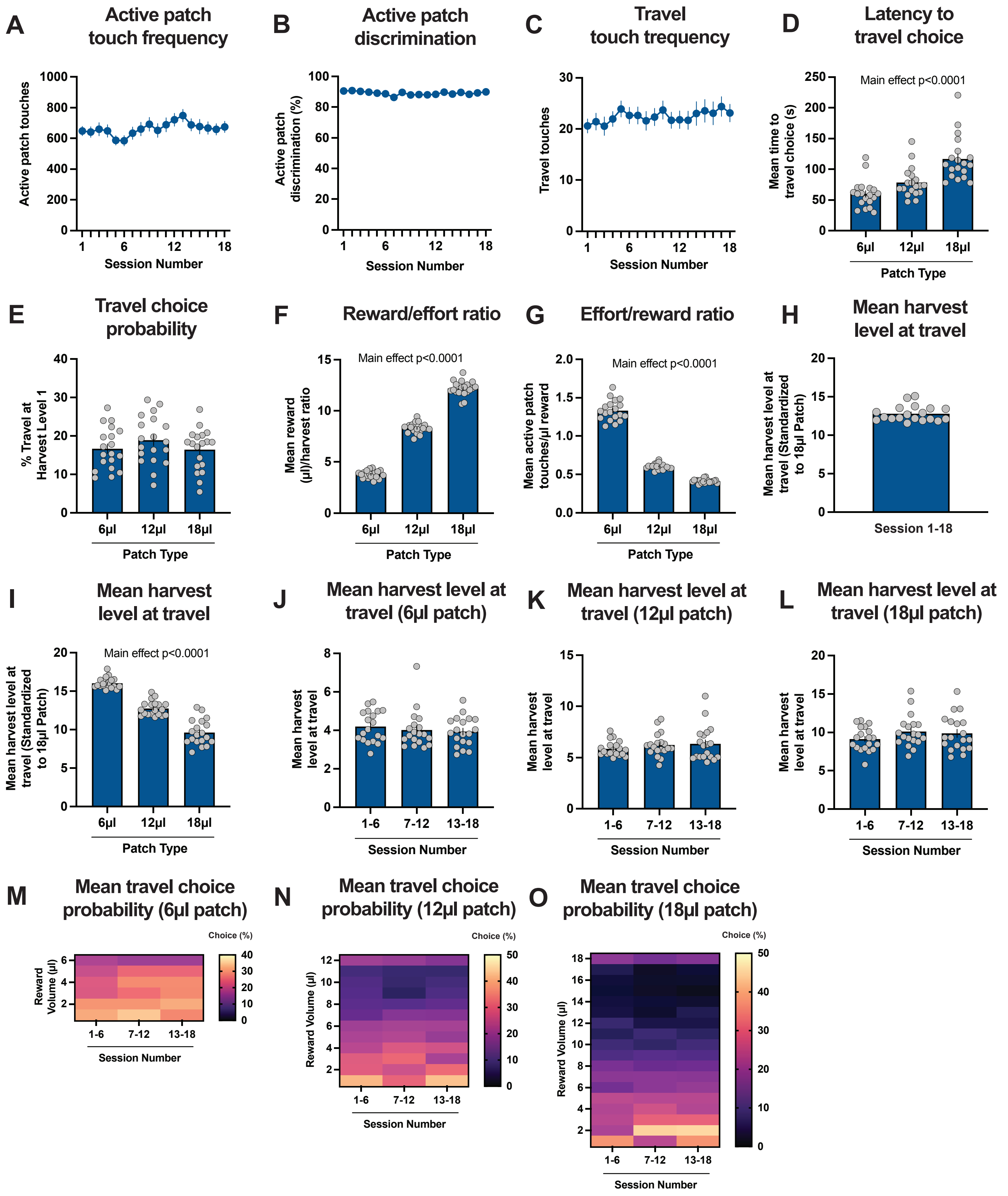

### Supplementary Figure 9

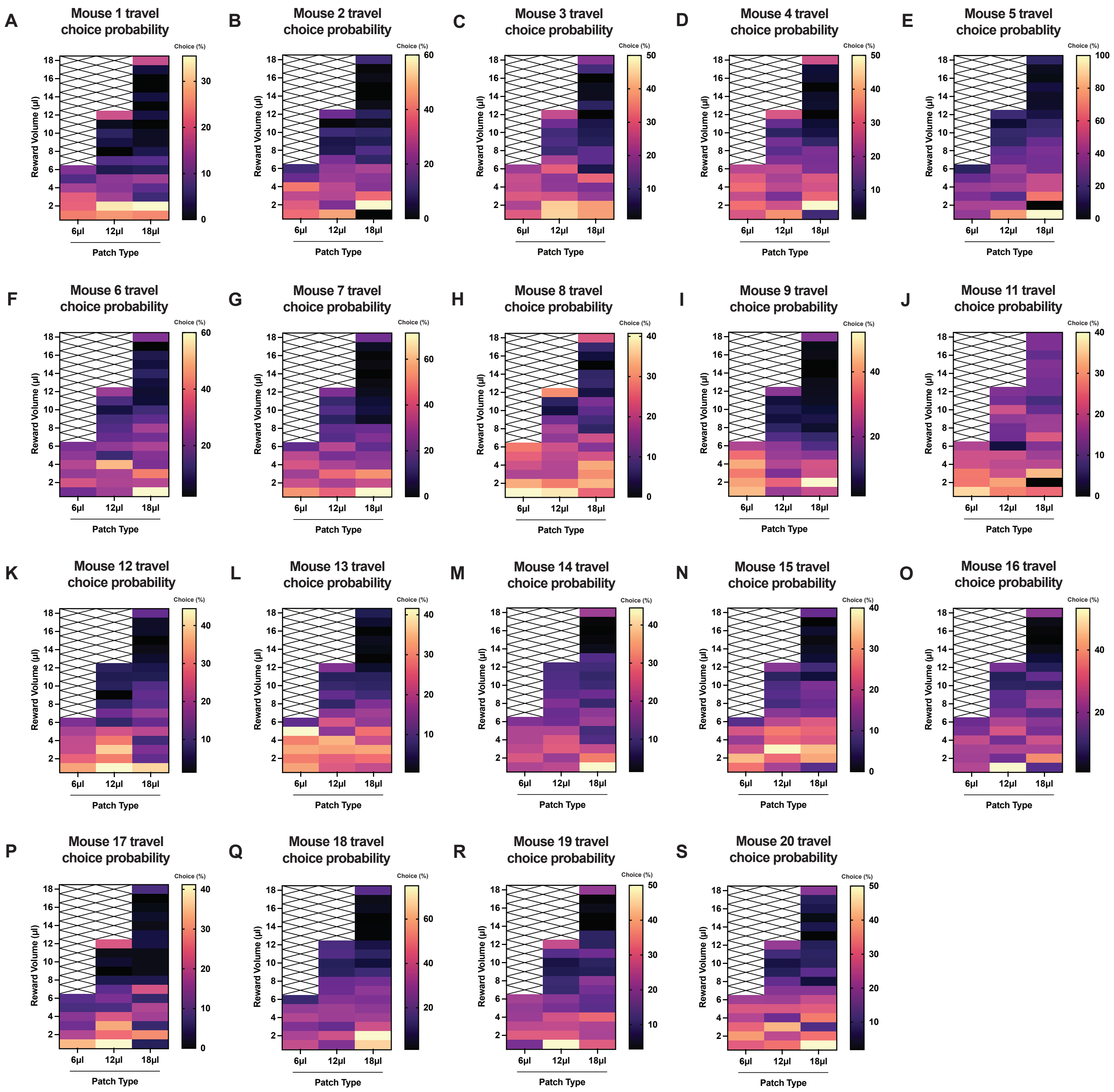

### Supplementary Figure 10

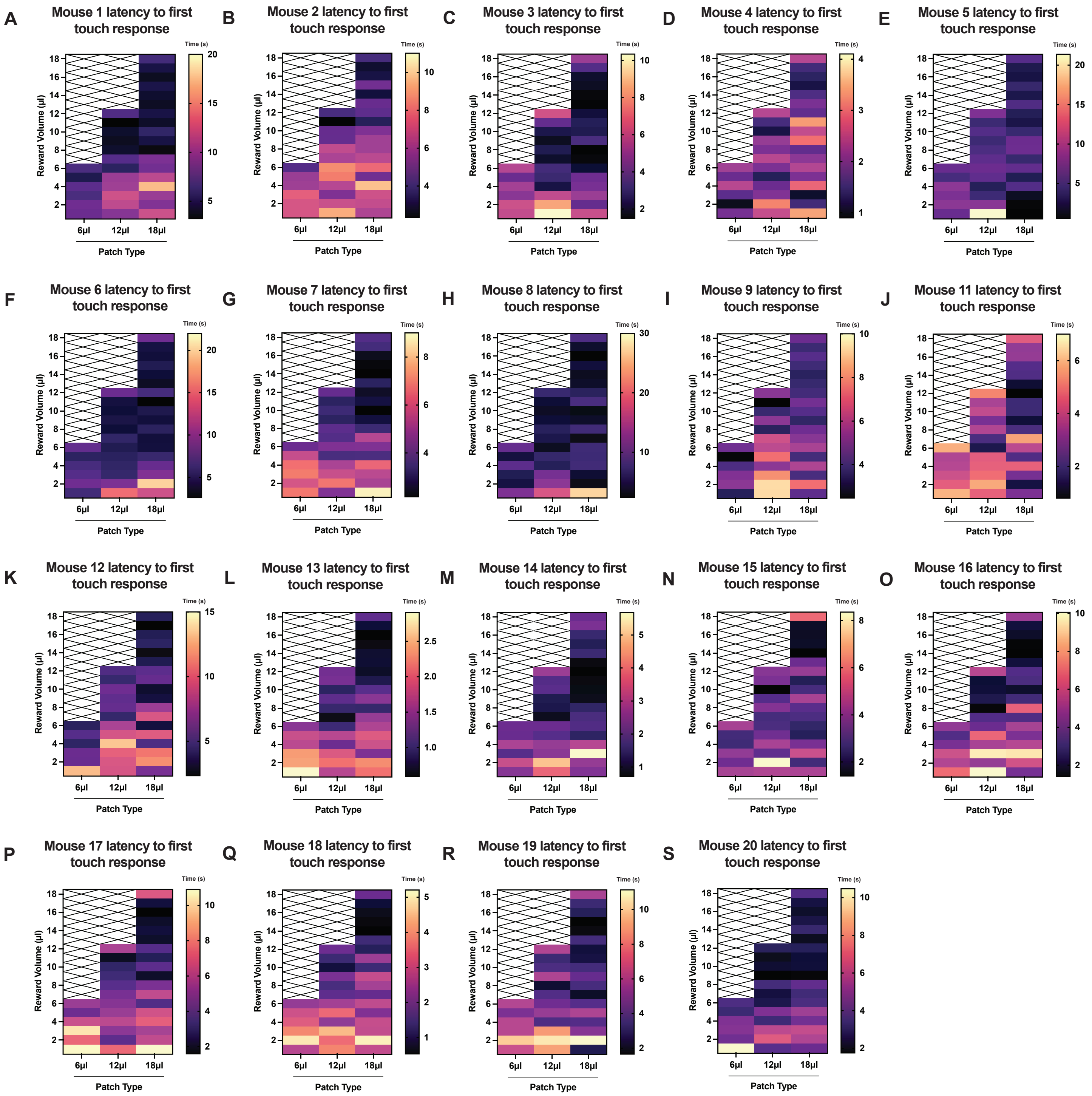

### Supplementary Figure 11

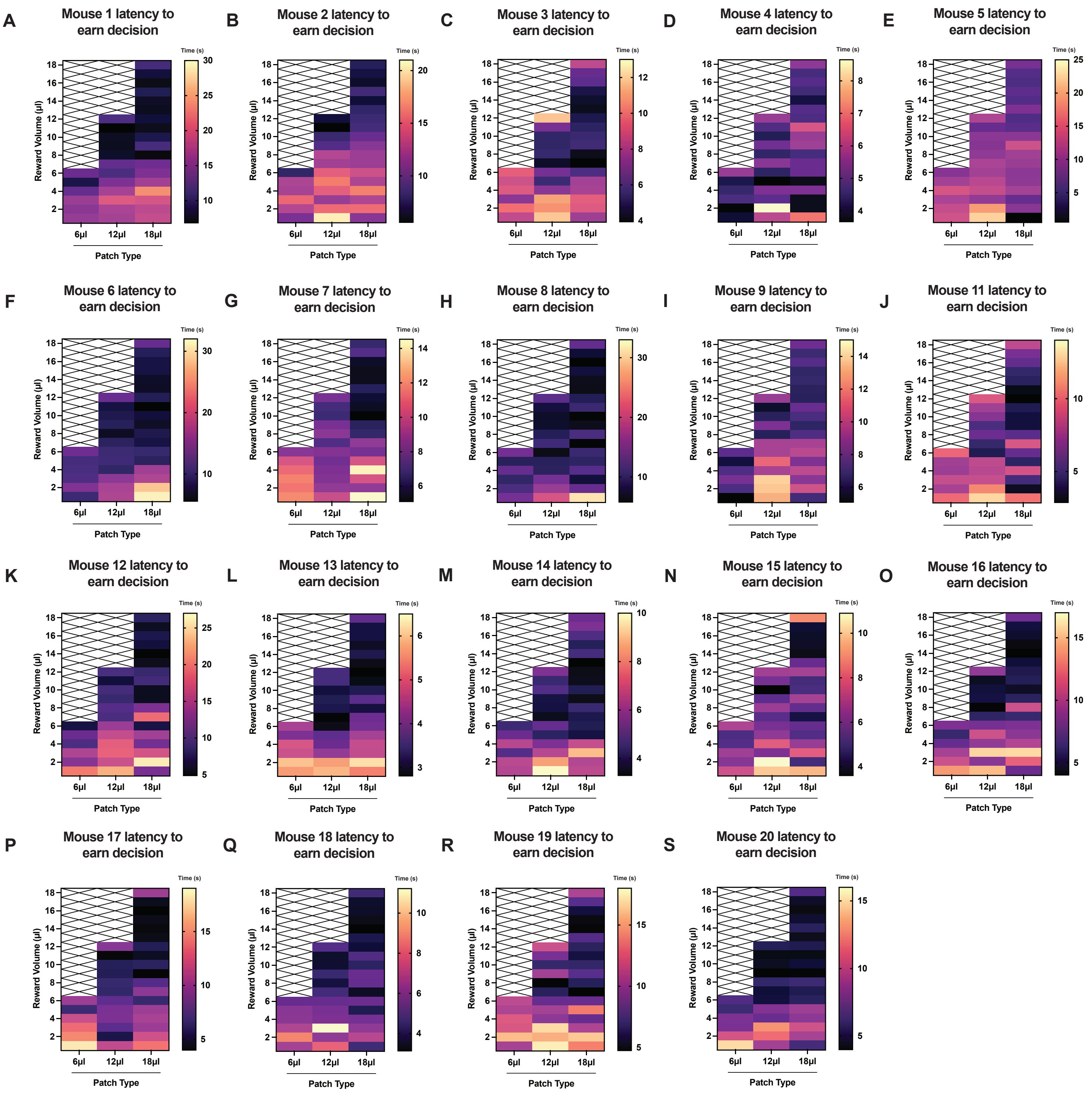

### Supplementary Figure 12

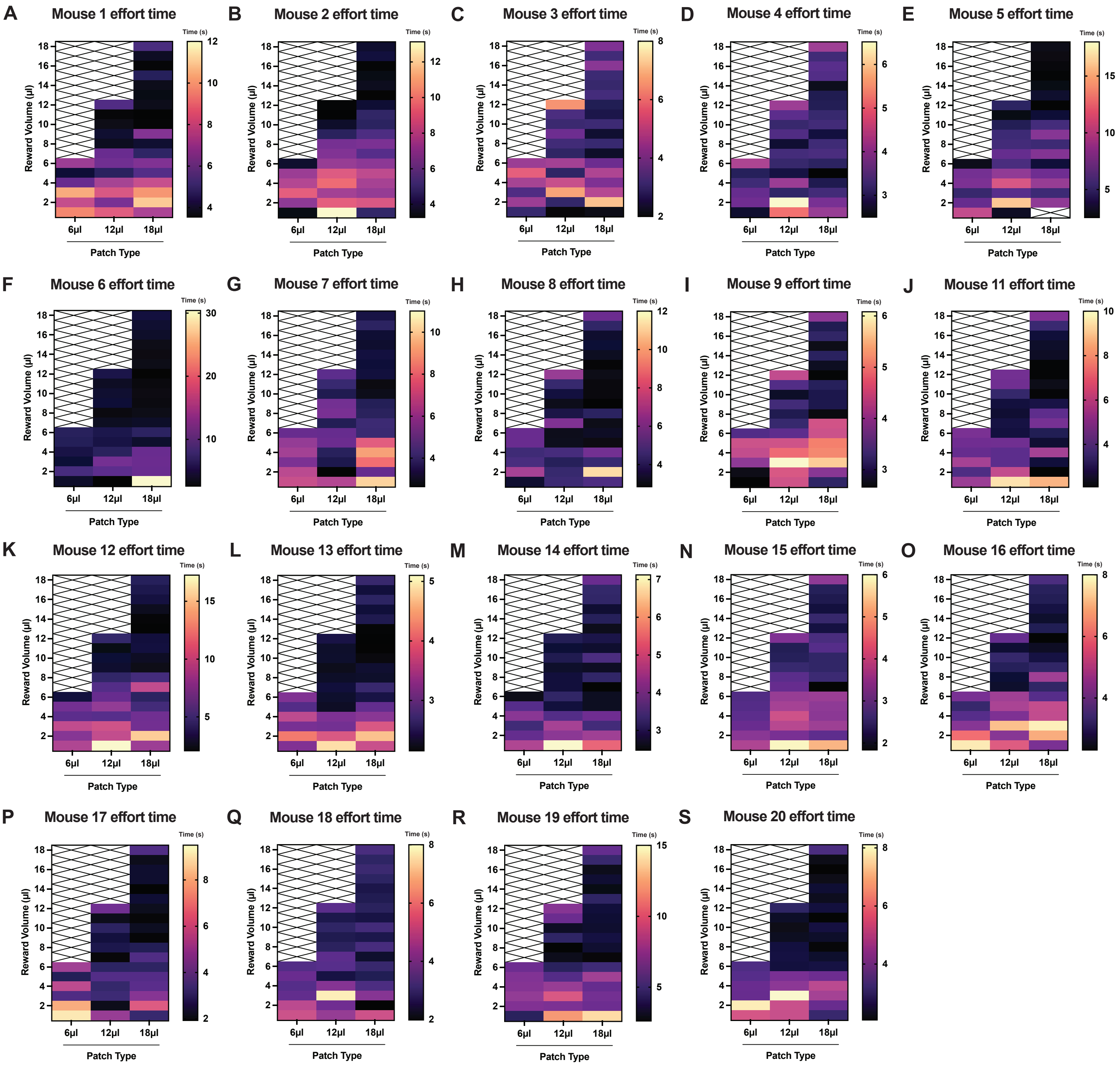

### Supplementary Figure 13

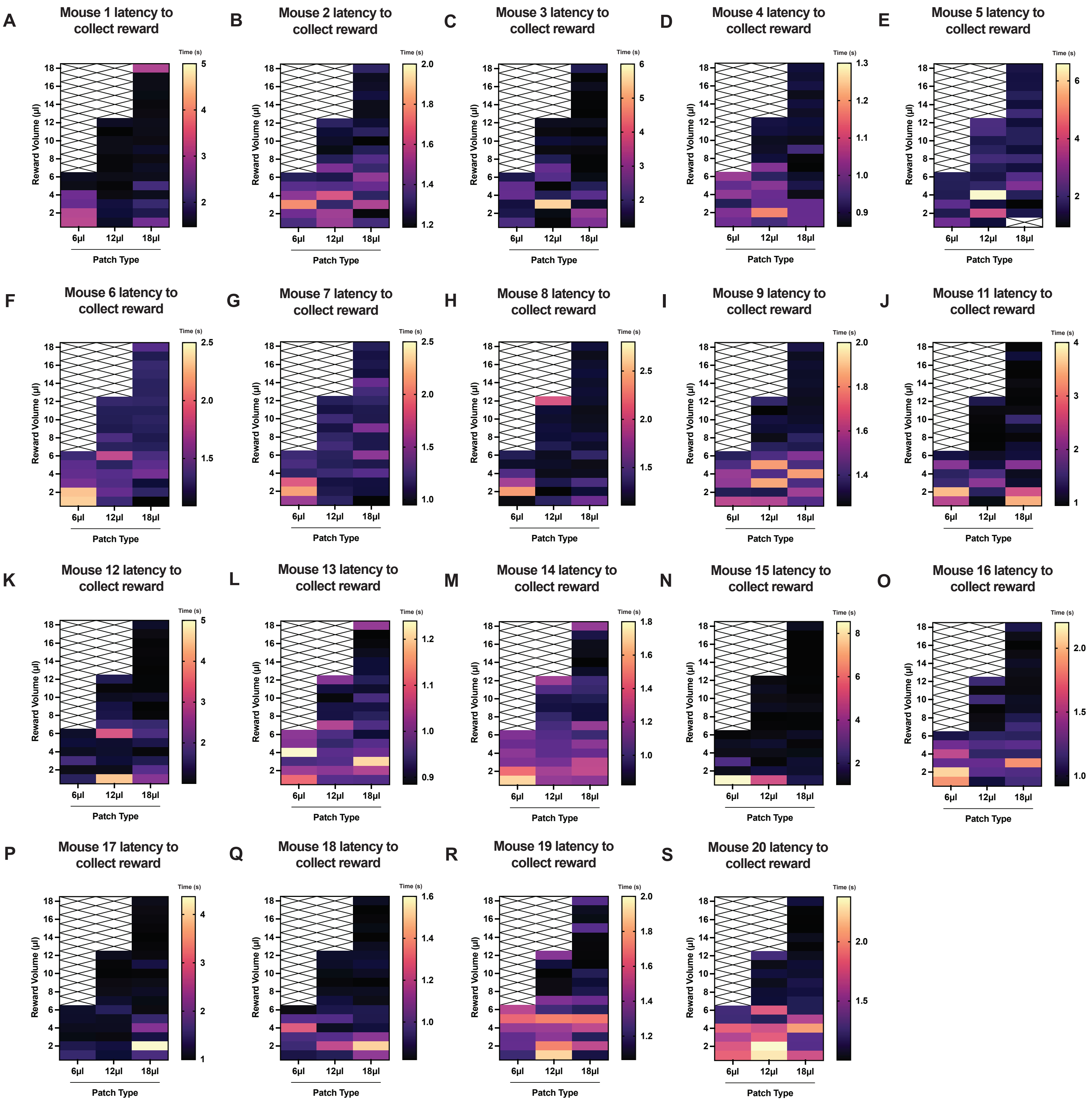

### Supplementary Figure 14

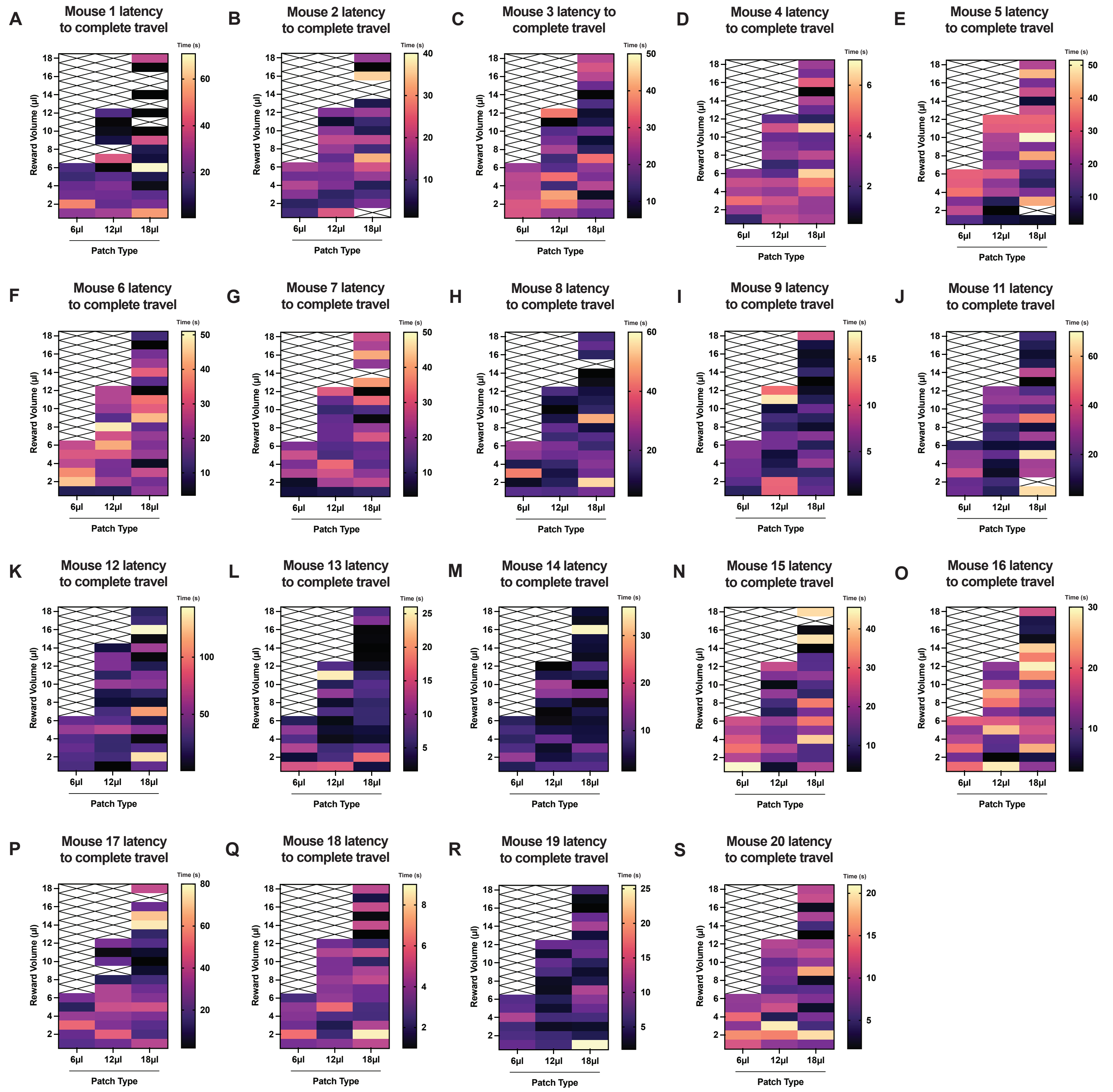

### Supplementary Figure 15

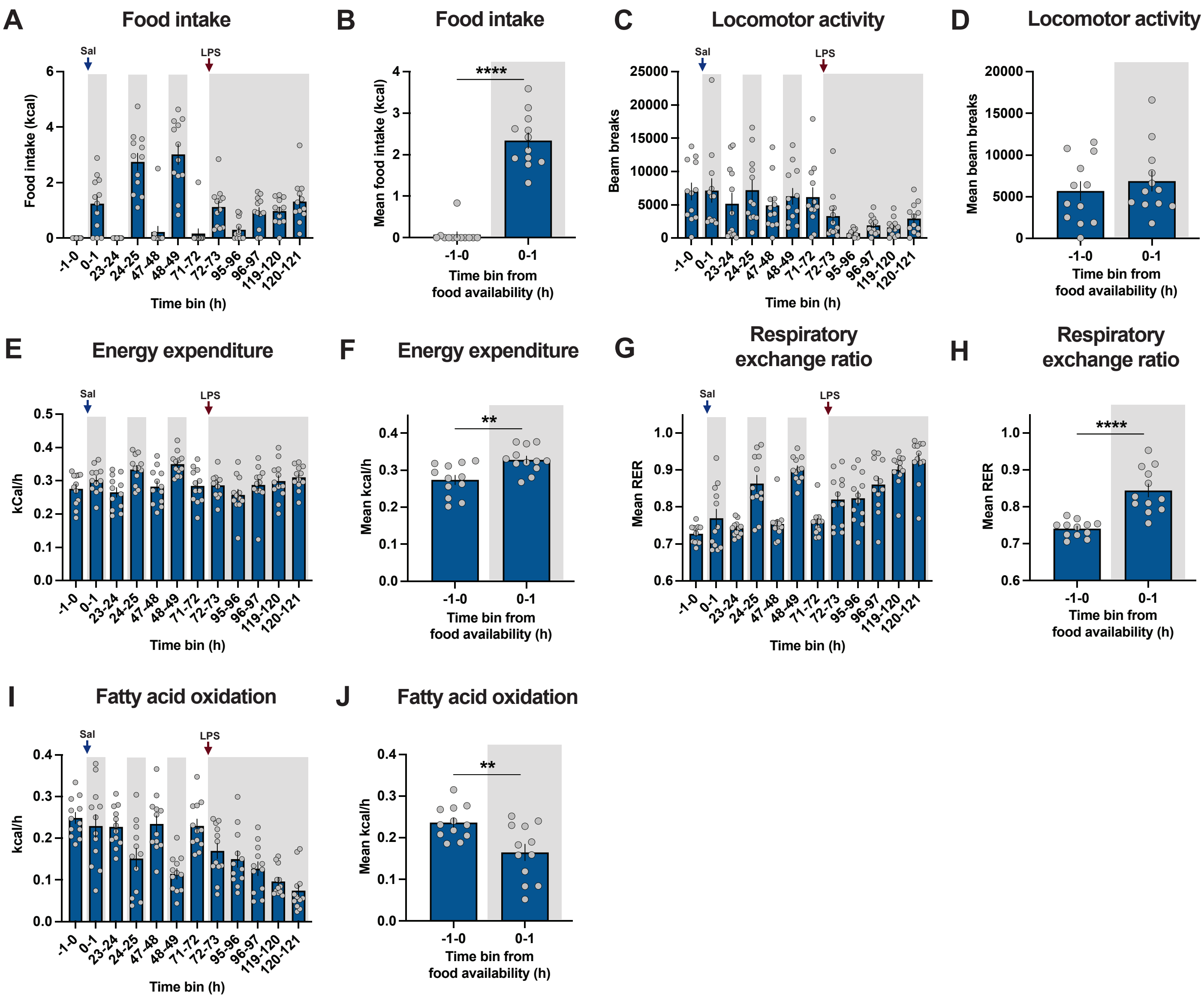

### Supplementary Figure 16

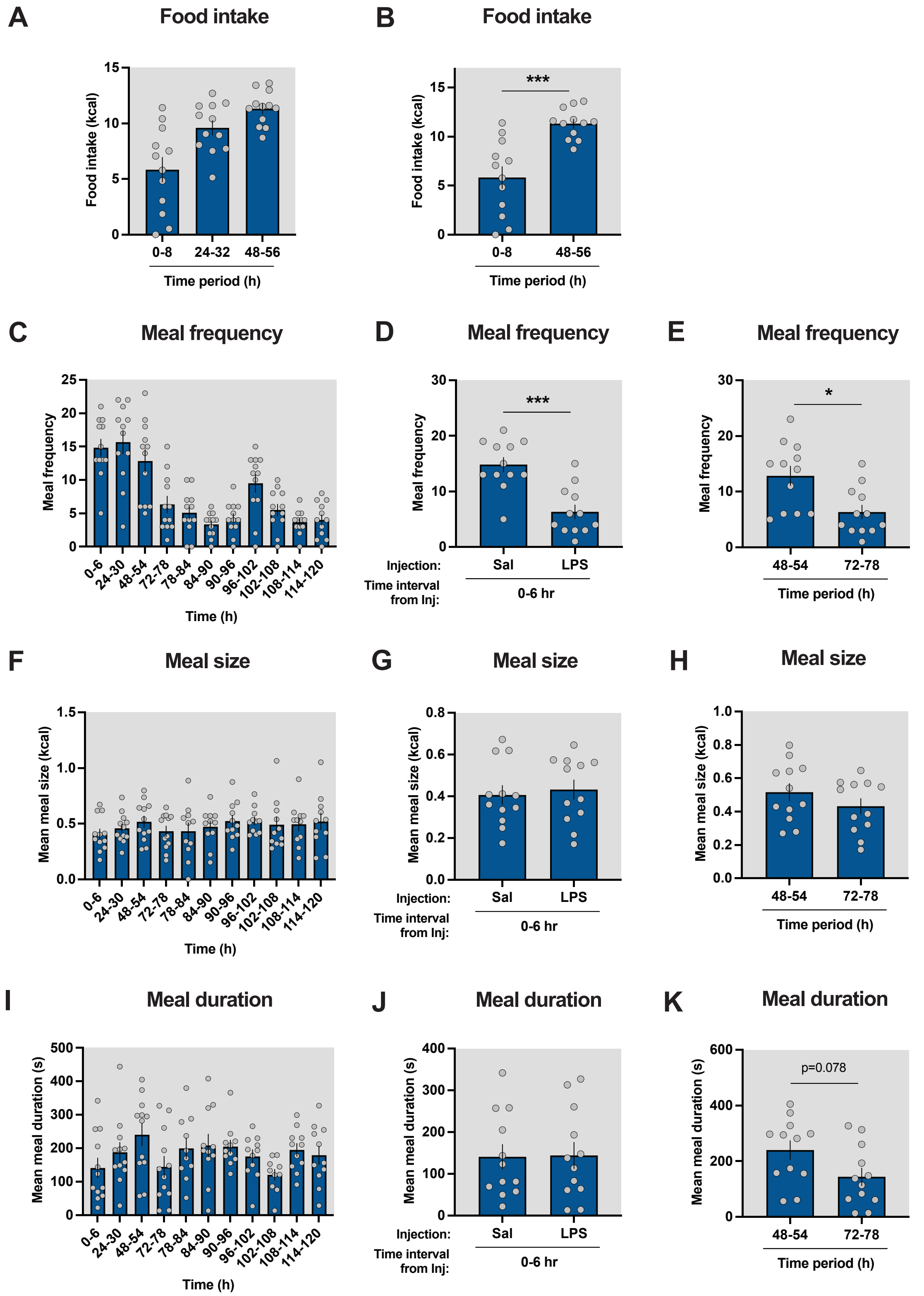

### Supplementary Figure 17

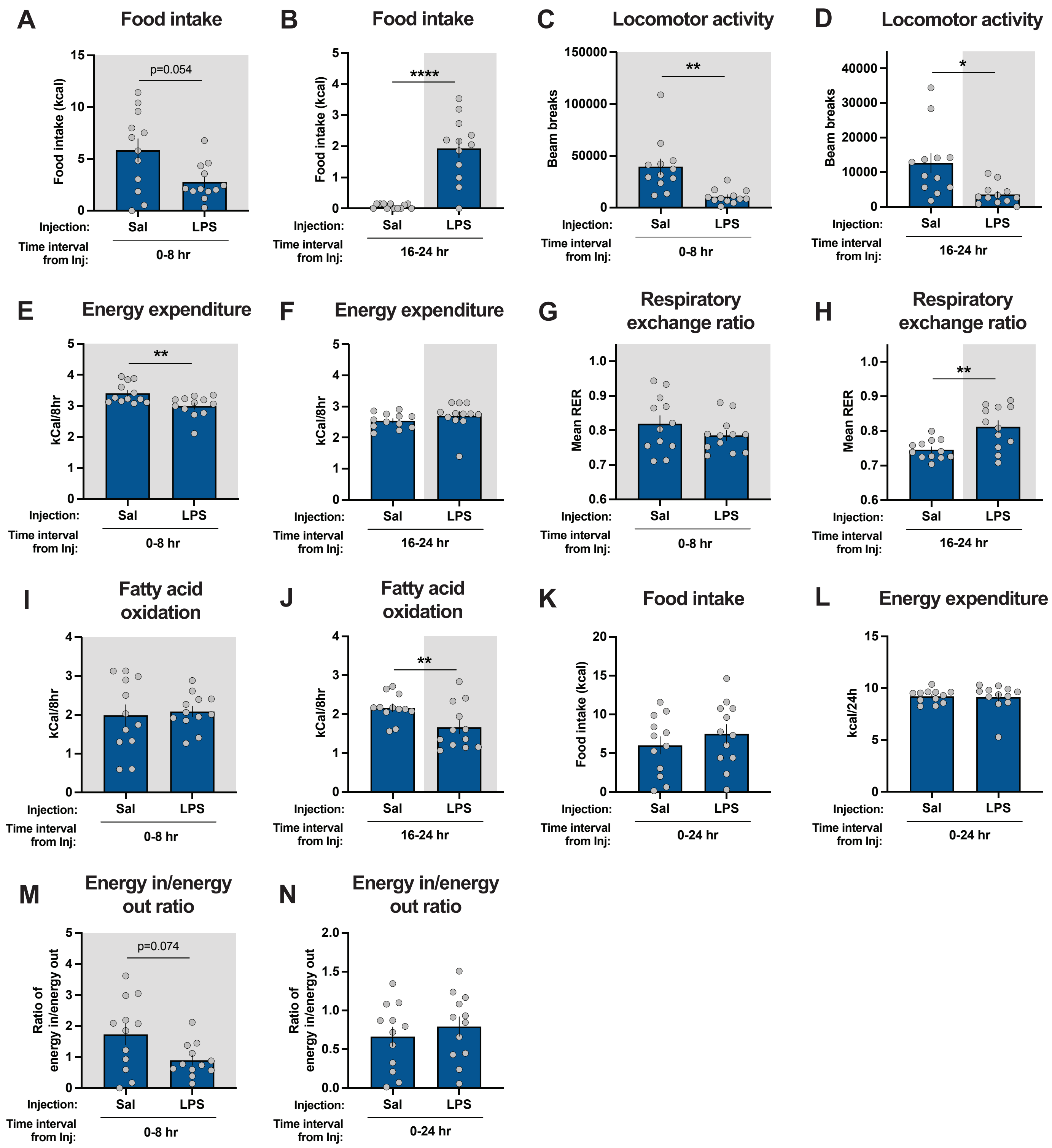

#### Supplementary Figure 18

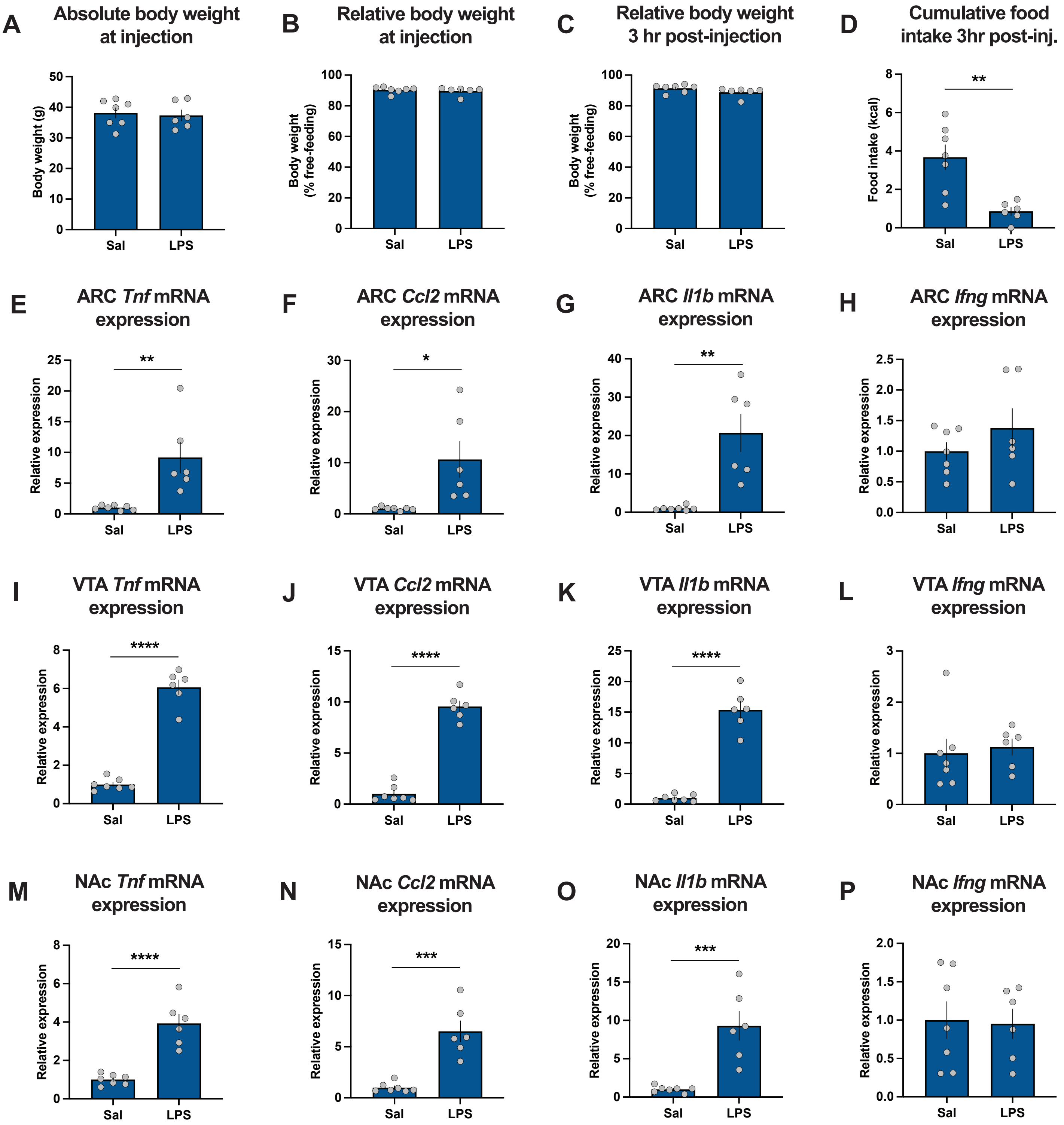

### Supplementary Figure 19

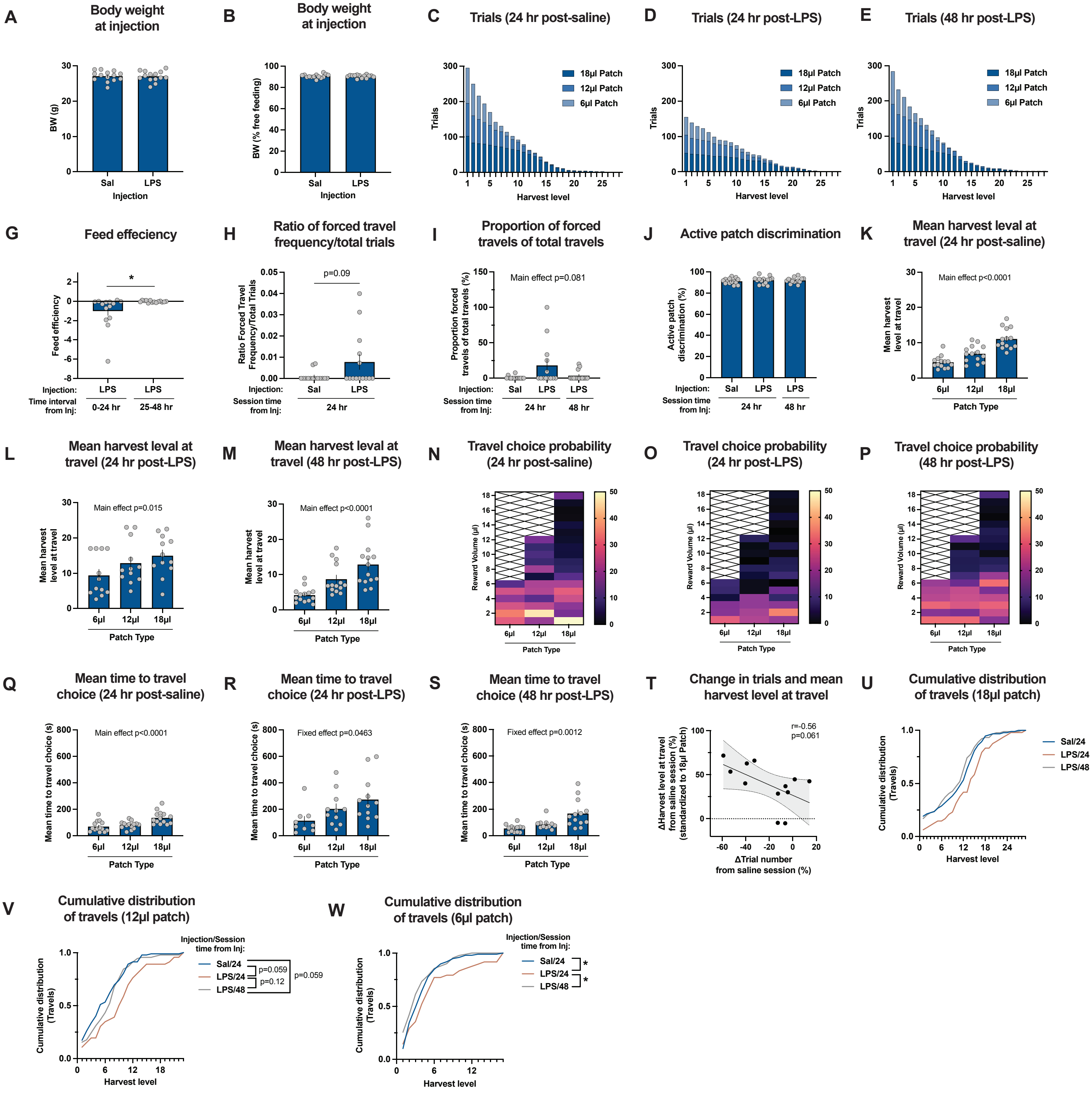

### Supplementary Figure 20

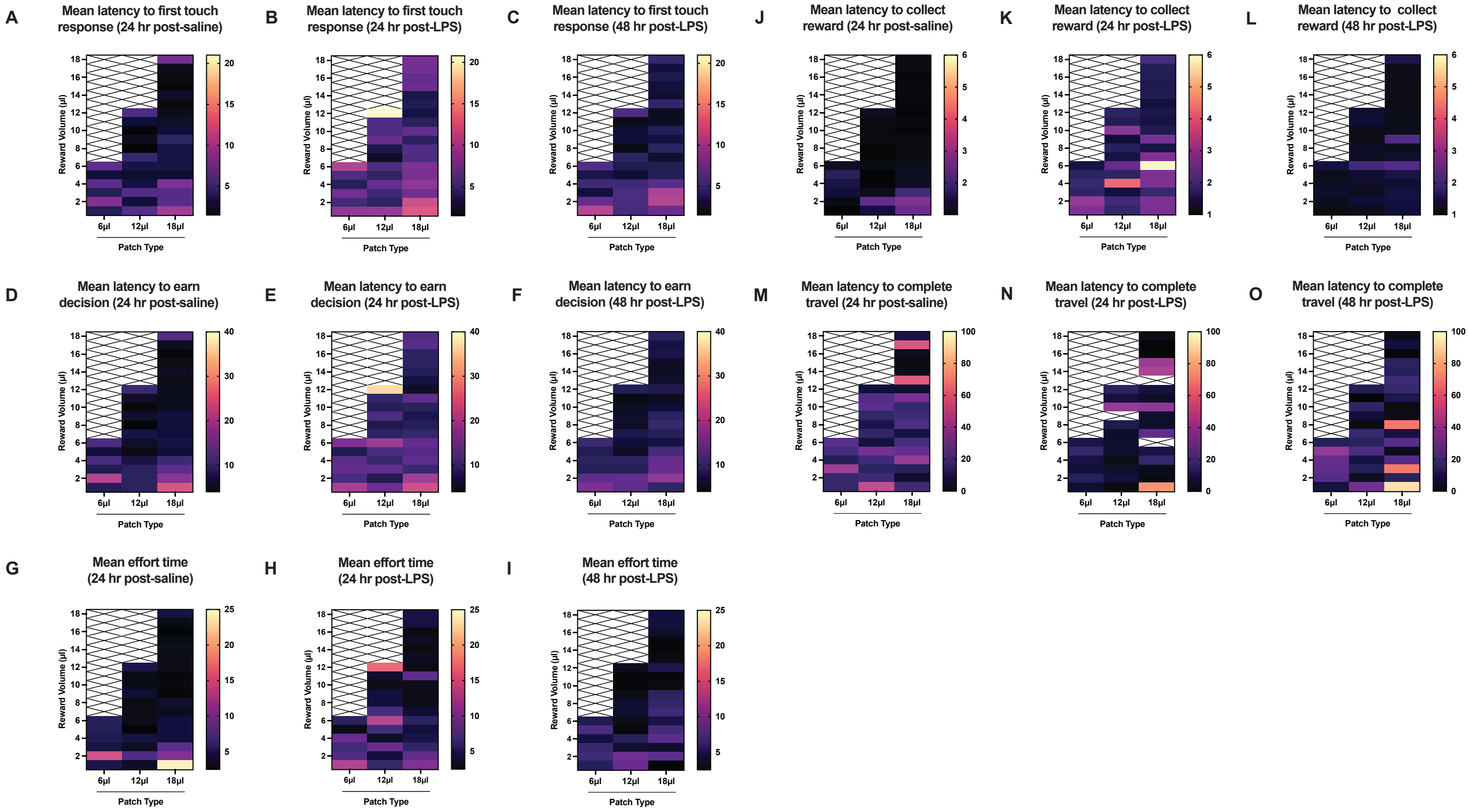

### Supplementary Figure 21

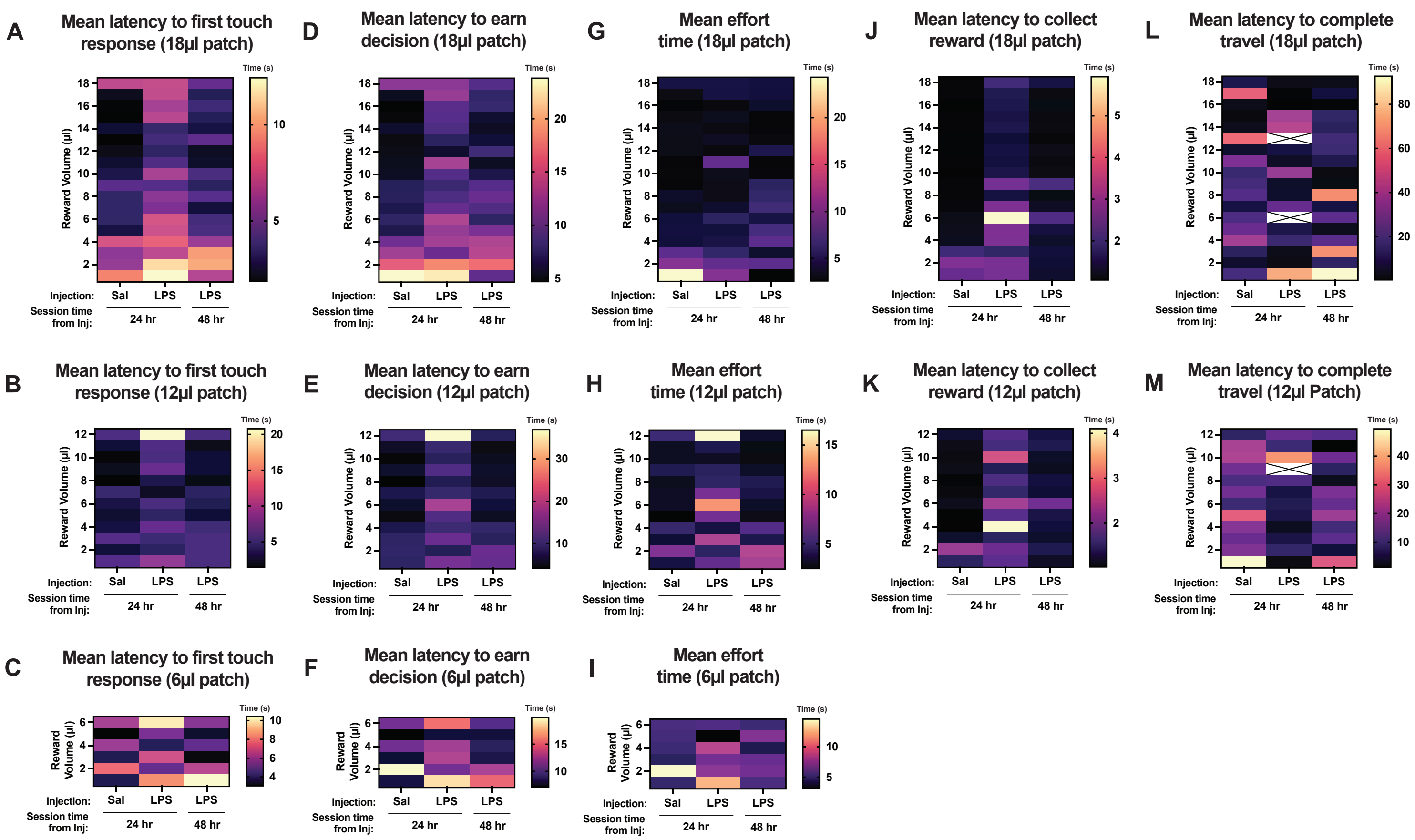

#### Supplementary Tables

**Supplementary Table 1.** List of Primer Sequences

| Gene | Forward sequence | Reverse sequence |
|---|---|---|
| *Cyclophilin* | GCTTTTCGCCGCTTGCTGCA | TGCAAACAGCTCGAAGGAGACGC |
| *Tnf* | GATCGGTCCCCAAAGGGATG | GCTCCTCCACTTGGTGGTTT |
| *Ccl2* | ATTGGGATCATCTTGCTGGT | CCTGCTGTTCACAGTTGCC |
| *Il1b* | TGCCACCTTTTGACAGTGATG | TGATGTGCTGCTGCGAGATT |
| *Ifng* | AAGTTTGAGGTCAACAACCCAC | AATCTCTTCCCCACCCCGAA |

**Supplementary Table 2.** Figure 2 Statistics

| Figure Panel | Response Variable | n | Statistical analysis | Result |
|---|---|---|---|---|
| Figure 2B | Mean harvest frequency | 20 mice (mean of 5 session blocks) | Two-tailed paired t-test | t=1.762, df=19, P=0.0942 |
| Figure 2D | Mean travel frequency | 20 mice (mean of 5 session blocks) | Two-tailed paired t-test | t=2.301, df=19, P=0.0329 |
| Figure 2F | Mean total reward per session | 20 mice (mean of 5 session blocks) | Two-tailed paired t-test | t=1.126, df=19, P=0.2743 |
| Figure 2H | Mean harvest level at travel | 20 mice (mean of 5 session blocks) | Two-tailed paired t-test | t=2.381, df=19, P=0.0279 |
| Figure 2I | Mean coefficient of variation | 5 sessions per block | Two-tailed paired t-test | t=4.234, df=4, P=0.0133 |
| Figure 2K | Reward/effort ratio | 20 mice (mean of 5 session blocks) | Two-tailed paired t-test | t=4.545, df=19, P=0.0002 |
| Figure 2L | Effort/reward ratio | 20 mice (mean of 5 session blocks) | Two-tailed paired t-test | t=4.609, df=19, P=0.0002 |

**Supplementary Table 3.** Figure 3 Statistics

| Figure Panel | Response Variable | n | Statistical analysis | Result |
|---|---|---|---|---|
| Figure 3C | Mean food intake (kcal) from -1h to 0h Pre-behaviour | 35 mice, 2 sessions/condition per mouse | Two-tailed paired t-test | t=15.84, df=69, P<0.0001 |
| Figure 3E | Mean body weight change from -1h to 0h Pre-behaviour | 35 mice, 2 sessions/condition per mouse | Two-tailed paired t-test | t=12.24, df=69, P<0.0001 |
| Figure 3G | Mean body weight (% free-feeding) at 0h Pre-behaviour | 35 mice, 2 sessions/condition per mouse | Two-tailed paired t-test | t=11.65, df=69, P<0.0001 |
| Figure 3J | Mean BW change (g) 0h Pre-Behaviour to Post-behaviour | 35 mice, 2 sessions/condition per mouse | Two-tailed paired t-test | t=4.967, df=69, P<0.0001 |
| Figure 3L | Mean harvest level at travel (0-5 min behaviour) | 35 mice, 2 sessions/condition per mouse | Two-tailed paired t-test | t=0.5130, df=69, P=0.6096 |
| Figure 3M | Mean harvest frequency (0-5 min behaviour) | 35 mice, 2 sessions/condition per mouse | Two-tailed paired t-test | t=3.073, df=34, P=0.0042 |
| Figure 3N | Mean reward earned per session (μl; 0-5 min behaviour) | 35 mice, 2 sessions/condition per mouse | Two-tailed paired t-test | t=2.668, df=69, P=0.0095 |
| Figure 3O | Reward/effort ratio (0-5 min behaviour) | 35 mice, 2 sessions/condition per mouse | Two-tailed paired t-test | t=2.533, df=69, P=0.0136 |

| | | | | |
|---|---|---|---|---|
| Figure 3P | Effort/reward ratio (0-5 min behaviour) | 35 mice, 2 sessions/condition per mouse | Two-tailed paired t-test | t=2.388, df=69, P=0.0197 |
| Figure 3Q | Mean harvest level at travel (0-5 min behaviour) | 35 mice, 2 sessions/condition per mouse (139 XY pairs) | Simple linear regression (mean harvest level at travel as a function of BW at 0h pre-behaviour) | Y=-0.2569*X+30.75, $R^2$=0.03366; slope significantly non-zero, $F_{(1,137)}$=4.771, P=0.0306 |
| Figure 3Q | Mean harvest level at travel (0-5 min behaviour) | 35 mice, 2 sessions/condition per mouse (139 XY pairs) | Pearson's correlation (mean harvest level as a function of BW at 0h pre-behaviour) | r= -0.1835, 95% CI: -0.3396 to -0.01749, $R^2$=0.03366, P=0.0306 |
| Figure 3R | Mean harvest level at travel (0-5 min behaviour) | 35 mice, 2 pre-fed sessions per mouse (70 XY pairs) | Simple linear regression (mean harvest level at travel as a function of pre-feeding food intake (kcal)) | Y=-0.8387*X+10.07, $R^2$=0.1268, slope significantly non-zero, $F_{(1,68)}$=9.872, P=0.0025 |
| Figure 3R | Mean harvest level at travel (0-5 min behaviour) | 35 mice, 2 pre-fed sessions per mouse (70 XY pairs) | Pearson's correlation (mean harvest level at travel as a function of pre-feeding food intake (kcal)) | r=-0.3560, 95% CI: -0.5454 to -0.1321, $R^2$=0.1268, P=0.0025 |
| Figure 3S | Mean harvest level at travel (0-5 min behaviour) | 35 mice, 2 sessions/condition per mouse (139 XY pairs) | Simple linear regression (mean harvest level at travel as a function of change in BW from -1 to 0h pre-behaviour) | Y=-0.2578*X+8.001, $R^2$=0.04129, slope significantly non-zero, $F_{(1,137)}$=5..900, P=0.0164 |
| Figure 3S | Mean harvest level at travel (0-5 min behaviour) | 35 mice, 2 sessions/condition per mouse (139 XY pairs) | Pearson's correlation (mean harvest level at travel as a function of change in BW from -1 to 0h pre-behaviour) | r=-0.2032, 95% CI: -0.3576 to -0.03798, $R^2$=0.04129, P=0.0164 |
| Figure 3U | Mean harvest level at travel (0-30 min behaviour) | 35 mice, 2 sessions/condition per mouse | Two-tailed paired t-test | t=0.1487, df=69, P=0.8822 |
| Figure 3V | Mean harvest frequency | 35 mice, 2 sessions/condition per mouse | Two-tailed paired t-test | t=4.523, df=69, P<0.0001 |
| Figure 3W | Mean total reward earned per session | 35 mice, 2 sessions/condition per mouse | Two-tailed paired t-test | t=4.006, df=69, P=0.0002 |
| Figure 3X | Mean reward/effort ratio | 35 mice, 2 sessions/condition per mouse | Two-tailed paired t-test | t=0.04286, df=69, P=0.9659 |
| Figure 3Y | Mean effort/reward ratio | 35 mice, 2 sessions/condition per mouse | Two-tailed paired t-test | t=0.1294, df=69, P=0.8974 |
| Figure 3Z | Mean harvest level at travel (0-30 min behaviour) | 35 mice, 2 sessions/condition per mouse (140 XY pairs) | Simple linear regression (mean harvest level at travel as a function | Y=-0.1056*X+18.39, $R^2$=0.01682, slope not significantly non-zero, $F_{(1,138)}$=2.361, P=0.1267 |

| | | | | |
|---|---|---|---|---|
| | | | of BW at 0h pre-behaviour) | |
| Figure 3Z | Mean harvest level at travel (0-30 min behaviour) | 35 mice, 2 sessions/condition per mouse (140 XY pairs) | Pearson's correlation (mean harvest level at travel as a function of BW at 0h pre-behaviour) | r=-0.1297, 95% CI: -0.2894 to 0.03700, $R^2$=0.01682, P=0.1267 |
| Figure 3AA | Mean harvest level at travel (0-30 min behaviour) | 35 mice, 2 pre-fed sessions per mouse (70 XY pairs) | Simple linear regression (mean harvest level at travel as a function of pre-feeding food intake (kcal)) | Y=-0.3705*X+10.02, $R^2$=-0.05899, slope significantly non-zero, $F_{(1,68)}$=4.262, P=0.0428 |
| Figure 3AA | Mean harvest level at travel (0-30 min behaviour) | 35 mice, 2 pre-fed sessions per mouse (70 XY pairs) | Pearson's correlation (mean harvest level at travel as a function of pre-feeding food intake (kcal)) | r=-0.2429, 95% CI: -0.4520 to -0.008373, $R^2$=-0.05899, P=0.0428 |
| Figure 3AB | Mean harvest level at travel (0-30 min behaviour) | 35 mice, 2 sessions/condition per mouse (140 XY pairs) | Simple linear regression (mean harvest level as a function of change in BW from -1 to 0h pre-behaviour) | Y=-0.1119*X+9.043, $R^2$=-0.02314, slope not significantly non-zero $F_{(1,138)}$=3.268, P=0.0728 |
| Figure 3AB | Mean harvest level at travel (0-30 min behaviour) | 35 mice, 2 sessions/condition per mouse (140 XY pairs) | Pearson's correlation (mean harvest level as a function of change in BW from -1 to 0h pre-behaviour) | r=-0.1521, 95% CI: -0.3102 to 0.01415, $R^2$=-0.02314, P=0.0728 |

**Supplementary Table 4.** Figure 4 Statistics

| **Figure Panel** | **Response Variable** | **n** | **Statistical analysis** | **Result** |
|---|---|---|---|---|
| Figure 4N | Mean harvest level at travel | 19 mice, 18 sessions/mouse | Repeated measures one-way ANOVA with Geisser-Greenhouse correction; Tukey's post-hoc multiple comparisons | $F_{(1.271,22.87)}$=265.7, P<0.0001, Geisser-Greenhouse's epsilon 0.6353 Tukey's post hoc comparisons: 6μl vs 12μl q=19.68, df=18, adjusted P<0.0001; 6μl vs 18μl q=24.90, df=18, adjusted P<0.0001; 12μl vs 18μl q=20.77, df=18, adjusted P<0.0001 |

**Supplementary Table 5.** Figure 5 Statistics

| Figure Panel | Response Variable | n | Statistical analysis | Result |
|---|---|---|---|---|
| Figure 5F | Food intake (kcal) at 48-56h or 72-80h | 12 mice (12 pairs) | Two-tailed paired t-test | t=12.37, df=11, P<0.0001 |
| Figure 5G | Food intake (kcal) at 64-72h or 88-96h | 12 mice (12 pairs) | Two-tailed paired t-test | t=6.114, df=11, P<0.0001 |
| Panel 5J | Locomotor activity (beam breaks) at 48-56h or 72-80h | 12 mice (12 pairs) | Two-tailed paired t-test | t=6.788, df=11, P<0.0001 |
| Panel 5K | Locomotor activity (beam breaks) at 64-72h or 88-96h | 12 mice (12 pairs) | Two-tailed paired t-test | t=4.481, df=11, P=0.0009 |
| Panel 5N | Energy expenditure (kcal/8hr) at 48-56h or 72-80h | 12 mice (12 pairs) | Two-tailed paired t-test | t=5.693, df=11, P=0.0001 |
| Panel 5O | Energy expenditure (kcal/8hr) at 64-72h or 88-96h | 12 mice (12 pairs) | Two-tailed paired t-test | t=0.7258, df=11, P=0.4831 |
| Panel 5P | Energy in/energy out ratio at 48-56h or 72-80h | 12 mice (12 pairs) | Two-tailed paired t-test | t=11.83, df=11, P<0.0001 |
| Panel 5Q | Food intake (kcal) at 48-72h or 72-96h | 12 mice (12 pairs) | Two-tailed paired t-test | t=3.648, df=11, P=0.0038 |
| Panel 5R | Energy expenditure (kcal/24hr) at 48-72h or 72-96h | 12 mice (12 pairs) | Two-tailed paired t-test | t=1.245, df=11, P=0.2391 |
| Panel 5S | Energy in/energy out ratio | 12 mice (12 pairs) | Two-tailed paired t-test | t=3.875, df=11, P=0.0026 |
| Panel 5V | Mean RER at 48-56h or 72-80h | 12 mice (12 pairs) | Two-tailed paired t-test | t=9.131, df=11, P<0.0001 |
| Panel 5W | Mean RER at 64-72h or 88-96h | 12 mice (12 pairs) | Two-tailed paired t-test | t=3.186, df=11, P=0.0087 |
| Panel 5Z | Mean FAO at 48-56h or 72-80h | 12 mice (12 pairs) | Two-tailed paired t-test | t=8.536, df=11, P<0.0001 |

**Supplementary Table 6.** Figure 6 Statistics

| Figure Panel | Response Variable | n | Statistical analysis | Result |
|---|---|---|---|---|
| Figure 6B | Cumulative food intake (kcal) at 8h post injection | 13 mice (13 pairs) | Two-tailed paired t-test | t=12.51, df=12, P<0.0001 |
| Figure 6B | Cumulative food intake (kcal) at 24h post injection | 14 mice (14 pairs) | Two-tailed paired t-test | t=4.811, df=13, P=0.0003 |
| Figure 6C | Proportion of available food left uneaten (%) at 8h post injection | 13 mice (13 pairs) | Two-tailed paired t-test | t=15.36, df=12, P<0.0001 |
| Figure 6C | Proportion of available food left uneaten (%) at 24h post injection | 14 mice (14 pairs) | Two-tailed paired t-test | t=6.542, df=13, P<0.0001 |
| Figure 6D | Body weight (% of free-feeding) at 8h post injection | 13 mice (13 pairs) | Two-tailed paired t-test | t=14.12, df=12, P<0.0001 |
| Figure 6D | Body weight (% of free-feeding) at 24h post injection | 14 mice (14 pairs) | Two-tailed paired t-test | t=3.086, df=13, P=0.0087 |

| | | | | |
|---|---|---|---|---|
| Figure 6E | Body weight change (% change from injection time) at 8h post injection | 13 mice (13 pairs) | Two-tailed paired t-test | t=13.10, df=12, P<0.0001 |
| Figure 6E | Body weight change (% change from injection time) at 8h post injection | 14 mice (14 pairs) | Two-tailed paired t-test | t=2.787, df=13, P=0.0154 |
| Figure 6F | Feed efficiency at 0-8h post injection | 13 mice (13 pairs) | Two-tailed paired t-test | t=2.647, df=12, P=0.0213 |
| Figure 6F | Feed efficiency at 0-24h post injection | 14 mice (14 pairs) | Two-tailed paired t-test | t=2.170, df=13, P=0.0491 |
| Figure 6G | Total trials completed per session | 14 mice, 3 sessions/mouse | Friedman's test with Dunn's multiple comparisons post hoc | Friedman statistic 13.20, approximate P=0.0014 Dunn's multiple comparisons: Sal 24 vs LPS 24 Rank sum diff. 16.50, adjusted P=0.0055, Sal 24 vs LPS 48 Rank sum diff. 0.000, adjusted P>0.9999, LPS 24 vs LPS 48 Rank sum diff. -16.50, adjusted P=0.0055 |
| Figure 6G | Change in total trials completed per session (%) from saline session | 14 mice, 3 sessions/mouse | Friedman's test with Dunn's multiple comparisons post hoc | Friedman statistic 13.20, approximate P=0.0014 Dunn's multiple comparisons: Sal 24 vs LPS 24 Rank sum diff. 16.50, adjusted P=0.0055, Sal 24 vs LPS 48 Rank sum diff. 0.000, adjusted P>0.9999, LPS 24 vs LPS 48 Rank sum diff. -16.50, adjusted P=0.0055 |
| Figure 6I | Harvest frequency per session | 14 mice, 3 sessions/mouse | Friedman's test with Dunn's multiple comparisons post hoc | Friedman statistic 7.000, approximate P=0.0302 Dunn's multiple comparisons: Sal 24 vs LPS 24 Rank sum diff. 13.00, adjusted P=0.0421, Sal 24 vs LPS 48 Rank sum diff. 2.000, adjusted P>0.9999, LPS 24 vs LPS 48 Rank sum diff. - |

| | | | | |
|---|---|---|---|---|
| | | | | 11.00, adjusted P=0.1129 |
| Figure 6J | Travel frequency per session (including forced travels) | 14 mice, 3 sessions/mouse | Repeated measures one-way ANOVA with Geisser-Greenhouse correction, and Tukey's multiple comparisons post hoc | $F_{(1.909,24.81)}$=25.12, P<0.0001, Geisser-Greenhouse's epsilon 0.9544<br>Tukey's multiple comparisons:<br>Sal 24 vs LPS 24 q=9.505, df=13, adjusted P<0.0001<br>Sal 24 vs LPS 48 q=0.6644, df=13, adjusted P=0.8865<br>LPS 24 vs LPS 48 q=6.617, df=13, adjusted P=0.0012 |
| Figure 6K | Mean harvest level at travel (6μl patch) | 14 mice, 3 sessions/mouse | Mixed-effects model (REML) with Geisser-Greenhouse correction, and Tukey's multiple comparisons post hoc | $F_{(1.181,14.17)}$=9.729, P=0.0056, Geisser-Greenhouse's epsilon 0.5905<br>Tukey's multiple comparisons:<br>Sal 24 vs LPS 24 q=4.978, df=11, adjusted P=0.0122<br>Sal 24 vs LPS 48 q=0.5574, df=13, P=0.9128<br>LPS 24 vs LPS 48 q=4.194, df=11, adjusted P=0.0318 |
| Figure 6L | Mean harvest level at travel (12μl patch) | 14 mice, 3 sessions/mouse | Mixed-effects model (REML) with Geisser-Greenhouse correction, and Tukey's multiple comparisons post hoc | $F_{(1.530,18.36)}$=10.31, P=0.0019, Geisser-Greenhouse's epsilon 0.7651<br>Tukey's multiple comparisons:<br>Sal 24 vs LPS 24 q=5.725, df=11, adjusted P=0.0050<br>Sal 24 vs LPS 48 q=2.600, df=13, P=0.1959<br>LPS 24 vs LPS 48 q=3.451, df=11, adjusted P=0.0776 |
| Figure 6M | Mean harvest level at travel (18μl patch) | 14 mice, 3 sessions/mouse | Mixed-effects model (REML) with Geisser-Greenhouse correction, and Tukey's multiple comparisons post hoc | $F_{(1.896,22.75)}$=3.638, P=0.0447, Geisser-Greenhouse's epsilon 0.9479<br>Tukey's multiple comparisons:<br>Sal 24 vs LPS 24 q=3.401, df=11, adjusted P=0.0822<br>Sal 24 vs LPS 48 q=1.737, df=13, P=0.4584<br>LPS 24 vs LPS 48 q=1.694, df=11, adjusted P=0.4784 |

| | | | | |
|---|---|---|---|---|
| Figure 6N | Mean harvest level at travel (standardized to 18μl patch) | 14 mice, 3 sessions/mouse | Mixed-effects model (REML) with Geisser-Greenhouse correction, and Tukey's multiple comparisons post hoc | $F_{(1.425,17.81)}$=8.181, P=0.0058, Geisser-Greenhouse's epsilon 0.7124 Tukey's multiple comparisons: Sal 24 vs LPS 24 q=5.956, df=12, adjusted P=0.0032 Sal 24 vs LPS 48 q=1.525, df=13, P=0.5432 LPS 24 vs LPS 48 q=3.197, df=12, adjusted P=0.1008 |
| Figure 6O | Difference in mean harvest level at travel (standardized to 18μl patch) from saline session | 14 mice, 3 sessions/mouse | Mixed-effects model (REML) with Geisser-Greenhouse correction, and Tukey's multiple comparisons post hoc | $F_{(1.452,27.59)}$=9.626, P=0.0018, Geisser-Greenhouse's epsilon 0.7261 Tukey's multiple comparisons: Sal 24 vs LPS 24 q=6.377, df=12, adjusted P=0.0019 Sal 24 vs LPS 48 q=1.525, df=13, P=0.5432 LPS 24 vs LPS 48 q=3.481, df=12, adjusted P=0.0716 |
| Figure 6P | Change in mean harvest level at travel (standardized to 18μl patch) from saline session (%) | 14 mice, 3 sessions/mouse | Mixed-effects model (REML) with Geisser-Greenhouse correction, and Tukey's multiple comparisons post hoc | $F_{(1.548,29.41)}$=9.701, P=0.0013, Geisser-Greenhouse's epsilon 0.7739 Tukey's multiple comparisons: Sal 24 vs LPS 24 q=7.030, df=12, adjusted P=0.0009 Sal 24 vs LPS 48 q=1.573, df=13, P=0.5236 LPS 24 vs LPS 48 q=3.567, df=12, adjusted P=0.0645 |
| Figure 6V | Change in BW (%) from 0-24h post LPS injection | 14 mice (14 XY pairs) | Simple linear regression (Change in BW (%) from 0-24h post LPS-injection as a function of proportion of food left uneaten (%) from 24h post LPS-injection) | Y=-0.1278*X+3.309, $R^2$=0.8779, slope significantly non-zero, $F_{(1,12)}$=86.29, P<0.0001 |
| Figure 6V | Change in BW (%) from 0-24h post LPS injection | 14 mice (14 XY pairs) | Pearson's correlation (Change in BW (%) from 0-24h post LPS-injection as a function of proportion of food | r=-0.9370, 95% CI: -0.9802 to -0.8082, $R^2$=0.8779, P<0.0001 |

| | | | | |
|---|---|---|---|---|
| | | | left uneaten (%) from 24h post LPS-injection) | |
| Figure 6W | Change in mean harvest level at travel (standardized to 18μl patch) from saline session (%) | 12 mice (12 XY pairs) | Simple linear regression (Change in mean harvest level at travel (standardized to 18μl patch) from saline session (%) as a function of proportion of food left uneaten (%) from 24h post LPS-injection) | Y=0.5396*X+10.02, $R^2$=0.5396, slope significantly non-zero, $F_{(1,10)}$=12.28, P=0.0057 |
| Figure 6W | Change in mean harvest level at travel (standardized to 18μl patch) from saline session (%) | 12 mice (12 XY pairs) | Pearson's correlation (Change in mean harvest level at travel (standardized to 18μl patch) from saline session (%) as a function of proportion of food left uneaten (%) from 24h post LPS-injection) | r=0.7424, 95% CI: 0.2936 to 0.9230, $R^2$=0.5512, P=0.0057 |
| Figure 6X | Change in trial number from saline session (%) | 12 mice (12 XY pairs) | Simple linear regression (Change in trial number from saline session (%) as a function of change in BW (%) from 0-24h post LPS-injection) | Y=3.843*X-13.51, $R^2$=0.5564, slope significantly non-zero, $F_{(1,10)}$=12.54, P=0.0053 |
| Figure 6X | Change in trial number from saline session (%) | 12 mice (12 XY pairs) | Pearson's correlation (Change in trial number from saline session (%) as a function of change in BW (%) from 0-24h post LPS-injection) | r=0.7459, 95% CI: 0.3007 to 0.9242, $R^2$=0.5564, P=0.0053 |
| Figure 6Y | Change in mean harvest level at travel (standardized to 18μl patch) from saline session (%) | 13 mice (13 XY pairs) | Simple linear regression (Change in mean harvest level at travel (standardized to 18μl patch) from saline session (%) as a function of change in BW (%) from 0-24h post LPS-injection) | Y=-3.249*X+27.63, $R^2$=0.3697, slope significantly non-zero, $F_{(1,10)}$=5.865, P=0.0360 |
| Figure 6Y | Change in mean harvest level at travel (standardized to 18μl patch) from saline session (%) | 13 mice (13 XY pairs) | Pearson's correlation (Change in mean harvest level at travel (standardized to 18μl patch) from saline session (%) | r=-0.0680, 95% CI: -0.8762 to -0.05238, $R^2$=0.3697, P=0.0360 |

| | | | |
|---|---|---|---|
| | | | as a function of change in BW (%) from 0-24h post LPS-injection) |

**Supplementary Table 7.** Supplementary Figure 1 Statistics

| Figure Panel | Response Variable | n | Statistical analysis | Result |
|---|---|---|---|---|
| Figure S1G | Change in BW (g) from pre- to post-session | 20 mice, 15 sessions/mouse (300 XY pairs) | Simple linear regression (Change in BW (g) as a function of total session reward (kcal)) | Y=1.267*X-0.6253, $R^2$=0.6253; slope significantly non-zero, $F_{(1,298)}$=522.1, P<0.0001 |
| Figure S1G | Change in BW (g) from pre- to post-session | 20 mice, 15 sessions/mouse (300 XY pairs) | Pearson's correlation (Change in BW (g) as a function of total session reward (kcal)) | r=0.7979, 95% CI: 0.7526 to 0.8356, $R^2$=0.6253, P<0.0001 |

**Supplementary Table 8.** Supplementary Figure 5 Statistics

| Figure Panel | Response Variable | n | Statistical analysis | Result |
|---|---|---|---|---|
| Figure S5D | BW change from -1h to 0h pre-behaviour | 35 mice, 2 sessions/condition per mouse | Two-tailed paired t-test | t=13.52, df=69, P<0.0001 |
| Figure S5F | BW (g) at 0hr pre-behaviour | 35 mice, 2 sessions/condition per mouse | Two-tailed paired t-test | t=12.72, df=69, P<0.0001 |
| Figure S5H | Mean active patch touch response frequency per session (0-30 min behaviour) | 35 mice, 2 sessions/condition per mouse | Two-tailed paired t-test | t=4.666, df=69, P<0.0001 |
| Figure S5J | Mean active patch discrimination (%) per session (0-30 min behaviour) | 35 mice, 2 sessions/condition per mouse | Two-tailed paired t-test | t=1.943, df=69, P=0.0561 |
| Figure S5L | Mean harvest frequency per session (0-30 min behaviour) | 35 mice, 2 sessions/condition per mouse | Two-tailed paired t-test | t=3.870, df=34, P=0.0005 |
| Figure S5N | Mean travel touch frequency per session (0-30 min behaviour) | 35 mice, 2 sessions/condition per mouse | Two-tailed paired t-test | t=0.4160, df=69, P=0.6787 |
| Figure S5P | Mean travel frequency per session (0-30 min behaviour) | 35 mice, 2 sessions/condition per mouse | Two-tailed paired t-test | t=0.3743, df=69, P=0.7093 |
| Figure S5R | Mean total trials per session (0-30 min behaviour) | 35 mice, 2 sessions/condition per mouse | Two-tailed paired t-test | t=4.592, df=69, P<0.0001 |

**Supplementary Table 9.** Supplementary Figure 7 Statistics

| Figure Panel | Response Variable | n | Statistical analysis | Result |
|---|---|---|---|---|
| Figure S7A | Mean harvest level at travel (0-5 min behaviour) | 35 mice, 2 sessions/condition per mouse (139 XY pairs) | Simple linear regression (mean harvest level at travel as a function of BW (% free feeding) at -1h pre-behaviour) | Y=0.1379*X–4.517, $R^2$=0.004015, slope not significantly non-zero, $F_{(1,137)}$=0.5522, P=0.4587 |
| Figure S7A | Mean harvest level at travel (0-5 min behaviour) | 35 mice, 2 sessions/condition per mouse (139 XY pairs) | Pearson's correlation (mean harvest level at travel as a function of BW (% free feeding) at -1h pre-behaviour) | r=0.06336, 95% CI: -0.1042 to 0.2275, $R^2$=0.004015, P=0.4587 |
| Figure S7B | Mean harvest level at travel (0-5 min behaviour) | 35 mice, 2 sessions/condition per mouse (139 XY pairs) | Simple linear regression (mean harvest level at travel as a function of BW (g) at -1h pre-behaviour) | Y=0.09487*X+4.939, $R^2$=0.007822, slope not significantly non-zero, $F_{(1,137)}$=1.080, P=0.3005 |
| Figure S7B | Mean harvest level at travel (0-5 min behaviour) | 35 mice, 2 sessions/condition per mouse (139 XY pairs) | Pearson's correlation (mean harvest level at travel as a function of BW (% free feeding) at -1h pre-behaviour) | r=0.08844, 95% CI: -0.07923 to 0.2512, $R^2$=0.007822, P=0.3005 |
| Figure S7C | Mean harvest level at travel (0-5 min behaviour) | 35 mice, 2 sessions/condition per mouse (139 XY pairs) | Simple linear regression (mean harvest level at travel as a function of BW (g) at 0h pre-behaviour) | Y=0.04760*X+6.298, $R^2$=0.001736, slope not significantly non-zero, $F_{(1,137)}$=0.2383, P=0.6262 |
| Figure S7C | Mean harvest level at travel (0-5 min behaviour) | 35 mice, 2 sessions/condition per mouse (139 XY pairs) | Pearson's correlation (mean harvest level at travel as a function of BW (g) at 0h pre-behaviour) | r=0.04167, 95% CI: -0.1257 to 0.2067, $R^2$=0.001736, P=0.6262 |
| Figure S7D | Mean harvest level at travel (0-5 min behaviour) | 35 mice, 2 sessions/condition per mouse (139 XY pairs) | Simple linear regression (mean harvest level at travel as a function of BW change from -1h to 0h pre-behaviour) | Y=-0.8232*X+7.986, $R^2$=0.03785, slope significantly non-zero, $F_{(1,137)}$=5.389, P=0.0217 |
| Figure S7D | Mean harvest level at travel (0-5 min behaviour) | 35 mice, 2 sessions/condition per mouse (139 XY pairs) | Pearson's correlation (mean harvest level at travel as a function of BW change from -1h to 0h pre-behaviour) | r=-0.1945, 95% CI: -0.3497 to -0.02898, $R^2$=0.03785, P=0.0217 |
| Figure S7E | Mean harvest level at travel (0-30 min behaviour) | 35 mice, 2 sessions/condition per mouse (140 XY pairs) | Simple linear regression (mean harvest level at travel as a function of BW (% of free- | Y=0.07563*X+2.216, $R^2$=0.003581, slope not significantly non-zero, $F_{(1,138)}$=0.4959, P=0.4825 |

| | | | | |
|---|---|---|---|---|
| | | | feeding) at -1h pre-behaviour) | |
| Figure S7E | Mean harvest level at travel (0-30 min behaviour) | 35 mice, 2 sessions/condition per mouse (140 XY pairs) | Pearson's correlation (mean harvest level at travel as a function of BW (% of free-feeding) at -1h pre-behaviour) | r=0.05984, 95% CI: -0.1071 to 0.2235, $R^2$=0.003581, P=0.4825 |
| Figure S7F | Mean harvest level at travel (0-30 min behaviour) | 35 mice, 2 sessions/condition per mouse (140 XY pairs) | Simple linear regression (mean harvest level at travel as a function of BW (g) at -1h pre-behaviour) | Y=0.1478*X+4.605, $R^2$=0.05757, slope significantly non-zero, $F_{(1,138)}$=8.430, P=0.0043 |
| Figure S7F | Mean harvest level at travel (0-30 min behaviour) | 35 mice, 2 sessions/condition per mouse (140 XY pairs) | Pearson's correlation (mean harvest level at travel as a function of BW (g) at -1h pre-behaviour) | r=0.2399, 95% CI: 0.07710 to 0.3903, $R^2$=0.05757, P=0.0043 |
| Figure S7G | Mean harvest level at travel (0-30 min behaviour) | 35 mice, 2 sessions/condition per mouse (140 XY pairs) | Simple linear regression (mean harvest level at travel as a function of BW (g) at 0hr pre-behaviour) | Y=0.1427*X+4.706, $R^2$=0.04706, slope significantly non-zero, $F_{(1,138)}$=6.815, P=0.0100 |
| Figure S7G | Mean harvest level at travel (0-30 min behaviour) | 35 mice, 2 sessions/condition per mouse (140 XY pairs) | Pearson's correlation (mean harvest level at travel as a function of BW (g) at 0hr pre-behaviour) | r=0.2169, 95% CI: 0.5294 to 0.3695, $R^2$=0.04706, P=0.0100 |
| Figure S7H | Mean harvest level at travel (0-30 min behaviour) | 35 mice, 2 sessions/condition per mouse (140 XY pairs) | Simple linear regression (mean harvest level at travel as a function of change in BW (g) from -1h to 0h pre-behaviour) | Y=-0.3580*X+9.037, $R^2$=0.02135, slope not significantly non-zero, $F_{(1,138)}$=3.011, P=0.0850 |
| Figure S7H | Mean harvest level at travel (0-30 min behaviour) | 35 mice, 2 sessions/condition per mouse (140 XY pairs) | Pearson's correlation (mean harvest level at travel as a function of change in BW (g) from -1h to 0h pre-behaviour) | r=-0.1461, 95% CI: -0.3046 to 0.02028, $R^2$=0.02135, P=0.0850 |

**Supplementary Table 10.** Supplementary Figure 8 Statistics

| Figure Panel | Response Variable | n | Statistical analysis | Result |
|---|---|---|---|---|
| Figure S8D | Mean latency to travel choice (s) | 19 mice, 18 sessions/mouse | Repeated measures one-way ANOVA with Geisser-Greenhouse correction; Tukey's post-hoc multiple comparisons | $F_{(1.576,28.37)}$=75.85, P<0.0001, Geisser-Greenhouse's epsilon 0.7882 Tukey's post hoc comparisons: 6µl vs 12µl q=6.473, df=18, adjusted P=0.0007; |

| | | | | |
|---|---|---|---|---|
| | | | | 6µl vs 18µl q=13.85, df=18, adjusted P<0.0001; 12µl vs 18µl q=13.31, df=18, adjusted P<0.0001 |
| Figure S8E | Mean travel choice probability (%) at Harvest Level 1 | 19 mice, 18 sessions/mouse | Repeated measures one-way ANOVA with Geisser-Greenhouse correction | $F_{(1.714,30.85)}$=3.067, P=0.0681, Geisser-Greenhouse's epsilon 0.8571 |
| Figure S8F | Mean reward/effort ratio | 19 mice, 18 sessions/mouse | Repeated measures one-way ANOVA with Geisser-Greenhouse correction; Tukey's post-hoc multiple comparisons | $F_{(1.399,25.18)}$=2926, P<0.0001, Geisser-Greenhouse's epsilon 0.6994 Tukey's post hoc comparisons: 6µl vs 12µl q=74.35, df=18, adjusted P<0.0001; 6µl vs 18µl q=84.09, df=18, adjusted P<0.0001; 12µl vs 18µl q=58.15, df=18, adjusted P<0.0001 |
| Figure S8G | Mean effort/reward ratio | 19 mice, 18 sessions/mouse | Repeated measures one-way ANOVA with Geisser-Greenhouse correction; Tukey's post-hoc multiple comparisons | $F_{(1.039,18.70)}$=974.9, P<0.0001, Geisser-Greenhouse's epsilon 0.5195 Tukey's post hoc comparisons: 6µl vs 12µl q=40.44, df=18, adjusted P<0.0001; 6µl vs 18µl q=46.68, df=18, adjusted P<0.0001; 12µl vs 18µl q=54.86, df=18, adjusted P<0.0001 |
| Figure S8I | Mean harvest level at travel (standardized to 18µl patch) | 19 mice, 18 sessions/mouse | Repeated measures one-way ANOVA with Geisser-Greenhouse correction; Tukey's post-hoc multiple comparisons | $F_{(1.275,22.96)}$=340.7, P<0.0001, Geisser-Greenhouse's epsilon 0.6377, Tukey's post hoc comparisons: 6µl vs 12µl q=29.88, df=18, adjusted P<0.0001; 6µl vs 18µl q=28.36, df=18, adjusted P<0.0001; 12µl vs 18µl q=18.94, df=18, adjusted P<0.0001 |
| Figure S8J | Mean harvest level at travel (6µl patch) | 19 mice, 18 sessions/mouse | Repeated measures one-way ANOVA with Geisser-Greenhouse correction | $F_{(1.953,33.15)}$=0.9557, P=0.3924, Geisser-Greenhouse's epsilon 0.9764 |

| | | | | |
|---|---|---|---|---|
| Figure S8K | Mean harvest level at travel (12μl patch) | 19 mice, 18 sessions/mouse | Repeated measures one-way ANOVA with Geisser-Greenhouse correction | $F_{(1.430,25.74)}$=1.263, P=0.2882, Geisser-Greenhouse's epsilon 0.7151 |
| Figure S8L | Mean harvest level at travel (18μl patch) | 19 mice, 18 sessions/mouse | Repeated measures one-way ANOVA with Geisser-Greenhouse correction | $F_{(1.705,30.69)}$=2.947, P=0.0749, Geisser-Greenhouse's epsilon 0.8526 |

**Supplementary Table 11.** Supplementary Figure 15 Statistics

| Figure Panel | Response Variable | n | Statistical analysis | Result |
|---|---|---|---|---|
| Figure S15B | Mean food intake (kcal) | 12 mice (12 pairs) | Two-tailed paired t-test | t=9.853, df=11, p<0.0001 |
| Figure S15D | Mean locomotor activity (beam breaks) | 12 mice (12 pairs) | Two-tailed paired t-test | t=0.7777, df=11, p=0.4531 |
| Figure S15F | Mean energy expenditure (kcal/h) | 12 mice (12 pairs) | Two-tailed paired t-test | t=4.398, df=11, p=0.0011 |
| Figure S15F | Mean RER | 12 mice (12 pairs) | Two-tailed paired t-test | t=6.859, df=11, p<0.0001 |
| Figure S15J | Mean FAO (kcal/h) | 12 mice (12 pairs) | Two-tailed paired t-test | t=3.616, df=11, p=0.0041 |

**Supplementary Table 12.** Supplementary Figure 16 Statistics

| Figure Panel | Response Variable | n | Statistical analysis | Result |
|---|---|---|---|---|
| Figure S16B | Food intake (kcal) | 12 mice (12 pairs) | Two-tailed paired t-test | t=5.359, df=11, p=0.0002 |
| Figure S16D | Meal frequency (0-6 h from injection) | 12 mice (12 pairs) | Two-tailed paired t-test | t=5.227, df=11, p=0.0003 |
| Figure S16E | Meal frequency 48-54h or 72-78h | 12 mice (12 pairs) | Two-tailed paired t-test | t=2.288, df=11, p=0.0429 |
| Figure S16G | Mean meal size (kcal) at 0-6h from injection | 12 mice (12 pairs) | Two-tailed paired t-test | t=0.4470, df=11, p=0.6635 |
| Figure S16H | Mean meal size (kcal) at 48-54h or 72-78h | 12 mice (12 pairs) | Two-tailed paired t-test | t=1.273, df=11, p=0.2294 |
| Figure S16J | Mean meal duration (s) at 0-6h from injection | 12 mice (12 pairs) | Two-tailed paired t-test | t=0.07886, df=11, p=0.9386 |
| Figure S16K | Mean meal duration (s) at 48-54h or 72-78h | 12 mice (12 pairs) | Two-tailed paired t-test | t=0.07886, df=11, p=0.9386 |

**Supplementary Table 13.** Supplementary Figure 17 Statistics

| Figure Panel | Response Variable | n | Statistical analysis | Result |
|---|---|---|---|---|
| Figure S17A | Food intake (kcal) at 0-8h from injection | 12 mice (12 pairs) | Two-tailed paired t-test | t=2.157, df=11, p=0.0540 |
| Figure S17B | Food intake (kcal) at 16-24h from injection | 12 mice (12 pairs) | Two-tailed paired t-test | t=6.184, df=11, p<0.0001 |

| | | | | |
|---|---|---|---|---|
| Figure S17C | Locomotor activity (beam breaks) at 0-8h from injection | 12 mice (12 pairs) | Two-tailed paired t-test | t=3.832, df=11, p=0.0028 |
| Figure S17D | Locomotor activity (beam breaks) at 16-24h from injection | 12 mice (12 pairs) | Two-tailed paired t-test | t=3.749, df=11, p=0.0032 |
| Figure S17E | Energy expenditure (kcal/8h) at 0-8h from injection | 12 mice (12 pairs) | Two-tailed paired t-test | t=3.758, df=11, p=0.0032 |
| Figure S17F | Energy expenditure (kcal/8h) at 16-24h from injection | 12 mice (12 pairs) | Two-tailed paired t-test | t=1.302, df=11, p=0.2196 |
| Figure S17G | Mean RER at 0-8h from injection | 12 mice (12 pairs) | Two-tailed paired t-test | t=1.095, df=11, p=0.2968 |
| Figure S17H | Mean RER at 16-24h from injection | 12 mice (12 pairs) | Two-tailed paired t-test | t=3.810, df=11, p=0.0029 |
| Figure S17I | Fatty acid oxidation (kcal/8h) at 0-8h from injection | 12 mice (12 pairs) | Two-tailed paired t-test | t=0.3489, df=11, p=0.7338 |
| Figure S17J | Fatty acid oxidation (kcal/8h) at 16-24h from injection | 12 mice (12 pairs) | Two-tailed paired t-test | t=3.392, df=11, p=0.0060 |
| Figure S17K | Food intake (kcal) at 0-24h from injection | 12 mice (12 pairs) | Two-tailed paired t-test | t=0.7595, df=11, p=0.4635 |
| Figure S17L | Energy expenditure (kcal/24h) at 0-24h from injection | 12 mice (12 pairs) | Two-tailed paired t-test | t=0.1867, df=11, p=0.8553 |
| Figure S17M | Energy in/energy out ratio at 0-8h from injection | 12 mice (12 pairs) | Two-tailed paired t-test | t=1.975, df=11, p=0.0739 |
| Figure S17N | Energy in/energy out ratio at 0-24h from injection | 12 mice (12 pairs) | Two-tailed paired t-test | t=0.6420, df=11, p=0.5340 |

**Supplementary Table 14.** Supplementary Figure 18 Statistics

| **Figure Panel** | **Response Variable** | **n** | **Statistical analysis** | **Result** |
|---|---|---|---|---|
| Figure S18A | Absolute BW (g) at injection time | n=7 saline, n=6 LPS | Unpaired two-tailed t-test | t=0.3249, df=11, P=0.7513 |
| Figure S18B | Relative BW (% free-feeding) at injection time | n=7 saline, n=6 LPS | Unpaired two-tailed t-test | t=0.5586, df=11, P=0.5876 |
| Figure S18C | Relative BW (% free-feeding 3 hr post-injection | n=7 saline, n=6 LPS | Unpaired two-tailed t-test | t=1.619, df=11, P=0.1338 |
| Figure S18D | Cumulative food intake (kcal) 3 hr post-injection | n=7 saline, n=6 LPS | Unpaired two-tailed t-test | t=3.844, df=11, P=0.0027 |
| Figure S18E | ARC *Tnf* mRNA expression | n=7 saline, n=6 LPS | Unpaired two-tailed t-test | t=3.534, df=133, P=0.0047 |
| Figure S18F | ARC *Ccl2* mRNA expression | n=7 saline, n=6 LPS | Unpaired two-tailed t-test | t=2.986, df=11, P=0.0124 |
| Figure S18G | ARC *Il1b* mRNA expression | n=7 saline, n=6 LPS | Unpaired two-tailed t-test | T4.387, df=11, P=0.0011 |
| Figure S18H | ARC *Ifng* mRNA expression | n=7 saline, n=6 LPS | Unpaired two-tailed t-test | t=1.148, df=11, P=0.2753 |
| Figure S18I | VTA *Tnf* mRNA expression | n=7 saline, n=6 LPS | Unpaired two-tailed t-test | t=13.78, df=11, P<0.0001 |

| | | | | |
|---|---|---|---|---|
| Figure S18J | VTA *Ccl2* mRNA expression | n=7 saline, n=6 LPS | Unpaired two-tailed t-test | t=14.31, df=11, P<0.0001 |
| Figure S18K | VTA *Il1b* mRNA expression | n=7 saline, n=6 LPS | Unpaired two-tailed t-test | t=11.50, df=11, P<0.0001 |
| Figure S18L | VTA *Ifng* mRNA expression | n=7 saline, n=6 LPS | Unpaired two-tailed t-test | t=0.3647, df=11, P=0.7222 |
| Figure S18M | NAc *Tnf* mRNA expression | n=7 saline, n=6 LPS | Unpaired two-tailed t-test | t=6.356, df=11, P<0.0001 |
| Figure S18N | NAc *Ccl2* mRNA expression | n=7 saline, n=6 LPS | Unpaired two-tailed t-test | t=5.730, df=11, P=0.0001 |
| Figure S18O | NAc *Il1b* mRNA expression | n=7 saline, n=6 LPS | Unpaired two-tailed t-test | t=4.770, df=11, P=0.0006 |
| Figure S18P | NAc *Ifng* mRNA expression | n=7 saline, n=6 LPS | Unpaired two-tailed t-test | t=0.1481, df=11, P=0.8849 |

#### **Supplementary Table 15.** Supplementary Figure 19 Statistics

| **Figure Panel** | **Response Variable** | **n** | **Statistical analysis** | **Result** |
|---|---|---|---|---|
| Figure S19A | Body weight (g) at injection time | 14 mice (14 pairs) | Two-tailed paired t-test | t=0.4490, df=13, P=0.6608 |
| Figure S19B | Body weight (% of free feeding) at injection time | 14 mice (14 pairs) | Two-tailed paired t-test | t=0.4698, df=13, P=0.6462 |
| Figure S19G | Feed efficiency at 0-24h or 25-28h post LPS injection | 14 mice, 2 time periods/mouse | Wilcoxon matched-pairs signed rank test | Sum of signed ranks (W) 67.00, Sum of positive ranks 86.00, Sum of negative ranks -19.00, P=0.0353 |
| Figure S19H | Ratio of forced travel frequency/total trials | 14 mice, 2 sessions/mouse | Two-tailed paired t-test | t=1.849, df=13, P=0.0873 |
| Figure S19I | Proportion of forced travels of total travels | 14 mice, 3 sessions/mouse | Mixed-effects model (REML) with Geisser-Greenhouse correction | $F_{(1.072,13.40)}$=3.518, P=0.0806, Geisser-Greenhouse's epsilon 0.5360 |
| Figure S19J | Active patch discrimination (%) | 14 mice, 3 sessions/mouse | Friedman's test | Friedman statistic 1.000, approximate P=0.6065 |
| Figure S19K | Mean harvest level at travel (24h post saline) | 14 mice, 3 patch types/mouse | Friedman's test with Dunn's multiple comparisons post hoc | Friedman statistic 25.78, approximate P<0.0001 Dunn's multiple comparisons: 6µl vs 12µl Rank sum diff. -11.00, adjusted P=0.1129, 6µl vs 18µl Rank sum diff.-26.50, adjusted P<0.0001, 12µl vs 18µl Rank sum diff.-15.50, adjusted P=0.0102 |
| Figure S19L | Mean harvest level at travel (24h post LPS) | 14 mice, 3 patch types/mouse | Mixed-effects model (REML) with Geisser-Greenhouse correction, and Tukey's multiple | $F_{(1.205,12.65)}$=7.230, P=0.0154, Geisser-Greenhouse's epsilon 0.6025 Tukey's multiple comparisons: |

| | | | | |
|---|---|---|---|---|
| | | | comparisons post hoc | 6μl vs 12μl q=6.193, df=11, adjusted P=0.0029<br>6μl vs 18μl q=3.688, df=10, adjusted P=0.0622<br>12μl vs 18μl q=1.693, df=10, adjusted P=0.4812 |
| Figure S19M | Mean harvest level at travel (48h post LPS) | 14 mice, 3 patch types/mouse | Friedman's test with Dunn's multiple comparisons post hoc | Friedman statistic 24.57, approximate P<0.0001<br>Dunn's multiple comparisons:<br>6μl vs 12μl Rank sum diff. -16.00, adjusted P<0.0001<br>6μl vs 18μl Rank sum diff. -26.00, adjusted P<0.0001<br>12μl vs 18μl Rank sum diff. -10.00, adjusted P=0.1763 |
| Figure S19Q | Mean time to travel choice (24h post saline) | 14 mice, 3 patch types/mouse | Repeated measures one-way ANOVA with Geisser-Greenhouse correction, and Tukey's multiple comparisons post hoc | $F_{(1.565,20.35)}$=110.6, P<0.0001<br>Tukey's multiple comparisons:<br>6μl vs 12μl q=1.362, df=13, adjusted P=0.6119,<br>6μl vs 18μl q=6.812, df=13, adjusted P=0.0009,<br>12μl vs 18μl q=9.359, df=13, adjusted P<0.0001 |
| Figure S19R | Mean time to travel choice (24h post LPS) | 14 mice, 3 patch types/mouse | Mixed-effects model (REML) with Geisser-Greenhouse correction, and Tukey's multiple comparisons post hoc | $F_{(1.293,11.64)}$=4.597, P=0.0463, Geisser-Greenhouse's epsilon 0.6466<br>Tukey's multiple comparisons:<br>6μl vs 12μl q=4.397, df=8, adjusted P=0.0199,<br>6μl vs 18μl q=3.907, df=8, adjusted P=0.0575,<br>12μl vs 18μl q=2.717, df=10, adjusted P=0.1830 |
| Figure S19S | Mean time to travel choice (48h post LPS) | 14 mice, 3 patch types/mouse | Mixed-effects model (REML) with Geisser-Greenhouse correction, and Tukey's multiple comparisons post hoc | $F_{(1.101,12.11)}$=16.73, P=0.0012, Geisser-Greenhouse's epsilon 0.5506<br>Tukey's multiple comparisons:<br>6μl vs 12μl q=5.823, df=10, adjusted P=0.0054, |

| | | | | |
|---|---|---|---|---|
| | | | | 6μl vs 18μl q=6.263, df=12, adjusted P=0.0022,<br>12μl vs 18μl q=7.830, df=10, adjusted P=0.0007 |
| Figure S19T | Change in mean harvest level at travel (standardized to 18μl patch) from saline session (%) | 12 mice (12 XY pairs) | Simple linear regression (Change in mean harvest level at travel (standardized to 18μl patch) from saline session (%) as a function of change in number of trials (%) from saline session) | Y=-0.5897*X+26.54, $R^2$=0.3092, slope not significantly non-zero, $F_{(1,10)}$=4.476, P=0.0605 |
| Figure S19T | Change in mean harvest level at travel (standardized to 18μl patch) from saline session (%) | 12 mice (12 XY pairs) | Pearson's correlation (Change in mean harvest level at travel (standardized to 18μl patch) from saline session (%) as a function of change in number of trials (%) from saline session) | r=-0.5561, 95% CI: -0.8566 to 0.02619, $R^2$=0.3092, P=0.0605 |
| Figure S19U | Cumulative travels (18μl patch) | 14 mice | Kolmogorov-Smirnov test | Sal 24 vs LPS 24 Kolmogorov-Smirnov D 0.2414, approximate P=0.3668,<br>LPS 24 vs LPS 48 Kolmogorov-Smirnov D 0.2414, approximate P=0.3668,<br>Sal 24 vs LPS 48 Kolmogorov-Smirnov D 0.1034, approximate P=0.9978 |
| Figure S19V | Cumulative travels (12μl patch) | 14 mice | Kolmogorov-Smirnov test | Sal 24 vs LPS 24 Kolmogorov-Smirnov D 0.3913, approximate P=0.0591,<br>LPS 24 vs LPS 48 Kolmogorov-Smirnov D 0.3478, approximate P=0.1237,<br>Sal 24 vs LPS 48 Kolmogorov-Smirnov D 0.3913, approximate P=0.0591 |
| Figure S19W | Cumulative travels (6μl patch) | 14 mice | Kolmogorov-Smirnov test | Sal 24 vs LPS 24 Kolmogorov-Smirnov D 0.5294, |

| | | | | |
|---|---|---|---|---|
| | | | | approximate P=0.0171<br>LPS 24 vs LPS 48 Kolmogorov-Smirnov D 0.4706, approximate P=0.0463<br>Sal 24 vs LPS 48 Kolmogorov-Smirnov D 0.2941, approximate P=0.4540 |
